# Predicting the Environmental Suitability and Population at Risk of Podoconiosis in Africa

**DOI:** 10.1101/2020.03.04.977827

**Authors:** Kebede Deribe, Hope Simpson, Rachel L. Pullan, Mbonigaba Jean Bosco, Samuel Wanji, Nicole Davis Weaver, Christopher J. L. Murray, Melanie J. Newport, Simon I. Hay, Gail Davey, Jorge Cano

**Affiliations:** Department of Global Heath and Infection, Brighton and Sussex Medical School, Falmer, Brighton, BN1 9PX, UK; School of Public Health, College of Health Sciences, Addis Ababa University, Addis Ababa, Ethiopia; Department of Disease Control, London School of Hygiene & Tropical Medicine, London, UK; Malaria and Other Parasitic Disease Division, Rwanda Biomedical Center–Ministry of Health, Kigali, Rwanda; Parasites and Vector Biology Research Unit (PAVBRU), Department of Microbiology and Parasitology, University of Buea, Buea, Cameroon; Research Foundation in Tropical Diseases and the Environment (REFOTDE), Buea, Cameroon; Institute for Health Metrics and Evaluation, University of Washington, Seattle, WA, USA; Department of Health Metrics Sciences, University of Washington, Seattle, WA, USA

## Abstract

**Background:** Podoconiosis is a type of tropical lymphedema that causes massive swelling of the lower limbs. The disease is associated with both economic insecurity, due to long-term morbidity-related loss of productivity, and intense social stigma. The geographical distribution and burden of podoconiosis in Africa is uncertain.

**Methods:** We applied statistical modelling to the most comprehensive database compiled to date to predict the environmental suitability of podoconiosis in the African continent. By combining climate and environmental data and overlaying population figures, we predicted the suitability and human population at risk.

**Results:** In Africa, environmental suitability for podoconiosis was predicted in 29 countries. By 2020, the total population in areas suitable for podoconiosis was estimated at 114.5 million people, (95% confidence interval: 109.4-123.9) with 16.9 million in areas suitable for both lymphatic filariasis and podoconiosis. Of the total 5,712 implementation units defined by WHO in Africa, 1,655 (29.0%) were found to be environmentally suitable for podoconiosis. The majority of IUs with high environmental suitability are located in Angola (80 IUs), Cameroon (170 IUs), the DRC (244 IUs), Ethiopia (495 IUs), Kenya (217 IUs), Uganda (116 IUs) and Tanzania (112 IUs). Of the 1,655 environmental suitable IUs, 960 (58.0%) require more detailed community-level mapping

**Conclusions:** Our estimates provide key evidence of the population at risk and geographical extent of podoconiosis in Africa, which will help decision-makers to better plan more integrated intervention programmes.

## Introduction

Podoconiosis is a neglected tropical disease (NTDs) caused by exposure to red clay soil^1^ in genetically susceptible people who do not use footwear. It is one of the leading causes of lymphoedema in Africa^2,3^. The disease is a disabling NTD which causes progressive bilateral swelling of the legs, significantly reducing quality of life^4^ and productivity^5^. People affected by podoconiosis also suffer comorbid mental distress and depression^6^, stigma and discrimination^7,8^. The current intervention includes prevention through consistent footwear usage starting from an early age, regular foot hygiene and covering housing floors^9^. For those with the disease, the WHO recommends simple lymphoedema management consisting of foot hygiene, foot care, wound care, compression, exercises and elevation, treatment of acute attacks and use of shoes and socks to reduce further exposure to the irritant soil^10,11^. Podoconiosis is one of the diseases with potential for elimination, due to the fact that it is non-infectious and there are proven prevention and treatment interventions^9^. Historical evidence shows that it has already been eliminated from northern African countries including Algeria, Morocco and Tunisia^3^.

Although precise estimates are lacking, evidence suggests that globally, four million people are disabled by podoconiosis in 27 countries thought to be endemic^12,13^. In Africa, podoconiosis has been reported in 18 countries, mostly in the East and West African regions. Nonetheless, most of the data originate from the 1970s^12^, such that the current situation is unclear. Podoconiosis is often misdiagnosed and confused with other causes of lymphoedema^14^. In a study conducted in an endemic zone in Ethiopia, around half of health workers interviewed inaccurately thought podoconiosis to be an infectious disease transmitted by mosquitoes, and of those who had treated podoconiosis, 71% prescribed diethylcarbamazine assuming that lymphatic filariasis (LF) was the cause^14^. Misdiagnosis of podoconiosis may not only lead to underestimation of the burden of podoconiosis, but also to underestimation of the success of LF elimination programmes, by overestimating the morbidity burden due to LF ^15,16^.

Three countries (Cameroon, Ethiopia and Rwanda) have recently mapped the distribution of podoconiosis through nationwide mapping activities ^17–20^, revealing widespread distribution of podoconiosis, constituting a considerable burden. These studies have also provided new epidemiological insights. Firstly, they identified climate and topographical factors, such as precipitation, elevation and land surface temperature; environmental factors such as enhanced vegetation index and proximity to waterways; and soil-related factors such as soil composition (i.e. clay, silt and clay content) and soil pH, as potential drivers of disease distribution. Other factors such as night-light emissivity and distance to stable night-lights, considered indirect indicators of poverty ^21–23^, were also found to predict the occurrence of podoconiosis ^17–20^. Although these factors cannot be considered *per se* to cause the disease, they have proven to be very informative in predicting the occurrence of podoconiosis by characterising its environmental niche.

Podoconiosis national programmes require a detailed understanding of the geographical distribution of the disease so that all endemic areas can be targeted. Previous attempts to map podoconiosis include a detailed literature review^12^ and mapping at national^17,19,20^ and subnational levels^24–26^. However, there are no maps outlining the potential distribution of podoconiosis at higher geographical levels. To develop national plans and investment case programmes, planners and policy makers require robust data on the geographical distribution and population at risk, and areas to be targeted for more detailed community-level mapping. Podoconiosis and lymphatic filariasis (LF) share a principal clinical manifestation—lymphoedema. In endemic areas these two diseases are confused. Understanding the graphical overlaps of these two diseases has clinical and programmatic benefits^9^. One of the pillars of LF elimination is morbidity management and disability prevention—i.e., providing access to basic care for LF-related diseases to every affected person in endemic areas^27^. Delineating the overlap of LF and podoconiosis will help join hands to provide access to care which will help both programmes. In addition, the existing LF programme platform and transmission assessments surveys can be used to integrate mapping of podoconiosis and case searches.

In the current investigation, we aimed to predict the environmental limits of podoconiosis across the African continent, using available data from field surveys and other sources, and empirical evidence of its distribution^17–19,28^. Building on methods used to map the distribution of podoconiosis in known endemic countries, we developed a model to outline the environmental suitability across Africa. The present work aimed to: i) map the geographical distribution of podoconiosis in Africa, ii) estimate the population at risk of podoconiosis, iii) determine the number of WHO implementation units (IUs) where mapping is required, and iv) delineate overlapping risk of podoconiosis and LF. This work was conducted within the context of the Global Atlas of Podoconiosis, which seeks to develop a global map of podoconiosis and estimate its burden^29^.

## Methods

### Data sources

We used data recently collected in country-wide and local surveys conducted in Ethiopia, Cameroon and Rwanda ^17–20^. These datasets were supplemented with data compiled under the Global Atlas of Podoconiosis, which comprises a number of relevant studies providing evidence of podoconiosis occurrence^12^. The podoconiosis database was first created in 2018, with published literature, case reports and series, and last updated 10 May, 2019. Briefly, we searched for studies that reported the epidemiology of podoconiosis. We searched databases including MEDLINE and SCOPUS from inception to 10 May, 2019 for all relevant studies that examined podoconiosis occurrence, prevalence, incidence and case reports. We used the following search terms; “podoconiosis” OR “mossy foot” OR “non-filarial elephantiasis”. No time or language limits were applied. We hand-searched the reference lists of all recovered documents for additional references. Abstracts of all reports were read and full papers retrieved for those appearing to fulfil selection criteria. Publications were eligible for inclusion in the occurrence if they reported geographical locations with evidence of podoconiosis. We contacted authors of studies to further obtain data and geographic coordinates. We searched the grey literature by seeking reports not published in peer-reviewed journals through contacting experts, a search of conference abstracts and reviewing Price’s monograph^3^. Further details on the development of this data repository for podoconiosis are provided elsewhere ^12,28^.

### Geo-positioning of survey data

Geographic information extracted from publications included geographical coordinates, when these were available, location names and administrative location (i.e. district, region, or state). When coordinates were not provided, we resorted to geocoders and online electronic gazetteers to identify the most accurate geographic coordinates. Briefly, automated georeferencing was implemented in R software through the Google Maps API Engine and Opencage Geocoder, using the *ggmap* and *opencage* packages respectively^30,31^. Locations that could not be georeferenced by these methods were manually searched in Google using different gazetteers: Bing Maps^32^, GeoNet Names Server^33^, Fuzzy gazetteers^34^ and the Open Street Map project^35^. All geographic coordinates were standardised to decimal degrees in order to be displayed in the WGS84 geographic coordinate system.

### Description of covariates

Data on extrinsic determinants of podoconiosis were assembled from remotely sensed environmental datasets (S1 Appendix). Geographic coordinates of each community were used to extract from gridded maps estimates of: 1) annual precipitation, 2) land surface temperature, 3) distance to water surfaces (water bodies and streams), 4) elevation, 5) enhanced vegetation index (EVI), 6) soil composition (i.e. silt and clay soil fraction) and 7) soil pH. These climate, topographical, environmental and soil-related factors have been found to be associated with the occurrence of podoconiosis in several studies ^18–20,36^.

Gridded continuous maps, namely raster datasets, of averaged EVI and land surface temperature (LST) for the period 2000-2017 were generated from MODIS satellite image data downloaded from the Earth Explorer NASA site (https://earthexplorer.usgs.gov/). The MOD13Q1 product from MODIS library, which is updated every 16 days at 250m spatial resolution, includes vegetation indices such as Normalized Difference Vegetation Index (NDVI) and EVI^37^. Day and night LST data were obtained from MOD11A2 products, and have a spatial and temporal resolution of 1km and 8 days respectively^38^.

Information on rainfall was extracted from a synoptic gridded map of annual precipitation calculated from monthly total precipitation gridded datasets obtained from WorldClim v2.0 Global Climate Database^39^. This database provides a set of global climate layers obtained by interpolation of precipitation data for the period 1970–2000 collected in weather stations distributed across the world ^40,41^. From the Consortium for Spatial Information (CGIAR-CSI), we obtained a raster dataset of elevation at 1 square-km^42^. This elevation layer resulted from processing and resampling the gridded digital elevation models (DEM) derived from the original 30-arcsecond DEM produced by the Shuttle Radar Topography Mission (SRTM).

Soil data including silt and clay fraction and soil-pH of the top soil, were obtained from the ISRIC-World Soil Information project^43^. This project provides gridded maps of soil composition at 250m resolution worldwide. We also generated continuous surfaces of straight-line distance (Euclidean distance) in km to the nearest water body and permanent rivers based on data obtained from OpenStreetMap project. Waterbodies and waterways were downloaded from the OpenStreetMap project (OSM)^44^ through the platform *Geofabrik*^45^.

Input grids were resampled to a common spatial resolution of 5km × 5km using a bilinear interpolation and clipped to the geographic extent of a map displaying Africa limits, and eventually aligned to it. Raster manipulation and processing was undertaken using *raster* package in R v3.3.2 and final map layouts created with ArcGIS 10.5 software (ESRI Inc., Redlands CA, USA).

### Data analysis and modelling

An ensemble of distribution models was generated based on the reported occurrence of podoconiosis in the surveyed communities and the environmental factors. Communities were reclassified as endemic (1) or non-endemic (0) for podoconiosis based on records of confirmed podoconiosis cases.

We used six algorithms available within the BIOMOD framework^46^ to obtain ensembles of predicted distributions: generalised linear models (GLM), generalised additive models (GAM), generalised boosted regression models (GBM), artificial neural networks (ANN), multiple adaptive regression splines (MARS), and random forest (RF). These models were run using the parameters set by default in the *biomod2* R package ^46^, except for the GBM models. For the latter, the learning rate (*lr*) and tree complexity (*tc*) parameters in boosted regression models, were set to enable up to five interactions, and reducing the speed (*lr*: 0.005) to allow the model to converge without over-fitting. This tuning was undertaken using the *gbm* package in R v3.5.3.

The species distribution models we developed all require training with presence and absence points in order to predict environmental suitability. True absence is difficult to determine in surveys, as the absence of cases identified does not rule out the existence of cases when only a subsample of total population is surveyed. The country-wide mapping surveys conducted in Cameroon, Rwanda and Ethiopia, compiled and used in the current study, targeted the total population of the communities randomly selected across all the administrative regions (i.e. regions/states, districts/zones). Health workers were trained to identify all cases of lymphoedema of lower limbs within the communities, and podoconiosis was subsequently confirmed by expert clinical diagnostic teams ^18,19,47,48^. Therefore, we considered communities not reporting cases as ‘*true*’ absences and give them the same weight (1) as the presence records in the analysis. To represent absence locations in areas where exhaustive surveys had not been undertaken, we generated random data points, termed ‘*pseudo-absence*’ records, in countries where the presence of the disease is considered unlikely. This approach has been used in similar modelling studies in the past, and has proven to be helpful when presence and absence data are only available for restricted geographical areas ^49,50^. To define the area of assumed unsuitability, we used a recently published map displaying the strength of evidence for presence and absence of podoconiosis across Africa ^28^. A sample of *pseudo-absence* points, equal to the number of compiled presence and absence records, was randomly generated from countries with an evidence score below 0 (‘*weak to complete evidence of absence*’) *Pseudo-absence* records were weighted based on a rescaled version (0 to 1 scale) of the evidence score assigned for this study to each country^28^.

Models were calibrated using an 80% random sample of the initial data and evaluated against the remaining 20% data using the area under the curve (AUC) of the receiver operation characteristic (ROC), the true skill statistic (TSS)^51^ and the proportion correctly classified (PCC). Projections were performed 50 times per algorithm, each time selecting a different 80% random sample while verifying model accuracy against the remaining 20%. The evaluation statistics (AUC and TSS) were used to select the models to be assembled based on the matching between predictions and observations. Here, models with AUC < 0.8 or TSS values < 0.7 were disregarded when assembling the final model.

The final ensemble model was obtained by estimating the weighted mean of probabilities across the selected models per grid cell. This algorithm returns the predicted mean weighted by the selected evaluation method scores, in our case the AUC statistic score. The range of uncertainties was also calculated by estimating the confidence intervals around the mean of probabilities across the ensemble per grid cell.

The base map of the global administrative areas was downloaded from the Natural Earth (https://www.naturalearthdata.com/)^52^. All maps were produced using ArcGIS Desktop v10.5 (Environmental Systems Research Institute Inc., Redlands CA, USA).

### Cross-validation and setting occurrence map

We used spatial block cross-validation to validate the environmental model. Briefly, the whole area of Africa was split in-to five regular squares of 2,000km by 2,000km (“folds”), which were tagged from 1 to 5 in a random order. Single models were trained using all the data (occurrences, absences and pseudo-absences) within 4 out of the 5 folds (training data) and the data in the remaining fold (test data) were used to assess the quality of the predictive performance of the trained models (S1 Appendix). Under this approach to model evaluation, test datasets are more spatially independent from training datasets compared to those in a random internal cross-validation appraoch, reducing the risk of over-optimistic evaluation due to shared ecological characteristics of nearby locations. An ensemble model was constructed using the same specifications as the model run with regular cross-validation. Both models were compared using the Pearson’s correlation coeficient.

The resulting predictive map quantifies the environmental suitability for podoconiosis. In order to convert this continuous metric into a binary map outlining the distribution limits (i.e. ecological limits), a threshold value of suitability was determined above which podoconiosis occurrence was assumed to be possible. This cut-off represents the best trade-off between sensitivity, specificity and accuracy (fraction correctly classified). In addition, partial dependence functions were performed separately for two modelling approaches, GBM and RF, to visualise dependencies between the probability of podoconiosis occurrence and covariates. The partial dependence function shows the marginal effect of each covariate on the response after averaging the effects of all other covariates.

### Estimation of population at-risk and potential geographical overlapping with LF

We estimated the number of individuals at risk by 2020 by overlaying the binary raster dataset displaying the potential suitability for podoconiosis occurrence on a gridded population density map^53,54^ and calculating the population in cells considered to be within the limits of podoconiosis occurrence. The 95% credible interval of the population at risk was calculated based on the uncertainty in environmental suitability, by summarising the 50 predictions by mean and 95% credible intervals. Potential for overlapping with lymphatic filariasis (LF) was analysed by overlaying a gridded map of predicted LF occurrence published by Cano et al. (2014)^55^ and our map of predicted occurrence for podoconiosis. Both maps were combined and the resulting map was used to estimate the population living in areas environmentally suitable for both podoconiosis and LF applying the above mentioned procedure.

## Results

### Characteristic of the data

The data search strategy identified 2,870 spatially unique data points from 42 studies undertaken between January, 1934, and May, 2019, in 12 countries in Africa. Of the 2,870 spatially referenced data points 1,311 (45.7%) represented podoconiosis presence. Almost all (99%) of the collected occurrences are distributed across eastern (Ethiopia, Kenya, Rwanda and Uganda) and western Africa (Cameroon, Equatorial Guinea and Sao Tomé and Principe) (Figure 1).

**Fig 1.**
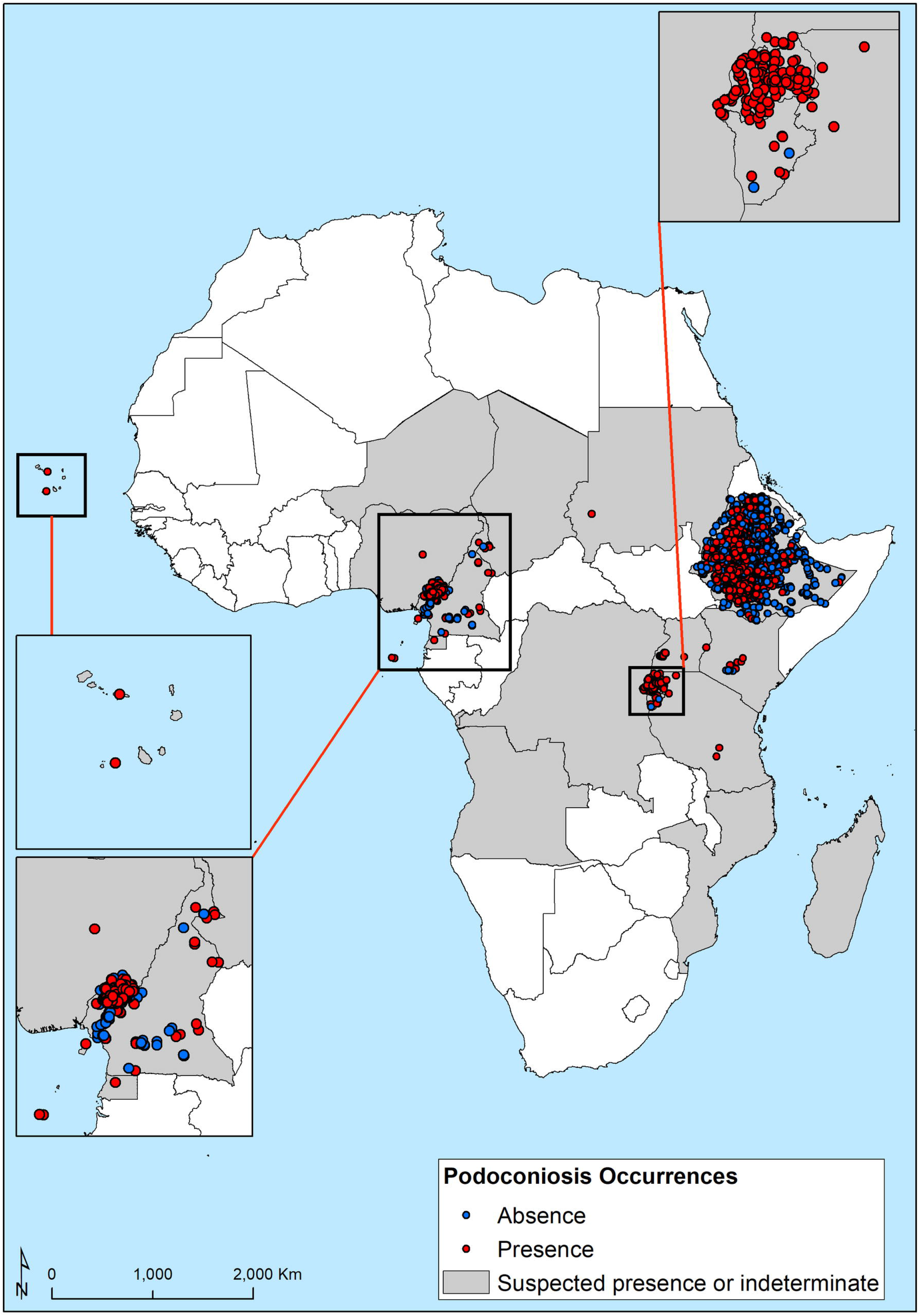
Geographical distribution of presence and absence surveys used in the predictive map of occurrence. Red dots correspond to survey locations where podoconiosis has been reported (n=1,311).

### Environmental limits of podoconiosis suitability in Africa

Elevation, annual precipitation, soil pH-H_2_0 and clay fraction in the topsoil were the major contributors to the GBM and RF ensembles (S1 Appendix). The marginal effect plots for these covariates indicates that the probability of podoconiosis occurrence increases with annual precipitation, reaching a peak between 1,000mm and 1,800mm, and then declining. For elevation, we observed an increased risk in areas located over 1,000masl (‘*meters above sea level*’) and sustained up to 2,500masl. Regarding soil composition, acid soils (pH H_2_0<7) and soils rich in clay and silt are more likely to increase human risk of triggering podoconiosis (S1 Appendix).

We used Global Burden of Disease (GBD) regions^56^ to present the results (S1 Appendix). In the Central Africa Region, three countries (Angola, Central African Republic and the Democratic Republic of Congo) were predicted to be suitable for the occurrence of podoconiosis. In the Democratic Republic of Congo, the suitability was predicted in the eastern mountainous part of the country, bordering with countries historically considered endemic (Uganda, Rwanda and Burundi). The mainland region of Equatorial Guinea was found to be unsuitable for the occurrence of podoconiosis, while Bioko Island -which has historically reported cases of podoconiosis, was still found to be suitable for the disease. Podoconiosis was predicted in Sao Tomé Island but not in Principe (Figure 2). For more detail, country maps displaying the predicted environmental suitability for podoconiosis and predicted overlapping with LF are provided in S2 Appendix.

**Fig 2.**
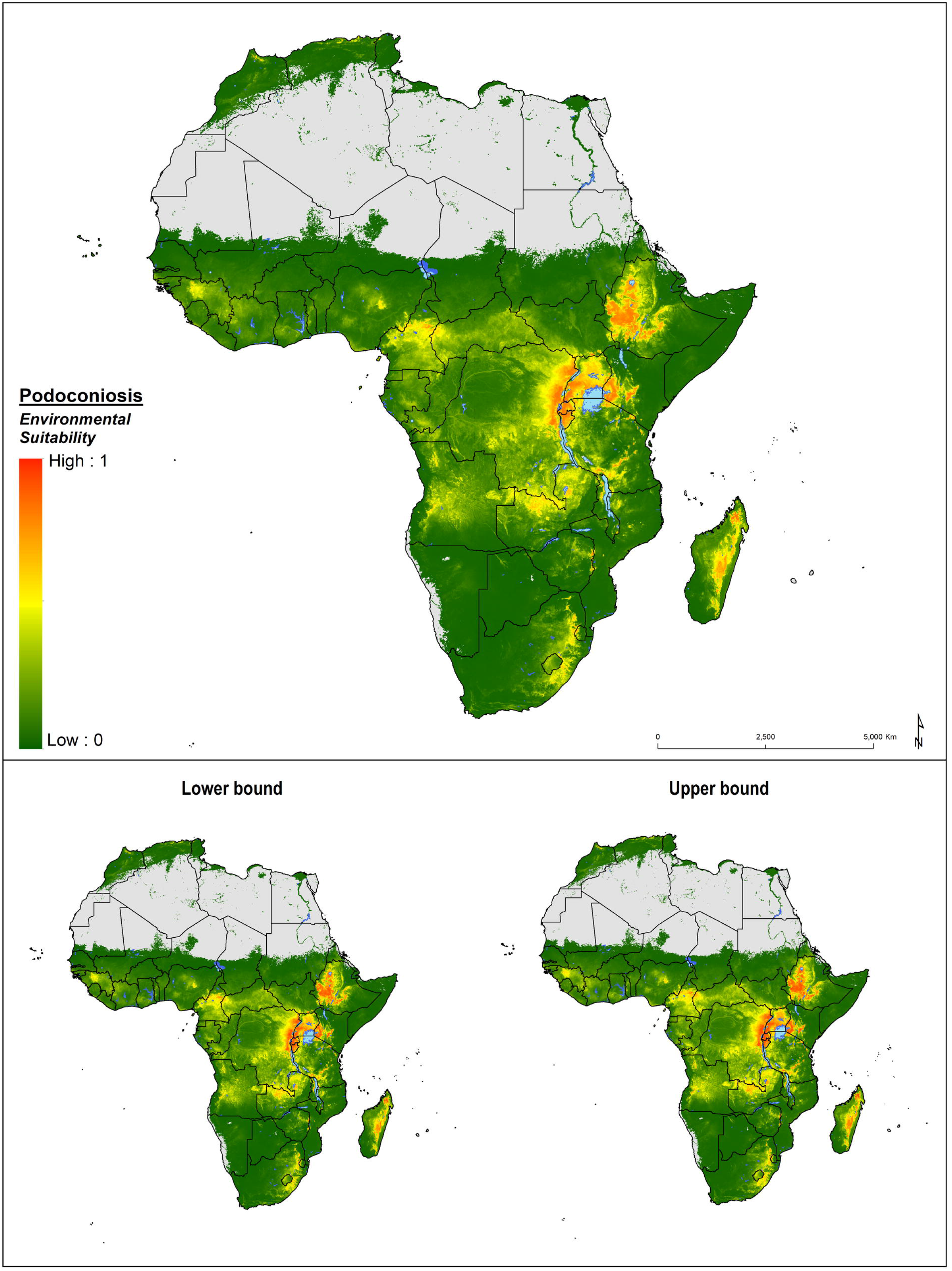
Environmental suitability for the occurrence of podoconiosis across Africa. Probability of occurrence is provided in a 0 to 1 scale; orange colour indicates highest probability of occurrence (1) and green lowest probability (0). Lower (2.5%) and upper (97.5%) bound of presence limits were obtained from fitting an enssemble of 120 BRT submodels to predict sets of different risk maps.

The highest suitability for podoconiosis was predicted in East Africa Region; most parts of Ethiopia, Uganda, Rwanda and Burundi were predicted to be suitable for podoconiosis occurrence. Northern and western parts of Tanzania and the western part of Kenya were also found to be suitable. Parts of the highlands of Madagascar and northern Zambia (which borders with the Democratic Republic of Congo) were also predicted to be suitable. Patchy areas in northern Mozambique, mountainous area in Niassa province, and Malawi, Nyika Plateau and Viphya mountains at the north and around Zomba Mountain and Mount Mulanje at the south, were found to be suitable for podoconiosis occurrence (S1 Appendix & S2 Appendix).

Most of the Southern Africa Region was not predicted to be suitable for the occurrence of podoconiosis. Pockets of environmental suitability were predicted in South Africa, along the mountain ranges of Blouberg (Blue mountains) and the Drakensberg (Dragons mountains); the Swaziland border with the Drakensberg mountains in South Africa; and the eastern highlands of Zimbabwe (S1 Appendix & S2 Appendix).

In the West Africa Region, our model predicted suitable areas in Cameroon, Cape Verde, Guinea, Nigeria and Sao Tomé Island. In Cameroon, at-risk areas were delineated in most of the central and northwest parts, whereas in Nigeria, suitability appeared to be localized in the central and eastern parts of the country, the latter bordering with Cameroon. Interestingly, the risk map reflects environmental conditions suitable for podoconiosis in countries of northern Africa where the disease is considered eliminated (Morocco and Tunisia) (Figure 2 and S2 Appendix).

Validation statistics indicated an excellent predictive performance of all the algorithms (S1 Appendix). However, RF and GBM outperformed the other models with AUC scores of 0.88 (95%CI: 0.88 – 0.9) and 0.89 (95%CI: 0.89 – 0.9), respectively. An environmental suitability threshold of 0.389 provided the best discrimination between presence and absence records; sensitivity 80.1%, specificity 85.5% and AUC score of 0.921, and this threshold value was used to classify the environmental suitability map into a binary map of the environmental limits of occurrence (Figure 3). The final ensemble model showed a high correlation (Pearson’s correlation coefficient: 0.9902) with a model constructed using regular spatial blocks for cross-validation, indicating a high stability of the model irrespective of the method chosen for cross-validation and model selection (S1 Appendix).

**Figure 3.**
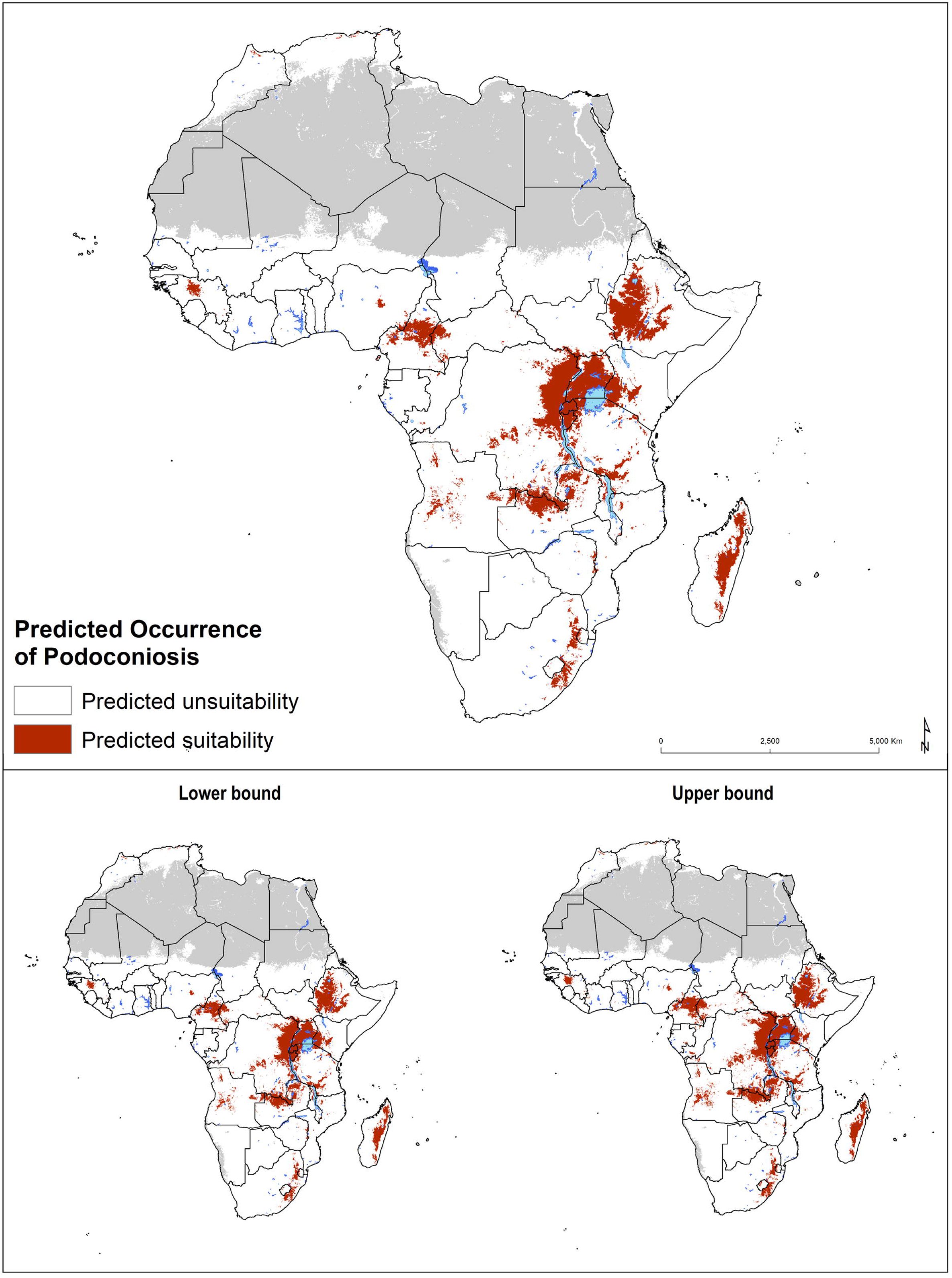
**Predicted occurrence of podoconiosis in Africa with the lower (2.5%) and upper (97.5%) bounds of the occurrence limits** based on the optimal threshold of environmental suitability (0.389).

### Estimating population at risk and geographical overlapping with LF

The population living in areas environmentally suitable for podoconiosis was estimated to be over 114.5 million (95% UI: 109.4 – 123.9) (Table 1). The largest proportion of the population at risk was found in east Africa (81.7%), followed by central Africa (10.9%). Northern, Southern and West African countries only accounted for 7.4% of the total population at-risk. Geographical overlap between podoconiosis and LF was mainly predicted in Eastern, Western and central Africa regions (Figure 4). Large areas of potential co-occurrence of podoconiosis and LF were predicted in Uganda, north of Lake Victoria, in Guinea, and in more restricted areas in Cameroon, Ethiopia and Madagascar. The population living in areas environmentally suitable for podoconiosis and LF was estimated to be 16.9 million, 14.7% of the population at risk for podoconiosis (Table 2). Eastern African countries accounted for 82.0% of the total population living in areas of potential geographic overlap for podoconiosis and LF.

**Fig 4.**
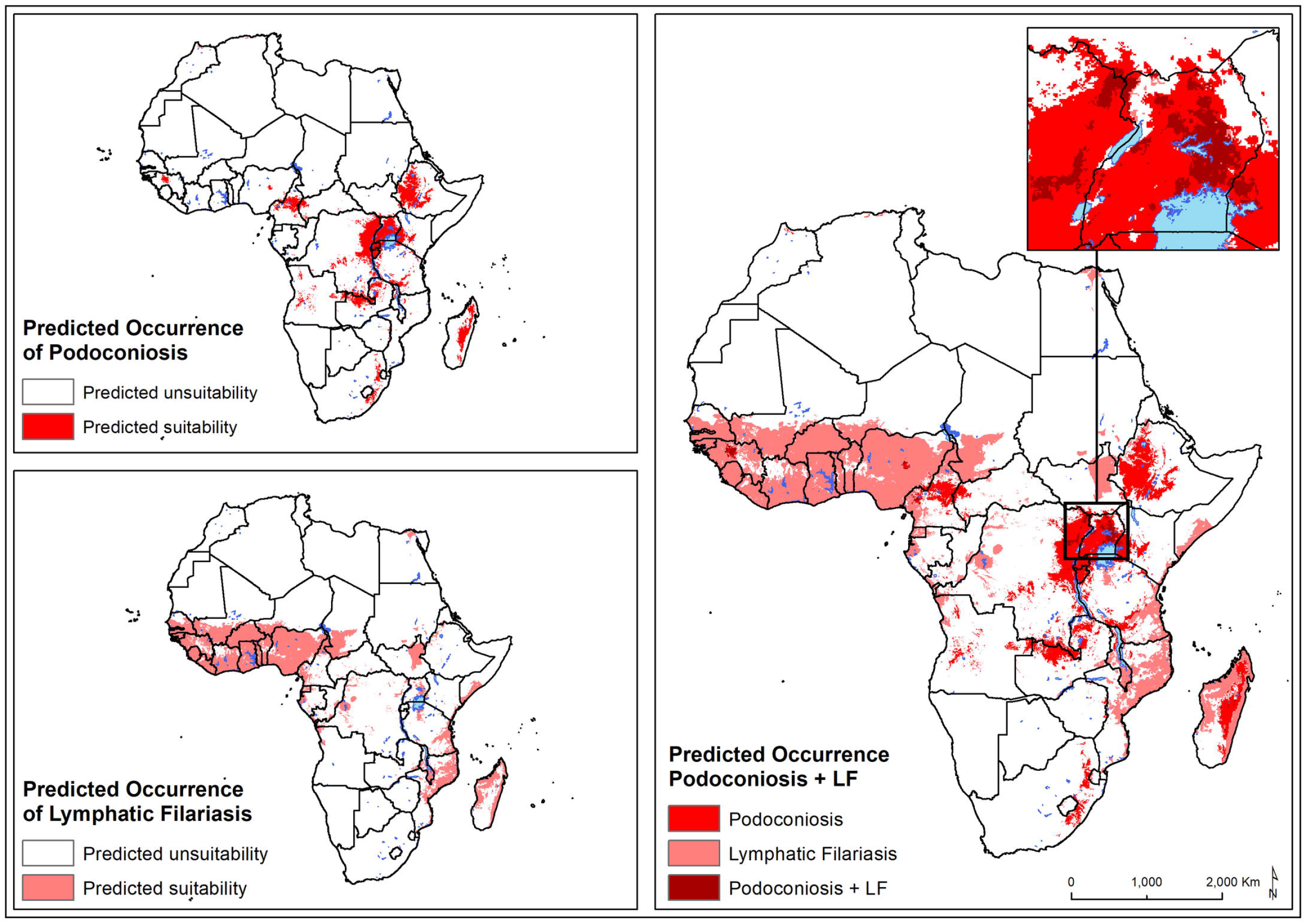
Predicted occurrence of podoconiosis and lymphatic filariasis in Africa. Environmental suitability for lymphatic filariasis according to the predictive map published by Cano et al. (2014).

**Table 1.** Estimates of population living in areas environmentally suitable for podoconiosis in Africa. Estimates were obtained from a gridded map of population density for 2020 (www.worldpop.org).

| Africa Region | Population at risk | 95% UI |
| --- | --- | --- |
| Central | 12,529,248 | (11,592,258 - 14,413,843) |
| East | 93,507,570 | (90,335,551- 98,217,585) |
| North | 360,038 | (300,847- 480,627) |
| South | 3,065,753 | (2,631,172- 4,458,757) |
| West | 5,014,597 | (4,517,515- 6,373,689) |
| **Total** | **114,477,207** | **(109,377,342- 123,944,501)** |

**Table 2.** Estimates of population living in areas with overlapping risk of podoconiosis and lymphatic filariasis in Africa.

| Africa Region | Only podoconiosis at risk population 2020 | Only LF at risk population | At risk population in LF and podoconiosis overlapping areas |
| --- | --- | --- | --- |
| Central | 11,660,060 | 15,758,872 | 869,188 |
| East | 79,675,440 | 36,548,506 | 13,832,130 |
| North | 360,038 | 42,376,960 | 0 |
| South | 3,063,100 | 36,686 | 2,653 |
| West | 2,856,736 | 182,910,848 | 2,157,861 |
| **Total** | **97,615,374** | **277,631,872** | **16,861,833** |

### Risk of podoconiosis in Implementation Units (IUs) in Africa

Based on the optimal threshold of environmental suitability (0.389), of the total 5,712 WHO implementation units (typically second administrative-level units, such as districts) in Africa 1,655 (29.0%) IUs were found to be environmentally suitable for podoconiosis. The majority of IUs with high environmental suitability are located in Angola (80 IUs), Cameroon (170 IUs), the DRC (244 IUs), Ethiopia (495 IUs), Kenya (217 IUs), Uganda (116 IUs) and Tanzania (112 IUs). Of the 1,655 environmental suitable IUs, 960 (58.0%) require more detailed community-level mapping (Figure 5).

**Fig 5.**
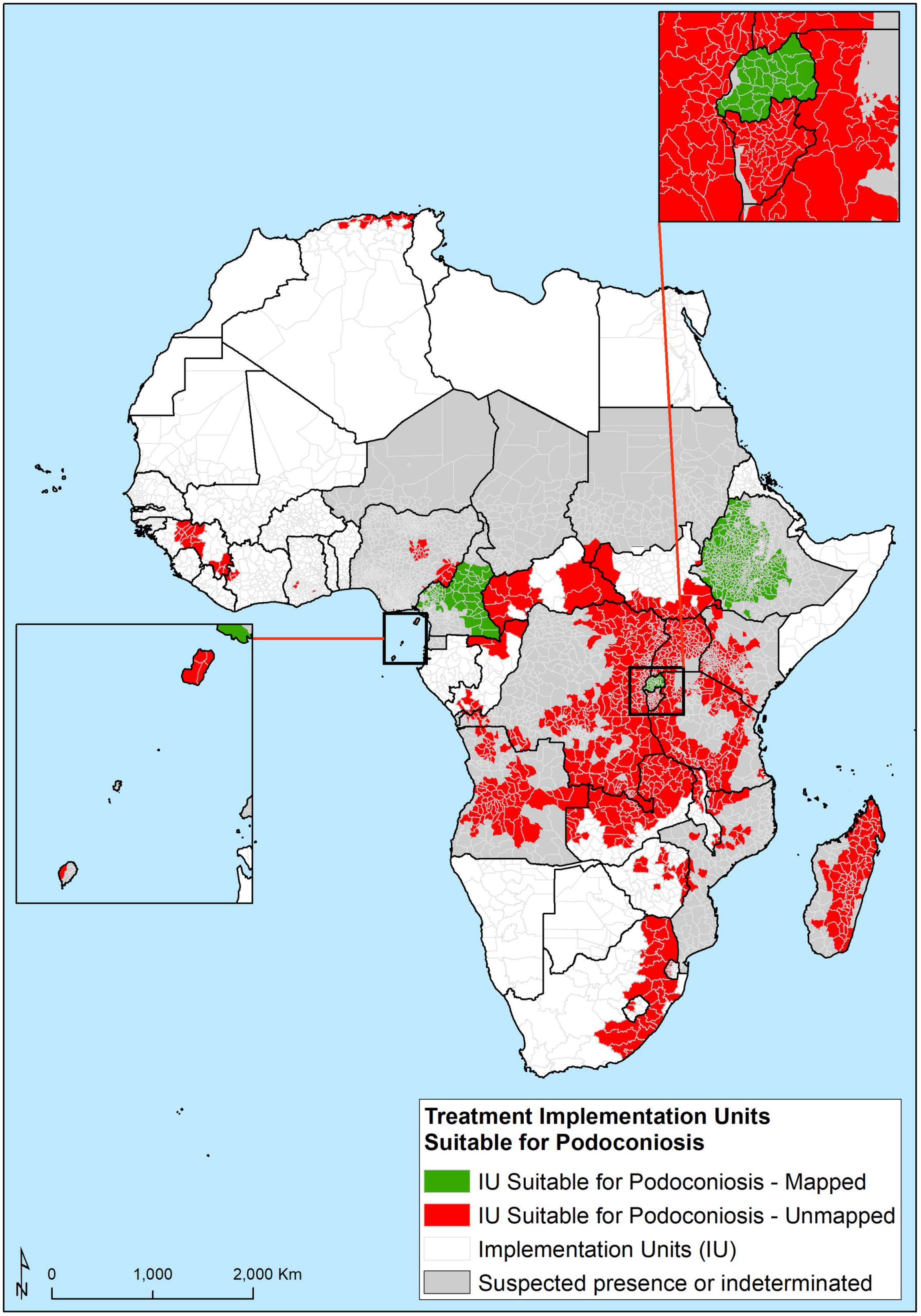
IU-level suitability classification based on WHO geographic units for implementation of control interventions. Red identifies suitable IUs which are not mapped and green identifies suitable IUs mapped for podoconiosis.

## Discussion

Our model predicted environmental suitability for podoconiosis in many areas in Africa, with the strongest prediction largely in East, West and Central Africa Regions. In agreement with previous work, our model showed podoconiosis environmental suitability is largely influenced by annual precipitation, elevation, clay fraction and pH of the soil ^17–19,36,47^. The novelty of this study was in translating suitability to the IU level: aggregated risk at the IU level aids in mapping podoconiosis. These findings are not only useful in understanding the risk, population at risk, and the environmental limits of podoconiosis across Africa, but also are valuable for targeted mapping of podoconiosis in the remaining endemic countries.

Our model predicted suitability in 29 countries in Africa, nonetheless we have identified 18 countries with evidence of presence of podoconiosis in Africa previously. This is a clear reflection that suitability only reflects the availability of ecological suitability for the occurrence of podoconiosis, not necessarily the presence of the disease currently. One good example is that historical evidence indicates the elimination of podoconiosis from northern African countries (e.g. Algeria, Morocco and Tunisia)^3^, nonetheless our model predicted suitable areas in parts of these countries. Although the disease has been eliminated, the ecological characteristics which are suitable for the occurrence of podoconiosis, such as the soil characteristics, remain the same. Hence our results should be put into context. For this reason, we have restricted the list of IUs needing mapping to those 18 countries with evidence of podoconiosis presence.

Mapping podoconiosis is an important prerequisite for the initiation of data-driven national control programmes^57^. Surveying all districts in suspected endemic countries will be resource intensive and will take time. Here we provide model-based guidance to identify environmentally suitable IUs eligible for mapping activity. By focusing efforts in suitable countries and ensuring suitable IUs within those countries, resources can be targeted and used efficiently. Our prediction can also guide surveillance efforts, targeting areas predicted to be suitable. Here we are not ruling out the need to conduct mapping, our results may provide evidence in prioritizing mapping surveys and surveillance efforts to high yield areas.

Environmental suitability of podoconiosis is predicted in ten countries with no historical report of the disease (including Côte d’Ivoire, Congo, Ghana, Guinea, Malawi, Lesotho, Swaziland, South Africa, Zambia and Zimbabwe)^3,12^. As podoconiosis is a gene-environment-behaviour interaction^13,58^, it is possible that it has not been reported because other elements (e.g. genetic susceptibility or lack of footwear) are not present in these countries. Several have strong health systems, making it unlikely that cases are missed without being detected ^59–61^. We encourage these countries to increase their index of suspicion for podoconiosis when lymphoedema cases are diagnosed and treated. We have identified areas suitable for both podoconiosis and LF. This has important implications in terms of mapping and implementation of interventions. Areas where the LF programme has already advanced, the existing platform and programme activities such as the transmission assessment surveys (TAS) can be used to integrate podoconiosis mapping. Once the mapping is completed integrated morbidity movement services can be provided to all cases, as has been found to be feasible in Ethiopia^62^.

Delineating the potentially suitable areas for the occurrence of podoconiosis in Africa has several benefits. First, it provides a framework for targeted mapping of podoconiosis in known and suspected endemic countries by limiting this to areas suitable for the occurrence of the disease. Second, it provides a basis to stratify surveillance activities by identifying areas with different level of risk. Finally, it provides an important input for estimating the burden of podoconiosis at the continent level.

### Limitations

There are some limitations of our analysis. First, our predictive mean environmental suitability is not a measure of disease prevalence or incidence. Our model characterised the similarity between locations based on covariates included with locations with confirmed cases of podoconiosis. This neither measures the magnitude of the problem nor confirms the presence of podoconiosis, it only measures the environmental suitability. Nonetheless, podoconiosis is a gene-environment disease^13,58^. To develop the disease, people must be exposed to the environment and must also be genetically susceptible. This means we might overestimate the population at risk by assuming all individuals living in areas of environmental suitability are at risk of podoconiosis. Nonetheless, our approach is stringent in identifying potential endemic areas for mapping, rather than missing potential endemic areas. Second, only a few of the surveys were conducted at the national level. Most of the surveys were conducted in potentially endemic areas, where investigators observed high prevalence of lymphoedema cases. Therefore, our modelling might be characterising areas with high prevalence rather than the whole spectrum of prevalence. Third, we were not able to account for important covariates, including footwear use, which is not available for analysis.

## Conclusions

In conclusion, our work has highlighted considerable environmental suitability of podoconiosis in Africa with a significant population at risk in the continent. We have identified areas suitable for podoconiosis, which needs priority both for mapping and intervention. Importantly, we have determined the number of IUs which require mapping based on the optimal cut-off point for suitability. Targeted mapping and intensified surveillance are required in areas where suitability is determined. We have also identified areas where podoconiosis and LF risk overlaps, and where it is possible to use the existing platforms to identify podoconiosis cases in LF-endemic districts, and establish a mechanism whereby podoconiosis cases can access morbidity management services in co-endemic areas. Thus, while our results and maps guide surveillance and survey activities to better define the local distribution and burden of podoconiosis in suitable areas, we encourage countries to collect primary data in pursuit of accelerating the progress towards a world without podoconiosis.

## Supporting information

**S1 Appendix:** covariates, marginal effects.

**S2 Appendix:** country level risk prediction.

