## Supplementary Appendix 1 for "Predicting the Environmental Suitability and Population at Risk of Podoconiosis in Africa"

### Supplementary file 1

**Figure 1S** Variables used to model environmental suitability of podoconiosis across Africa. A) Annual Precipitation (mm), B) Land surface temperature averaged for the period 2000-2017 (ºC), C) Distance to water bodies (km), and D) Elevation (metres above sea level, masl).

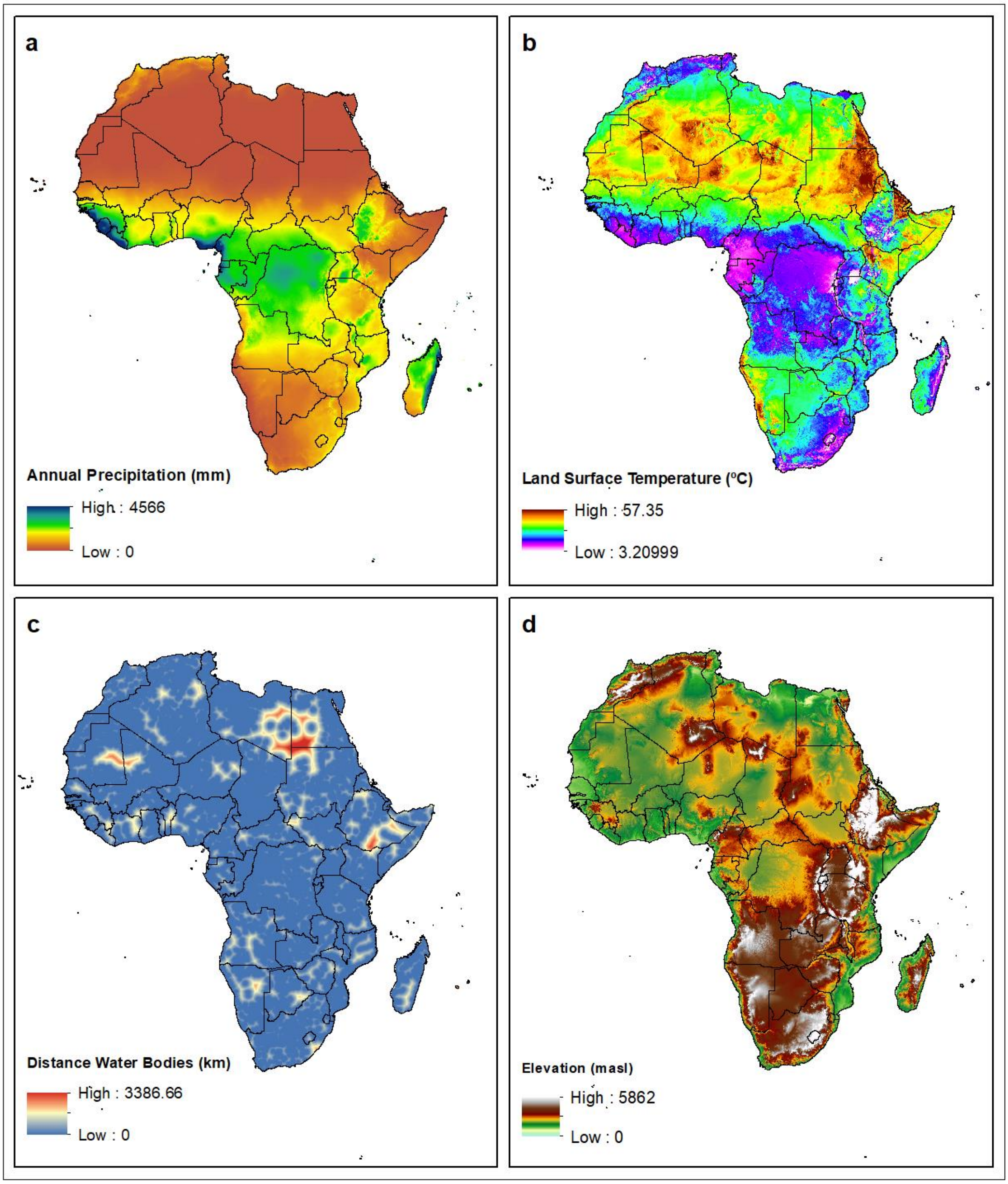

**Figure 2S** Variables used to model environmental suitability of podoconiosis across Africa. A) Enhanced vegetation index (EVI), B) Clay fraction at the top-soil (%), C) Silt fraction at the top-soil (%), and D) pH $H_20$ at the top-soil.

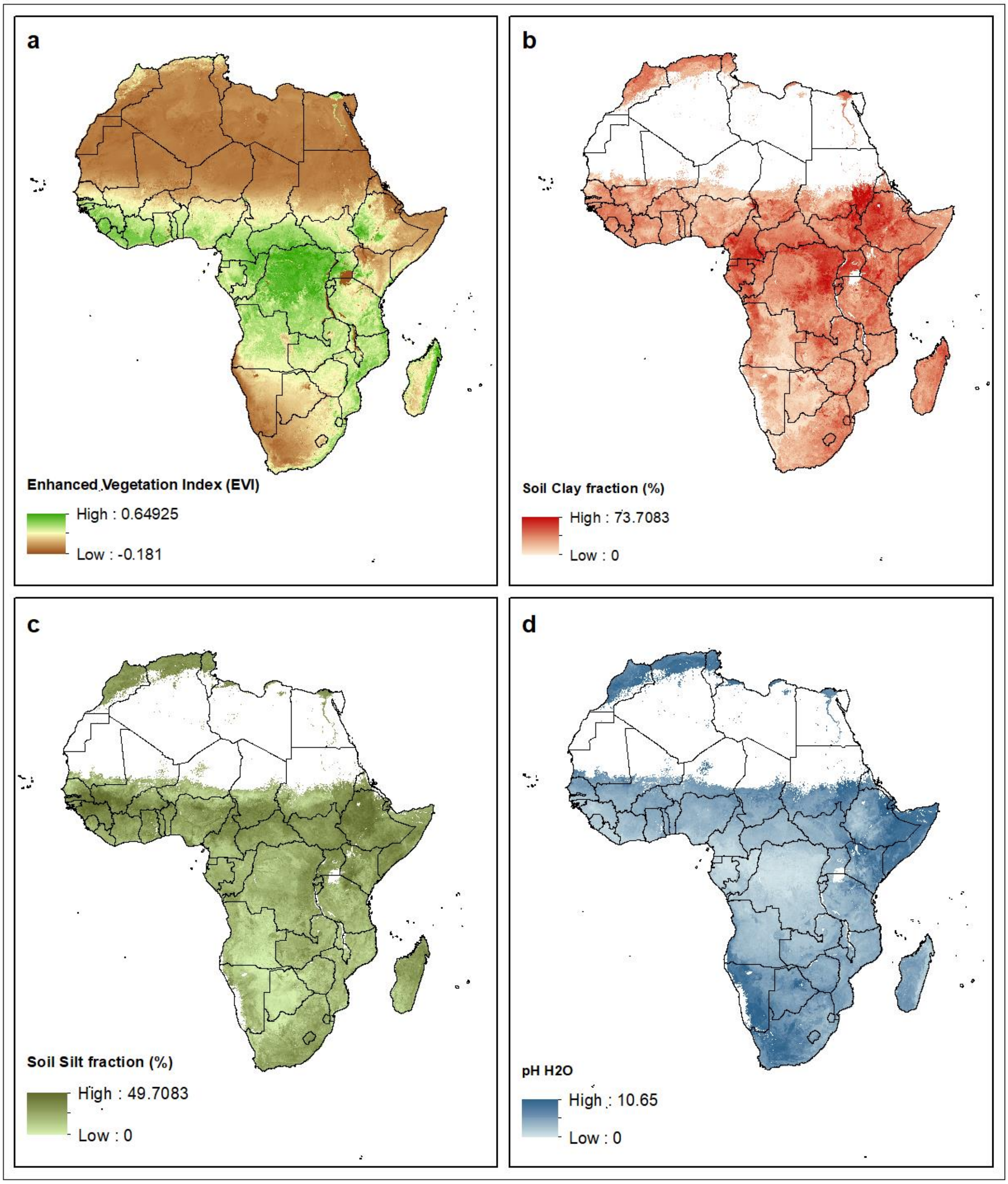

**Figure 3S. Variable contribution of final ensemble models for podoconiosis in Africa based on *boosted regression trees* (BRT) and *random forest* (RF).** Variable contribution is provided as percentage, and it shows the relative contribution of selected environmental predictors to the final ensemble model.

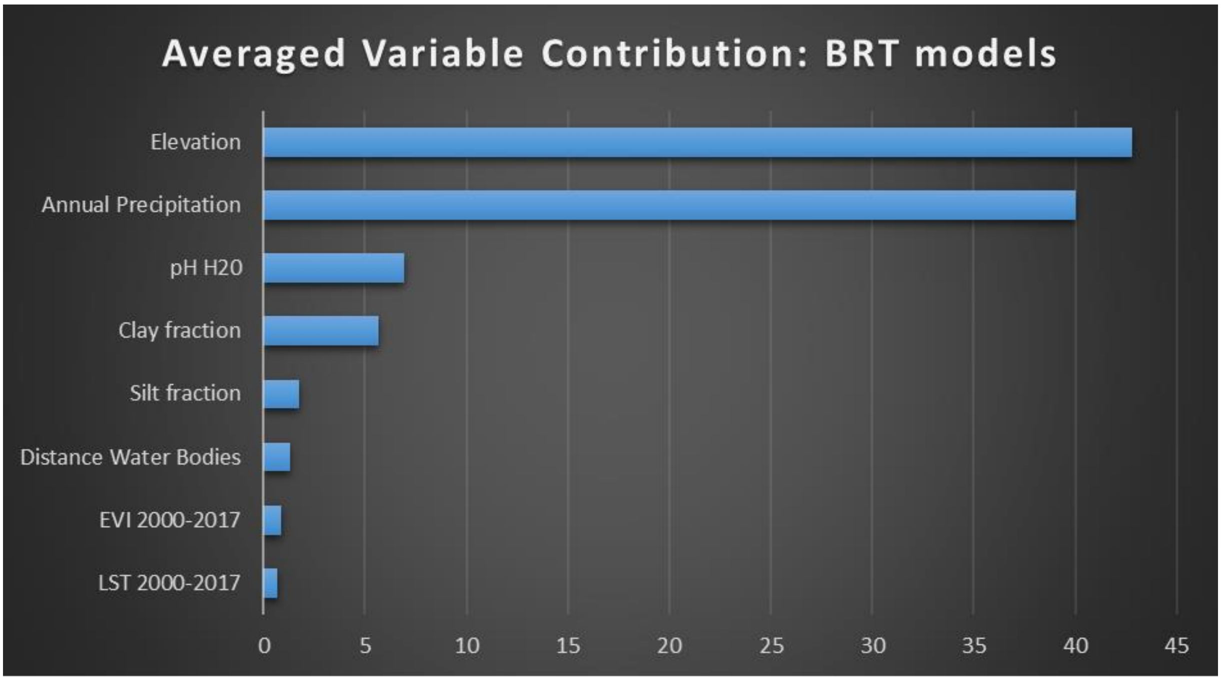


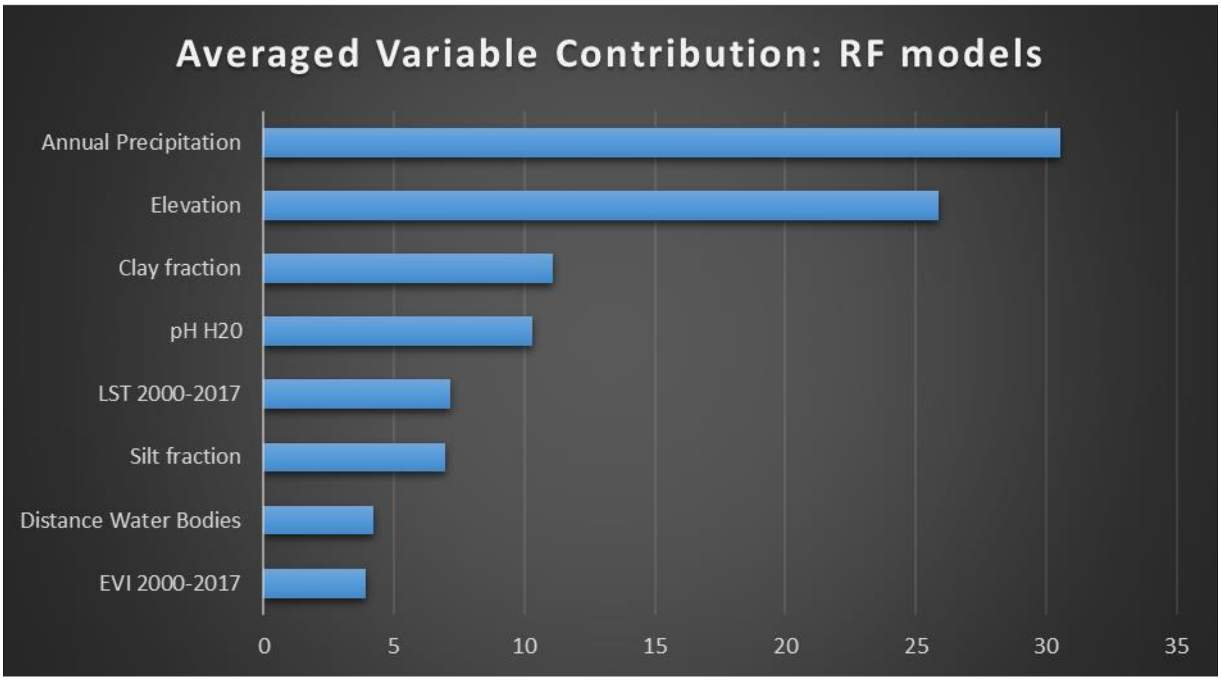

**Figure 4S. Partial dependence plots of the relative contribution of covariates (annual total precipitation, land surface temperature, distance to water bodies and elevation) to the boosted regression tree (BRT) model for podoconiosis, averaged over 50 ensembles.** Blue lines represent the mean partial dependence over all 50 BRT ensembles and grey envelopes the standard deviation from the mean. The *y*-axis is the transformed logit response and *x*-axis is the full range of covariates values.

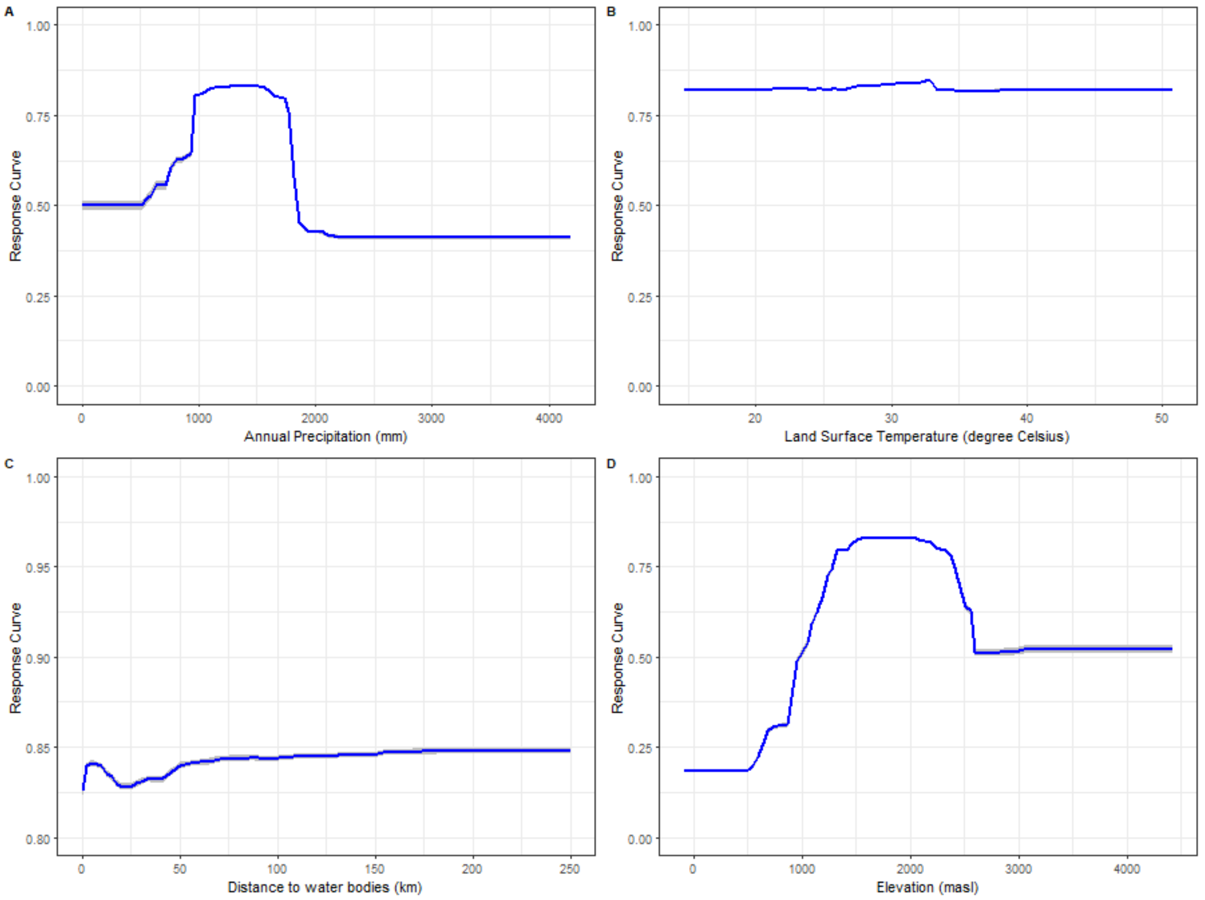

**Figure 5S. Partial dependence plots of the relative contribution of covariates (EVI, clay soil fraction, silt soil fraction, and soil pH-$H_2O$) to the boosted regression tree (BRT) model for podoconiosis, averaged over 50 ensembles.** Blue lines represent the mean partial dependence over all 50 BRT ensembles and grey envelopes the standard deviation from the mean. The *y*-axis is the transformed logit response and *x*-axis is the full range of covariates values.

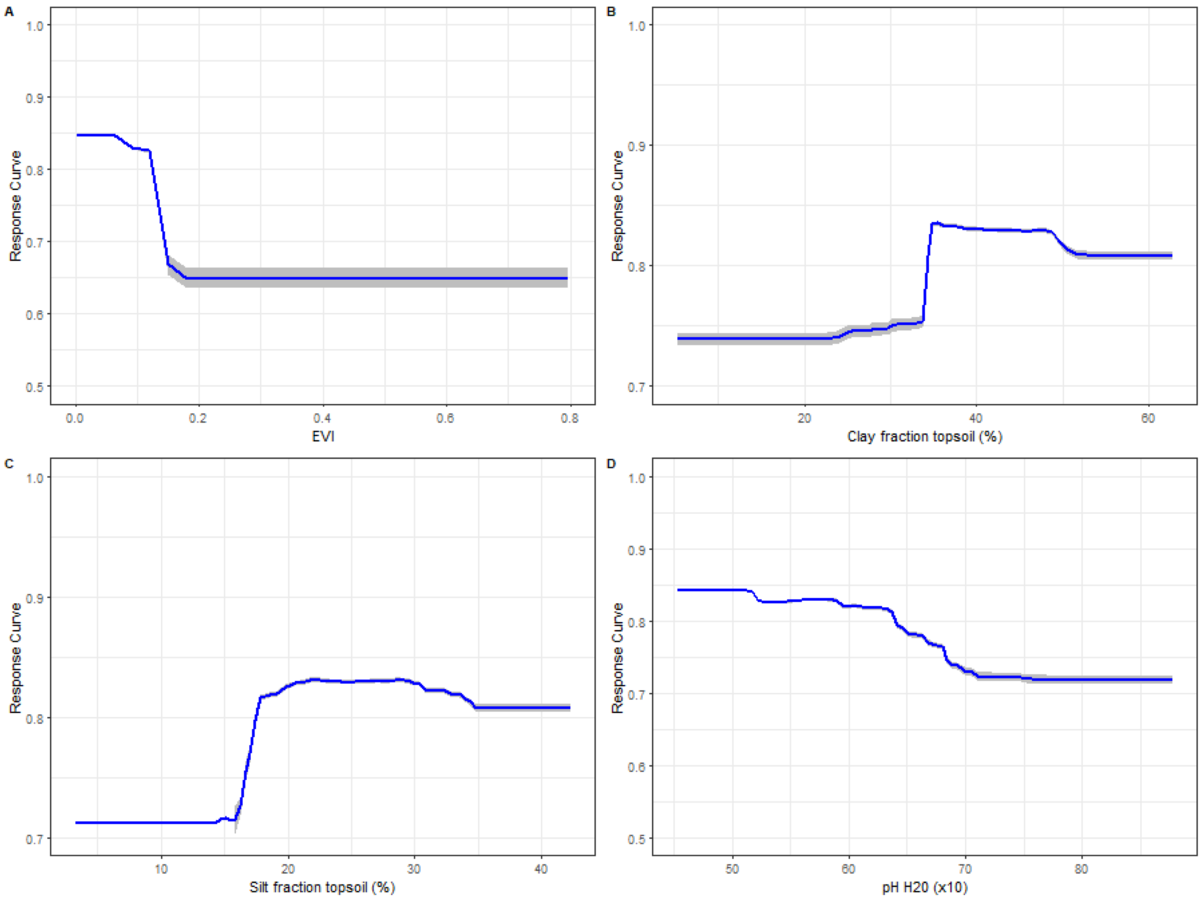

**Figure 6S. Partial dependence plots of the relative contribution of covariates (annual total precipitation, land surface temperature, distance to water bodies and elevation) to the random forest (RF) model for podoconiosis, averaged over 50 ensembles.** Blue lines represent the mean partial dependence over all 50 RF ensembles and grey envelopes the standard deviation from the mean. The *y*-axis is the transformed logit response and *x*-axis is the full range of covariates values.

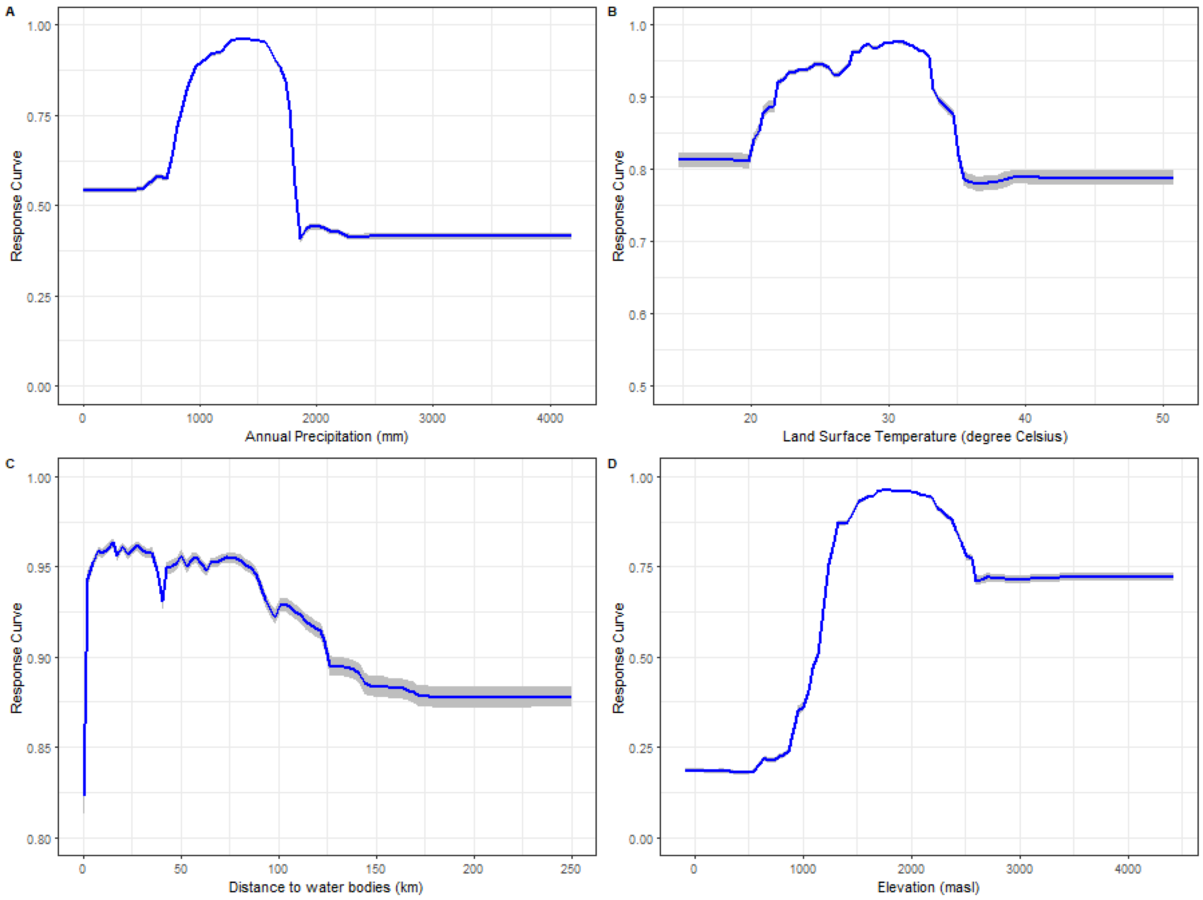

**Figure 7S. Partial dependence plots of the relative contribution of covariates (EVI, clay soil fraction, silt soil fraction, and soil pH-$H_2O$) to the random forest (RF) model for podoconiosis, averaged over 50 ensembles.** Blue lines represent the mean partial dependence over all 50 RF ensembles and grey envelopes the standard deviation from the mean. The *y*-axis is the transformed logit response and *x*-axis is the full range of covariates values.

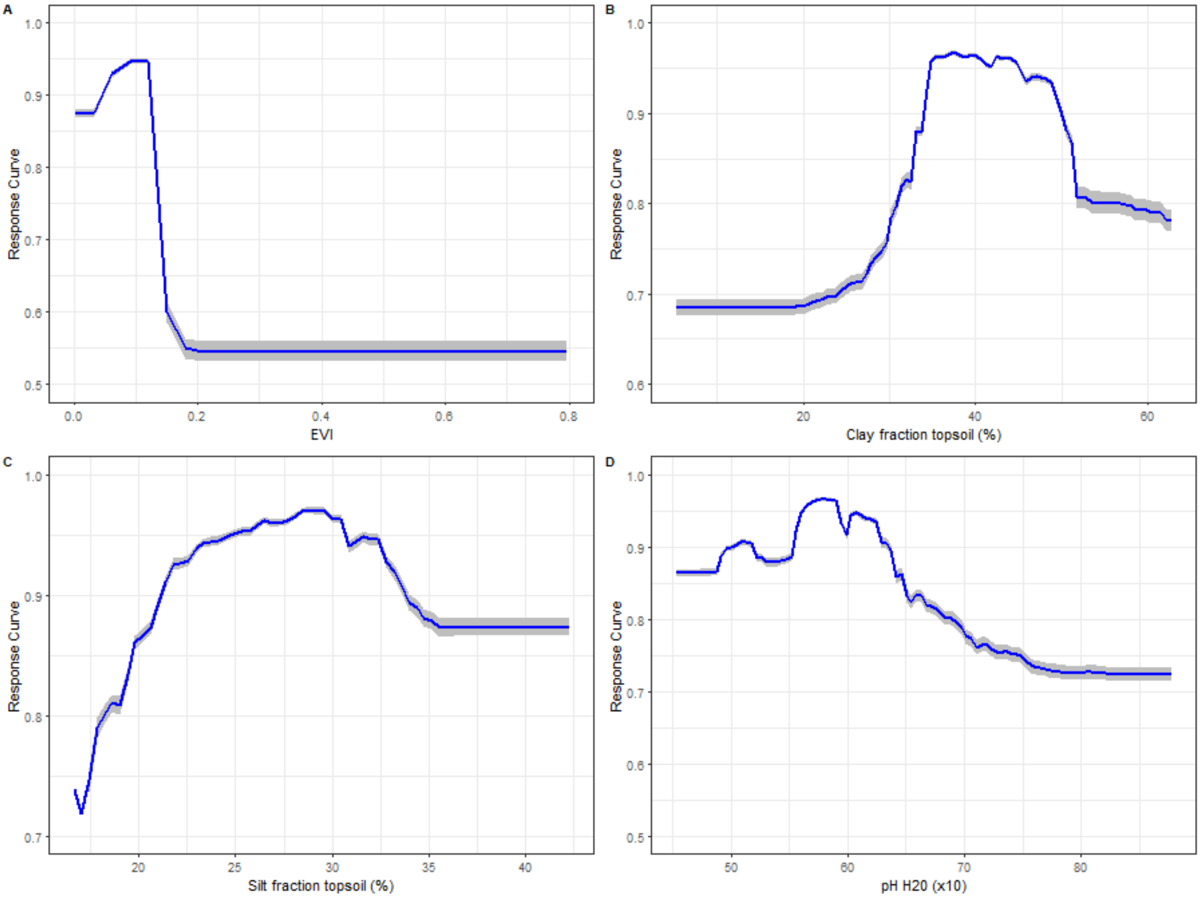

**Figure 8S. Map displaing randomly setting up spatial units (5 regular blocks) for internal cross-validation.** The whole of Africa was split in to five regular squares of 2,000km by 2,000km and then randomly tagged from 1 to 5. Single models were trained using all the data within 4 out of the 5 folds (train data) and the data in the remaining fold (test data) were used to assess the quality of the predictive performance of the fitted model.

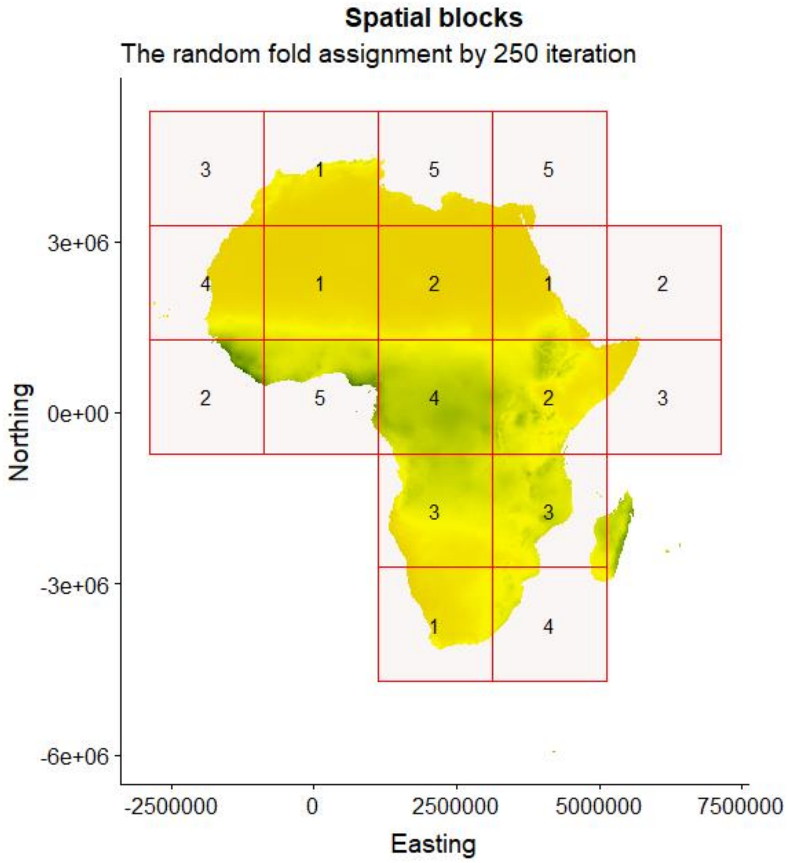

**Figure 9S. Environmental suitability for podoconiosis across Africa and prediction uncertainty (95% confidence interval)** as estimated by the final ensemble model constructed through five regular spatial blocks. This model showed a high correlation (Pearson's correlation coefficient of 0.9902) with the final ensemble model constructed using a 20% heldout cross-validation apprioach (single model trained using randomly selected 80% point records)

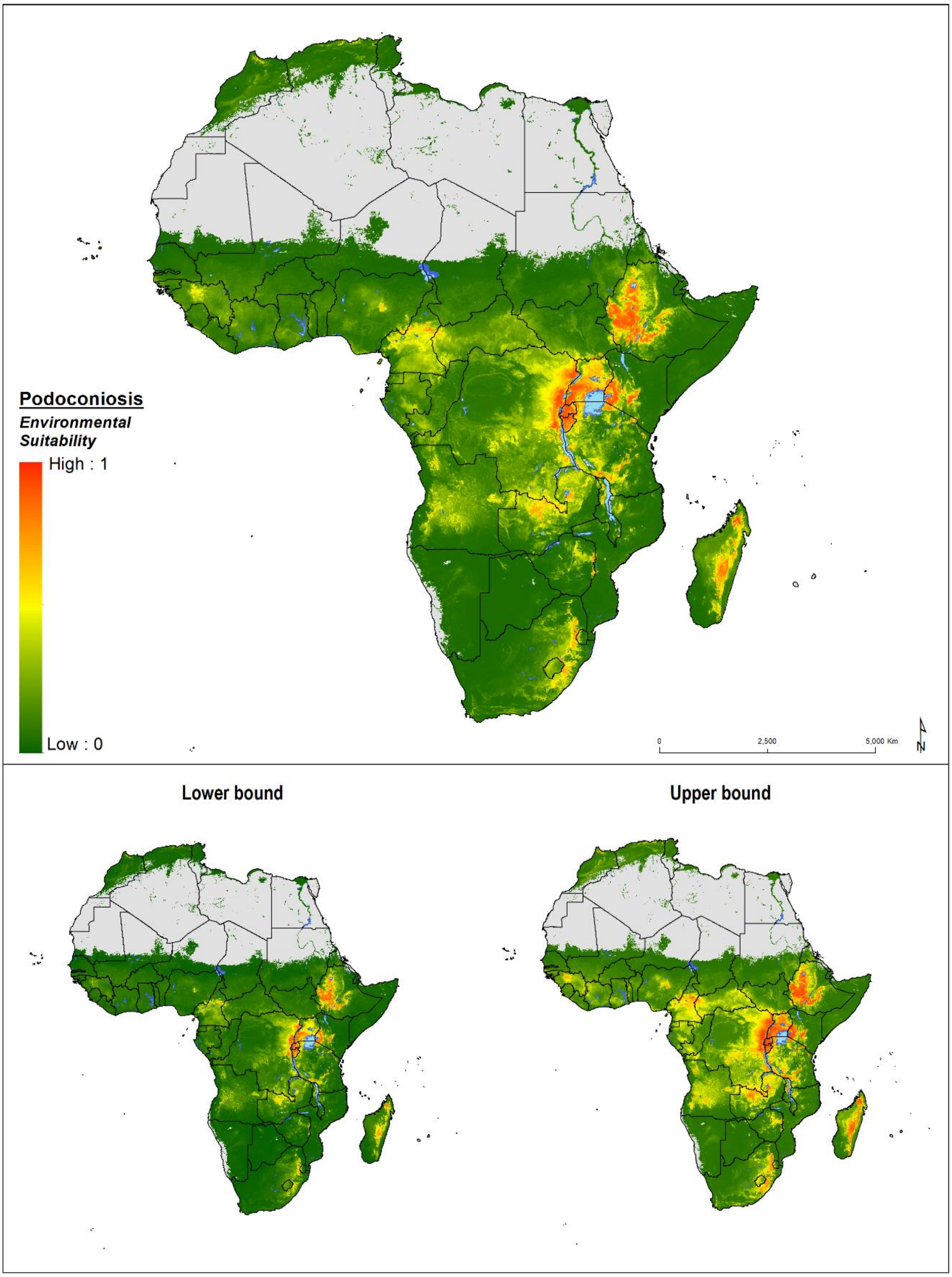

**Figure 10S. Map of GBD regions.** Modelling regions were defined as the five GBD regions of Central (central SSA), East (eastern SSA), North (North Africa and the Middle East), South (southern SSA) and West Africa (western SSA)[1]. As this study was limited to mainland Africa and African island nations, select countries were excluded from the North Africa and Middle East region (Afghanistan, Bahrain, Iran, Iraq, Jordan, Kuwait, Lebanon, Oman, Palestinian territories, Qatar, Saudi Arabia, Syria, Turkey, United Arab Emirates, and Yemen). Western Sahara was included as part of the North region.

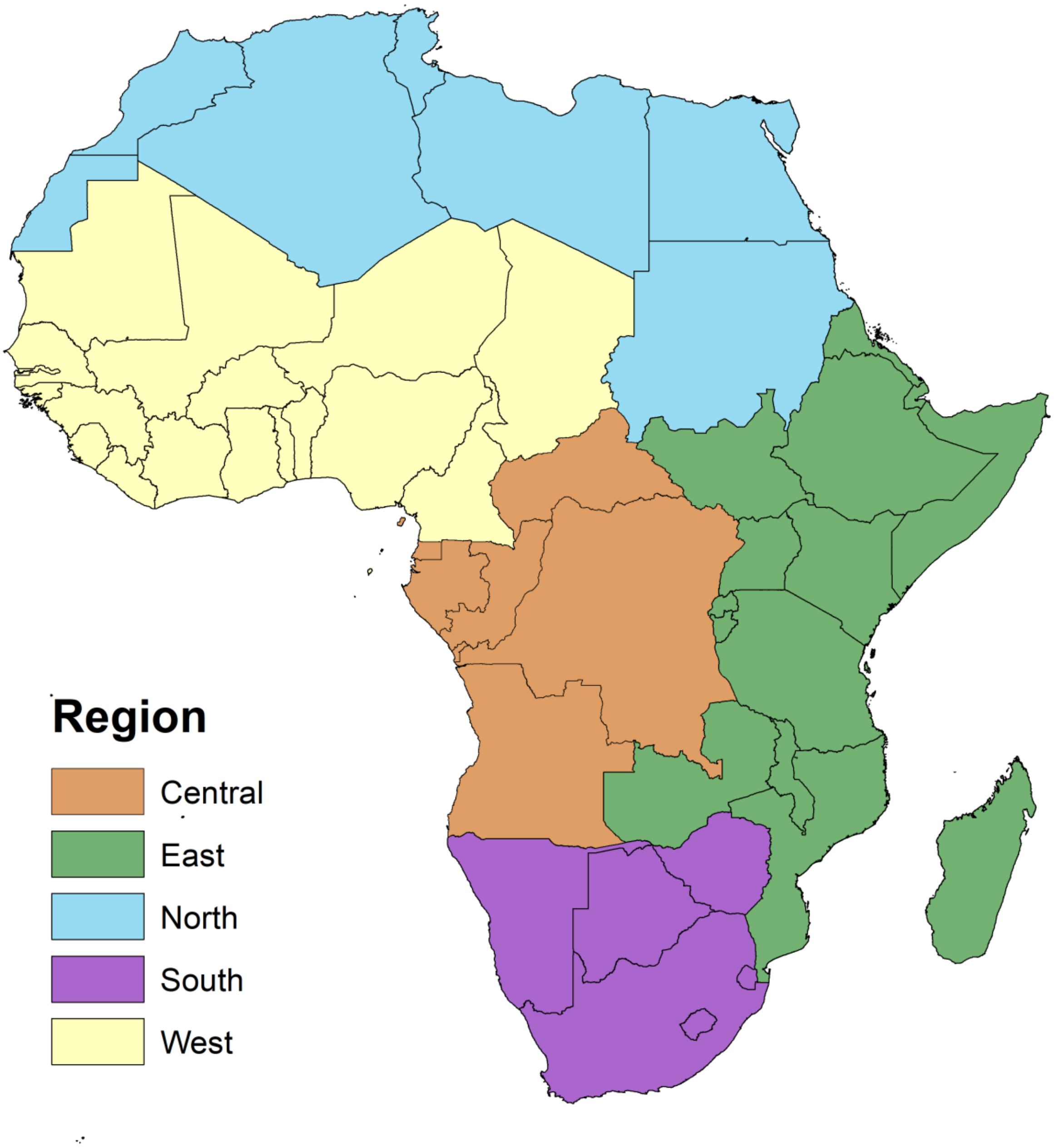

**Table 1S. Median and 95% confidence intervals of validation indicators (TSS, AUC and PCC) for the modelling approaches tested: generalized linear models (GLM), generalized additive models (GAM), boosted regression trees (BRT), artificial neural networks (ANN), multiple adaptive regression splines (MARS), and random forest (RF).** TSS: true skill statistic, ROC: the area under the receiver operation characteristic (ROC) curve, and PCC: positive ("occurrence") correctly classified.

| | GLM | | GAM | | GBM | | ANN | | MARS | | RF | |
|---|---|---|---|---|---|---|---|---|---|---|---|---|
| | **Median** | **95%CI** | **Median** | **95%CI** | **Median** | **95%CI** | **Median** | **95%CI** | **Median** | **95%CI** | **Median** | **95%CI** |
| **TSS** | 0.55 | 0.53 - 0.57 | 0.56 | 0.54 - 0.58 | 0.61 | 0.6 - 0.63 | 0.56 | 0.54 - 0.58 | 0.54 | 0.53 - 0.56 | 0.61 | 0.6 - 0.63 |
| **AUC** | 0.85 | 0.84 - 0.86 | 0.86 | 0.86 - 0.88 | 0.89 | 0.89 - 0.9 | 0.85 | 0.85 - 0.87 | 0.85 | 0.85 - 0.87 | 0.88 | 0.88 - 0.9 |
| **PCC** | 0.78 | 0.78 - 0.79 | 0.80 | 0.8 - 0.81 | 0.82 | 0.82 - 0.83 | 0.81 | 0.8 - 0.82 | 0.79 | 0.79 - 0.8 | 0.83 | 0.83 - 0.84 |
