## Supplementary Appendix 2 for "Predicting the Environmental Suitability and Population at Risk of Podoconiosis in Africa"

### Angola

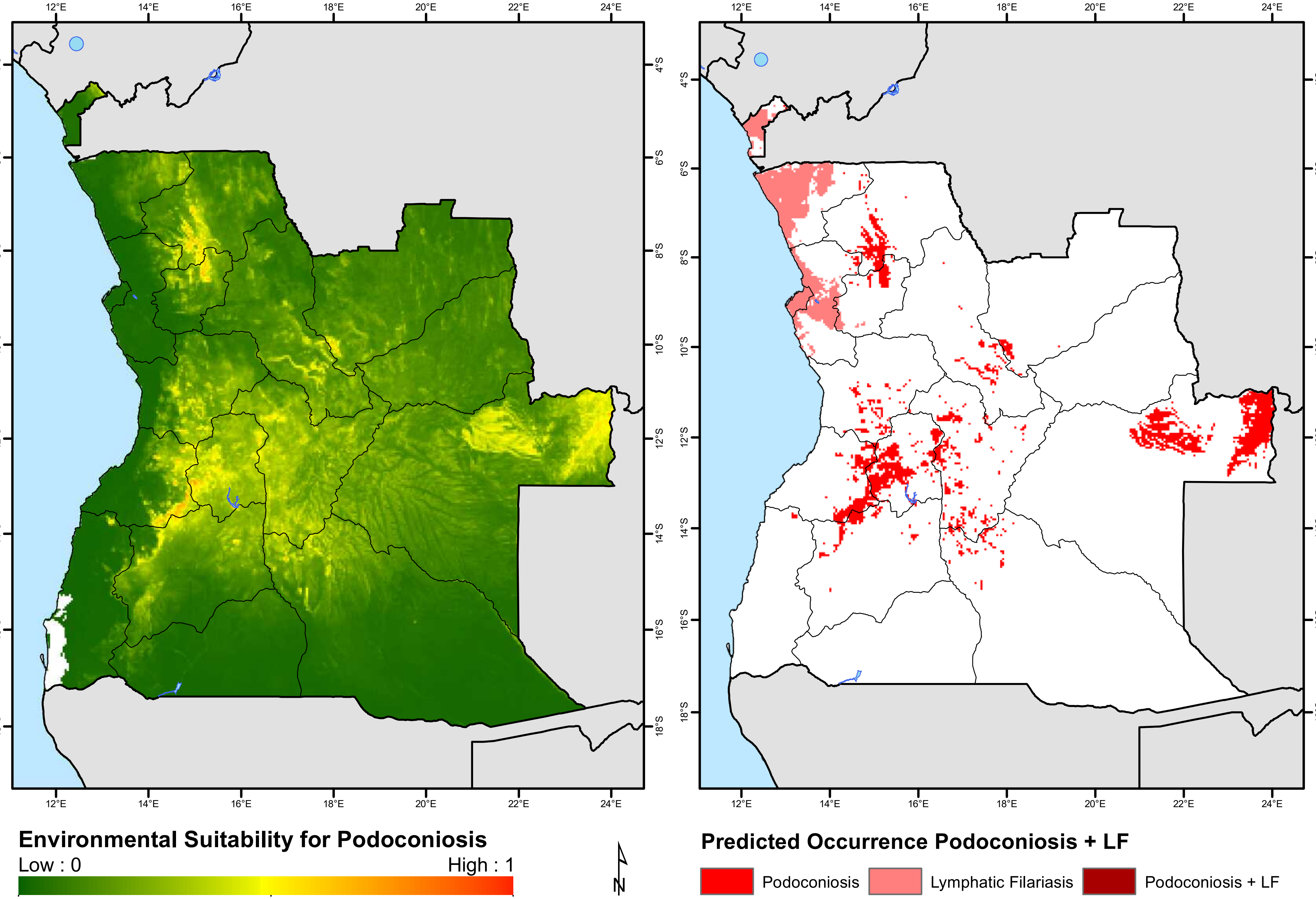
12°E
14°E
16°E
18°E
20°E
22°E
24°E
4°S
6°S
8°S
10°S
12°S
14°S
16°S
18°S
Environmental Suitability for Podoconiosis
Low : 0
High : 1
N
Predicted Occurrence Podoconiosis + LF
Podoconiosis
Lymphatic Filariasis
Podoconiosis + LF

### Benin

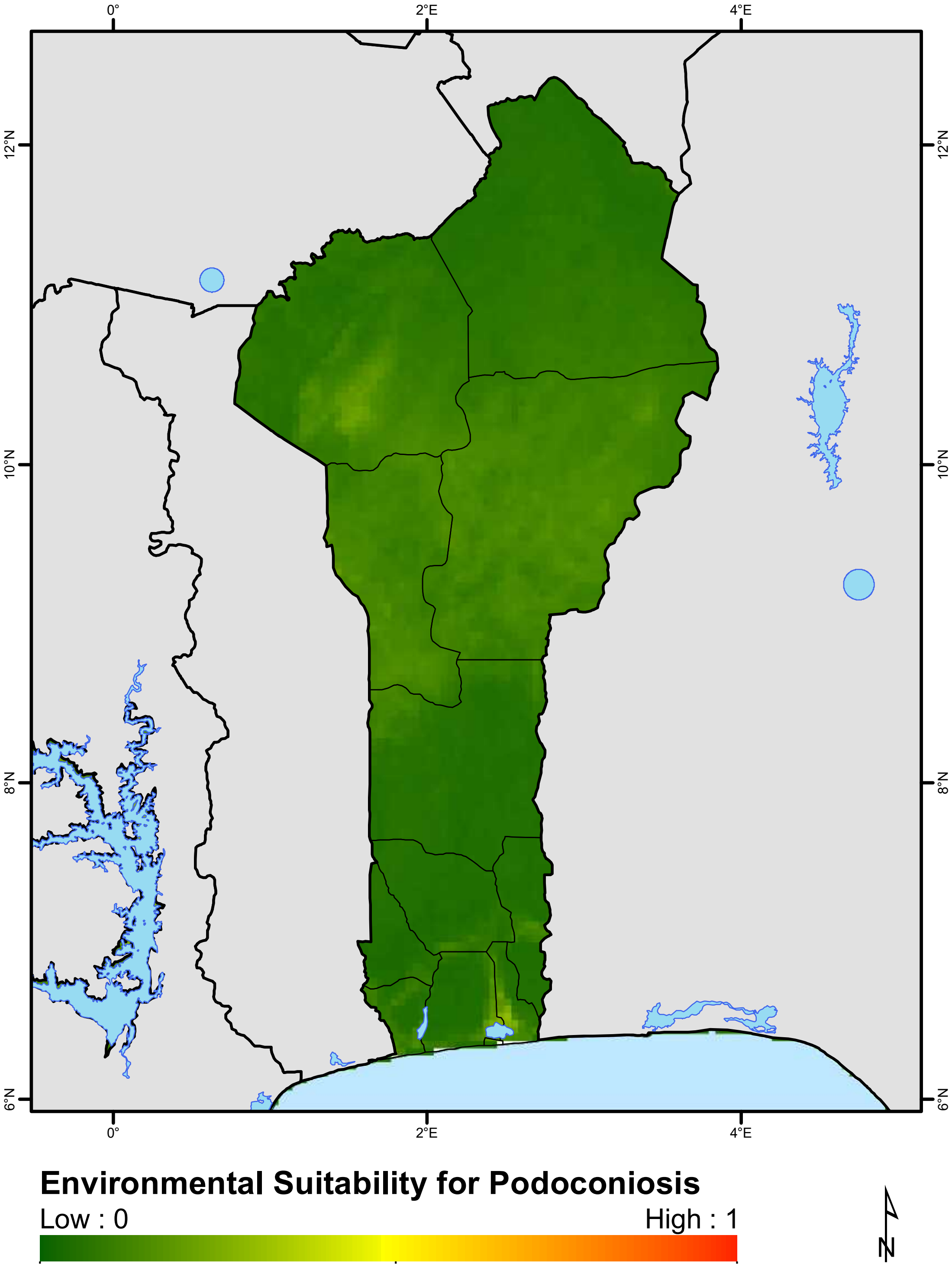
0°
2°E
4°E
12°N
10°N
8°N
6°N
Environmental Suitability for Podoconiosis
Low : 0
High : 1
N

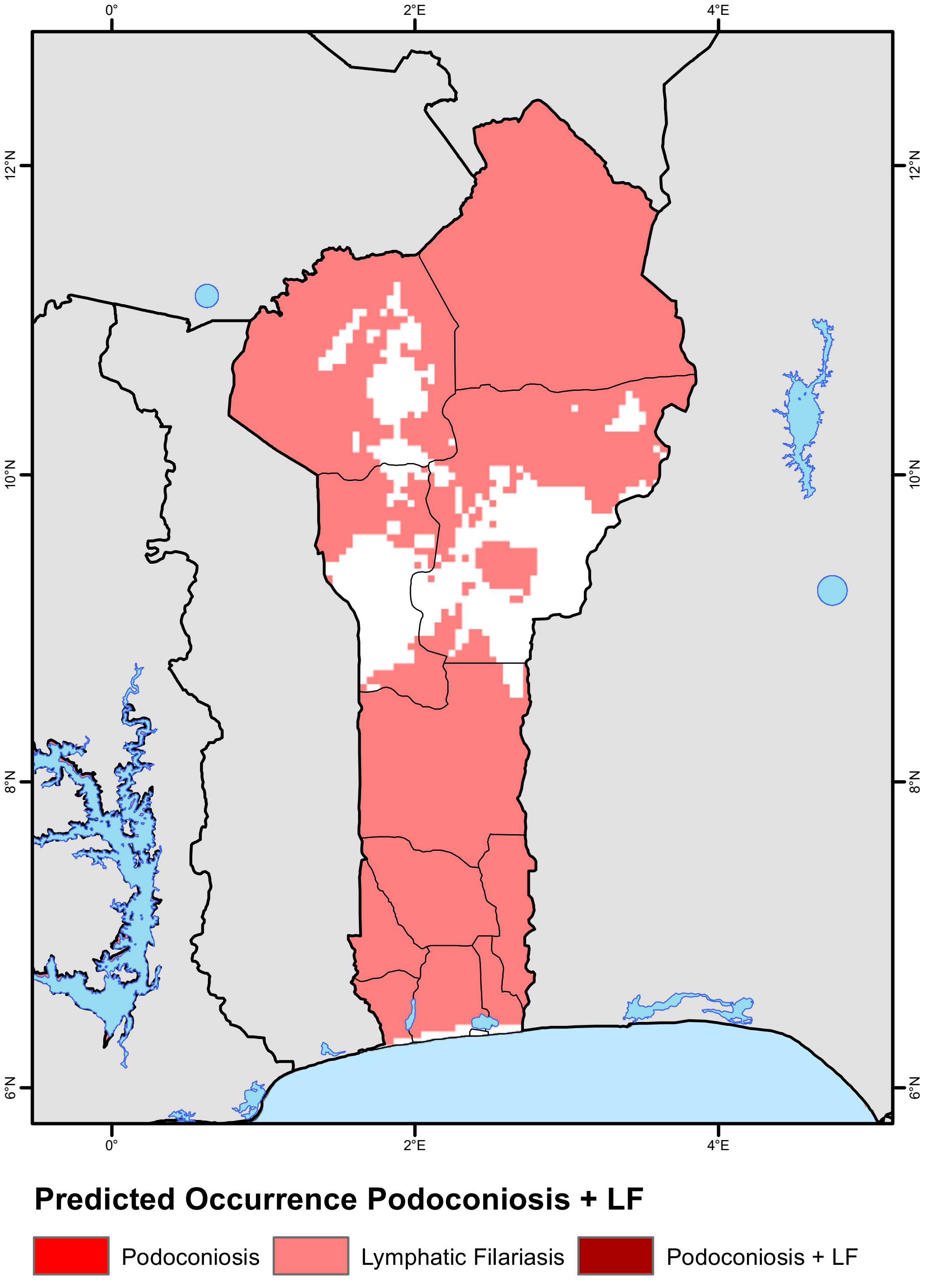
0°
2°E
4°E
12°N
10°N
8°N
6°N
Predicted Occurrence Podoconiosis + LF
Podoconiosis
Lymphatic Filariasis
Podoconiosis + LF

### Burkina Faso

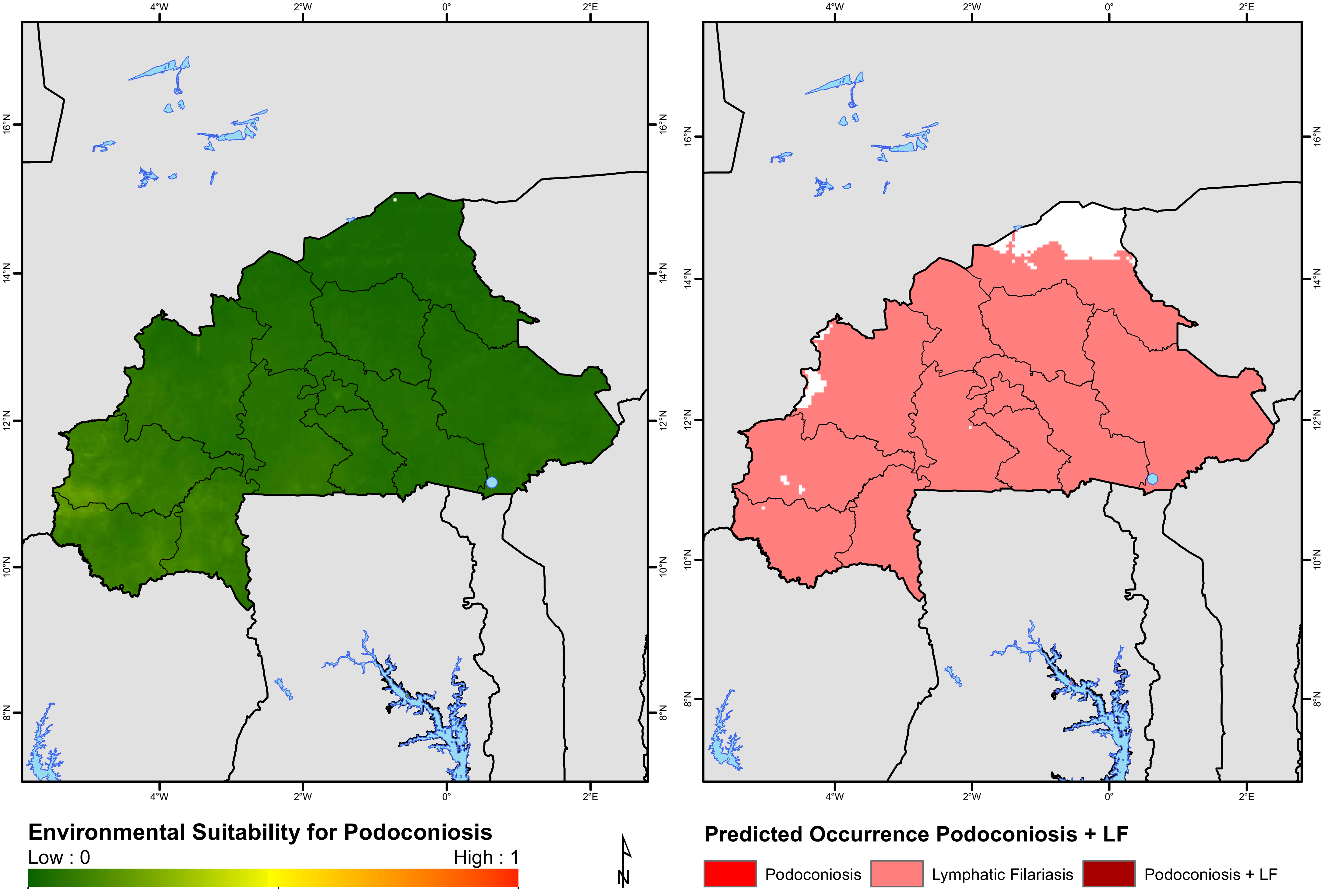

### Burundi

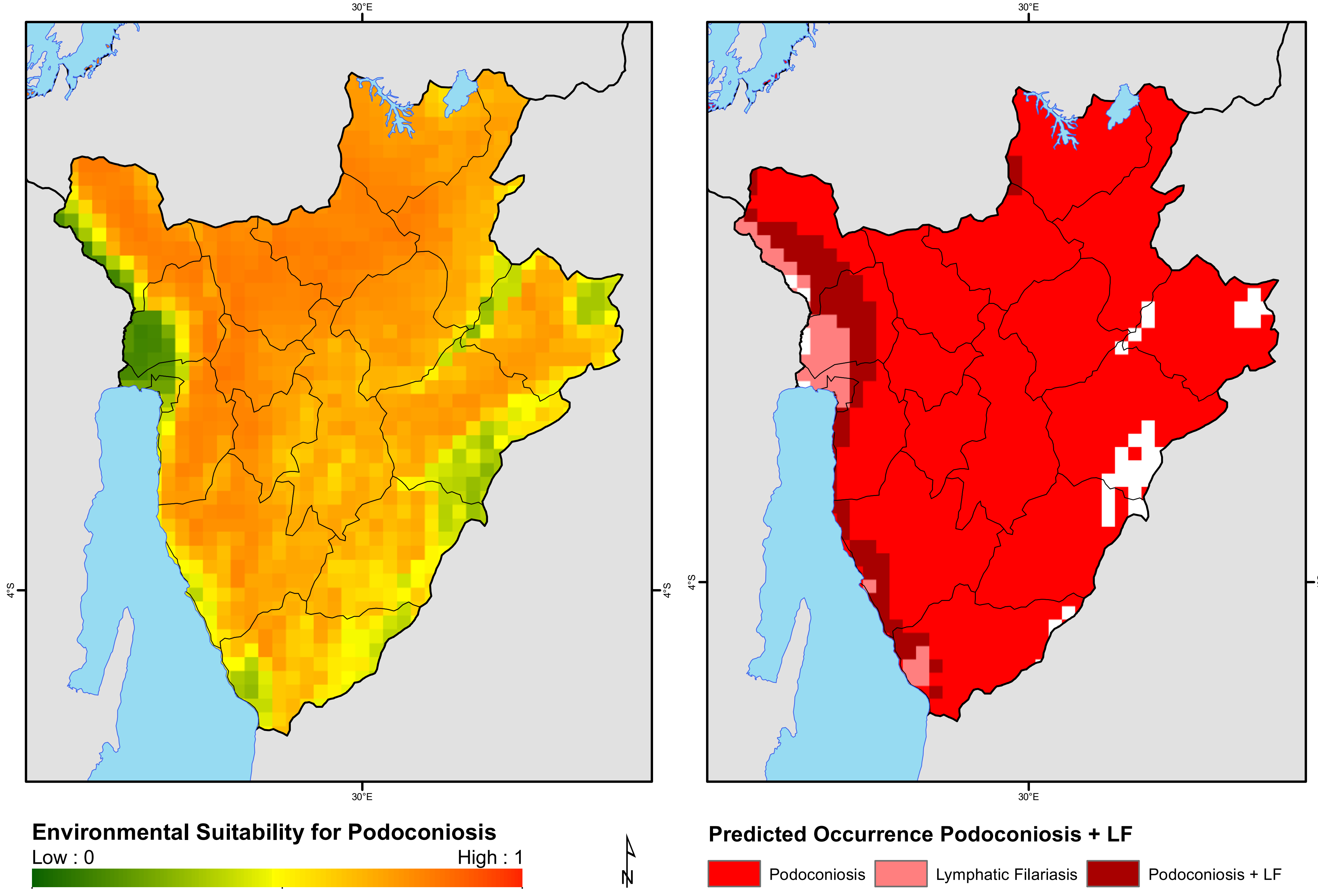
30°E
30°E
4°S
4°S
30°E
30°E
4°S
4°S
Environmental Suitability for Podoconiosis
Low : 0
High : 1
N
Predicted Occurrence Podoconiosis + LF
Podoconiosis
Lymphatic Filariasis
Podoconiosis + LF

### Cameroon

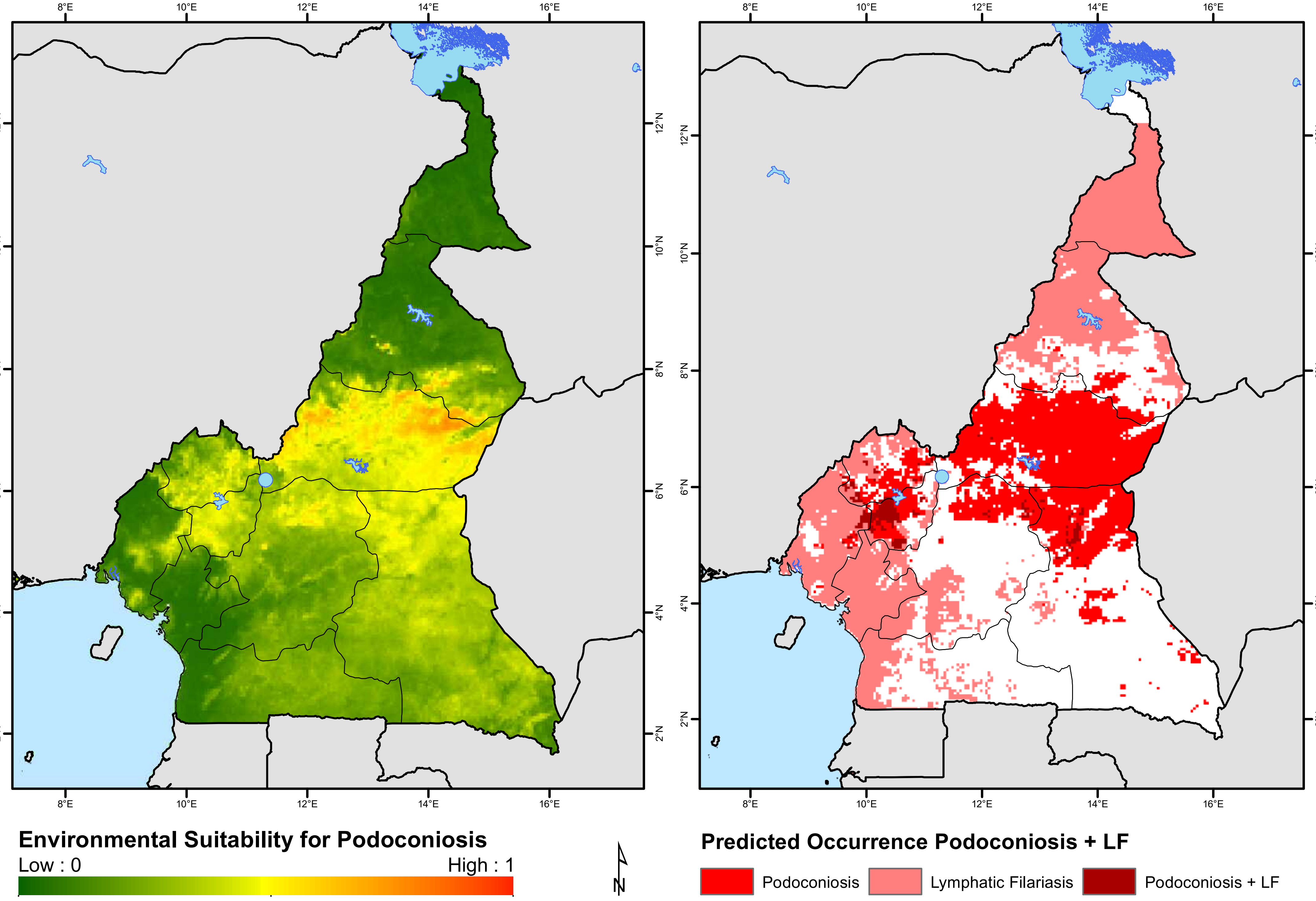
8°E
10°E
12°E
14°E
16°E
2°N
4°N
6°N
8°N
10°N
12°N
Environmental Suitability for Podoconiosis
Low : 0
High : 1
N
Predicted Occurrence Podoconiosis + LF
Podoconiosis
Lymphatic Filariasis
Podoconiosis + LF

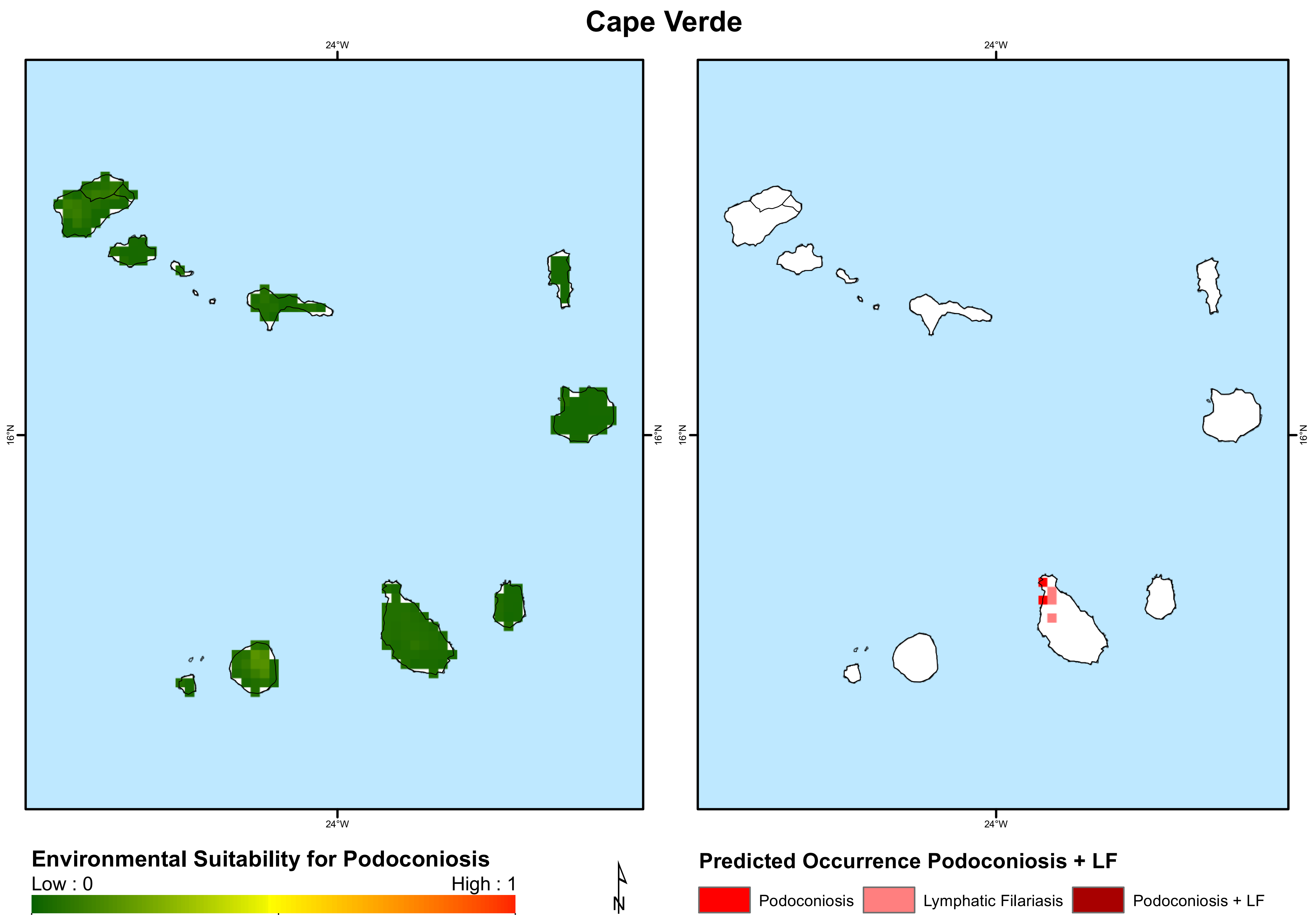
Cape Verde
24°W
16°N
24°W
16°N
Environmental Suitability for Podoconiosis
Low : 0
High : 1
N
Predicted Occurrence Podoconiosis + LF
Podoconiosis
Lymphatic Filariasis
Podoconiosis + LF

### Central African Republic

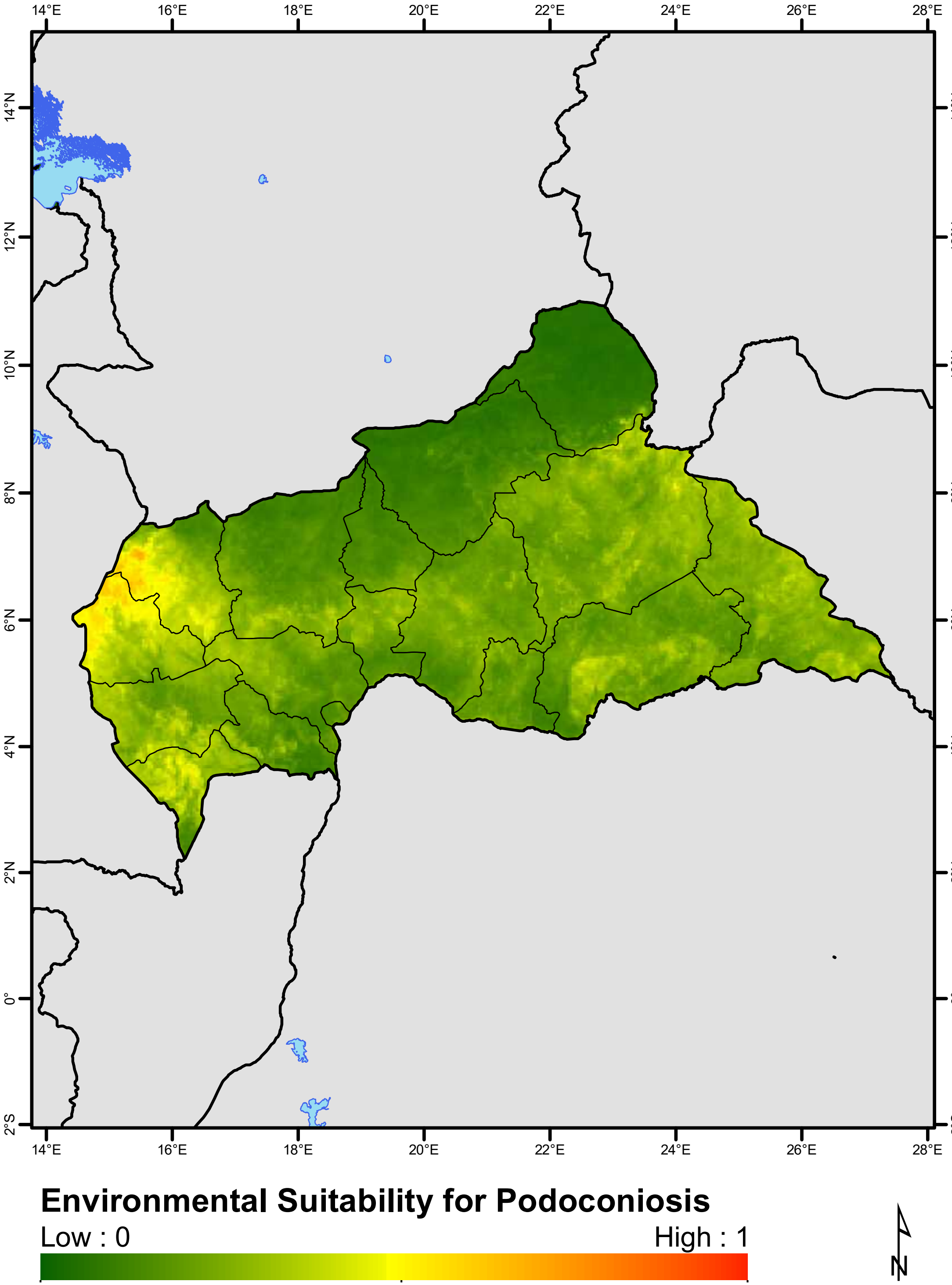


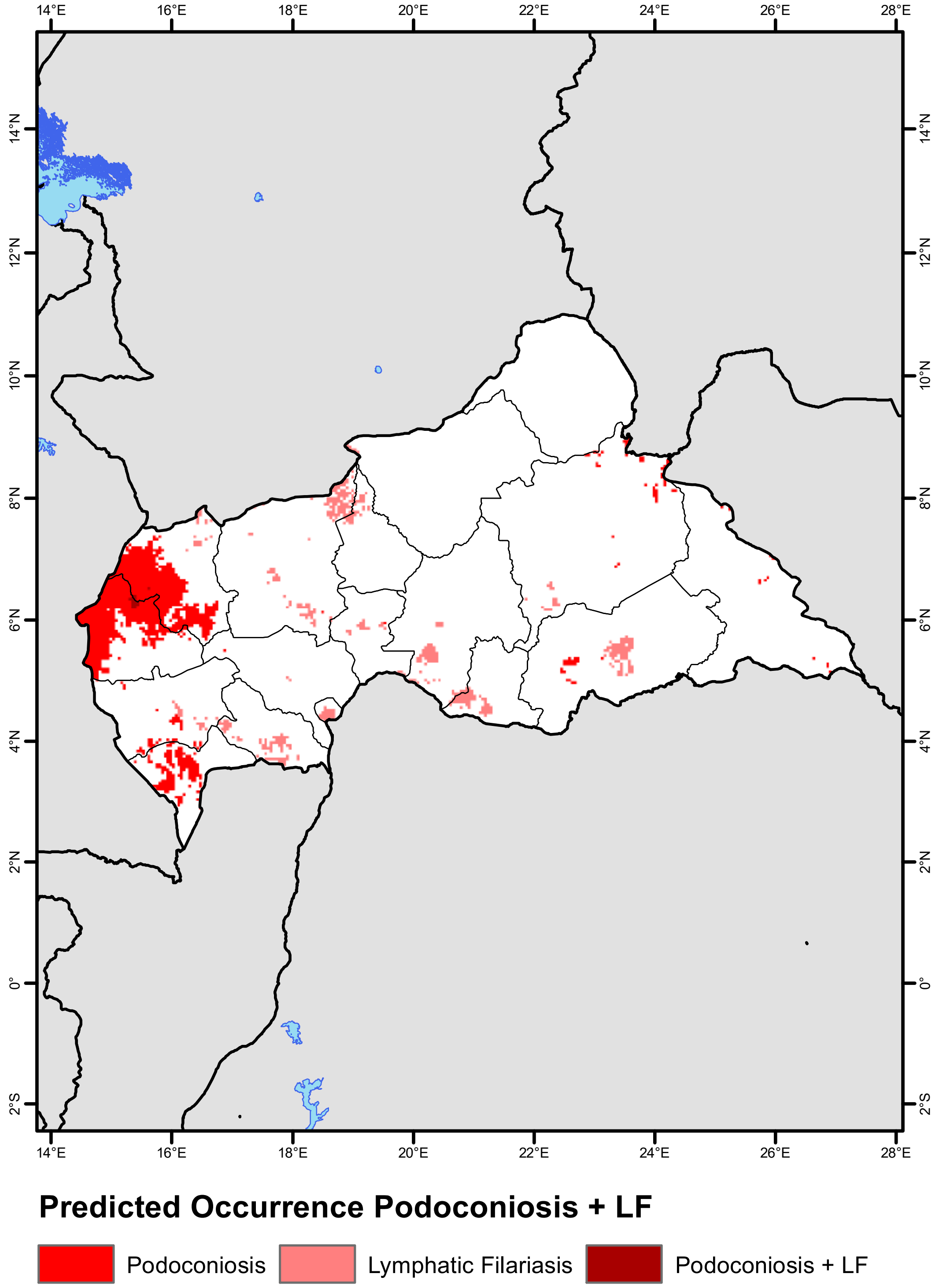

### Chad

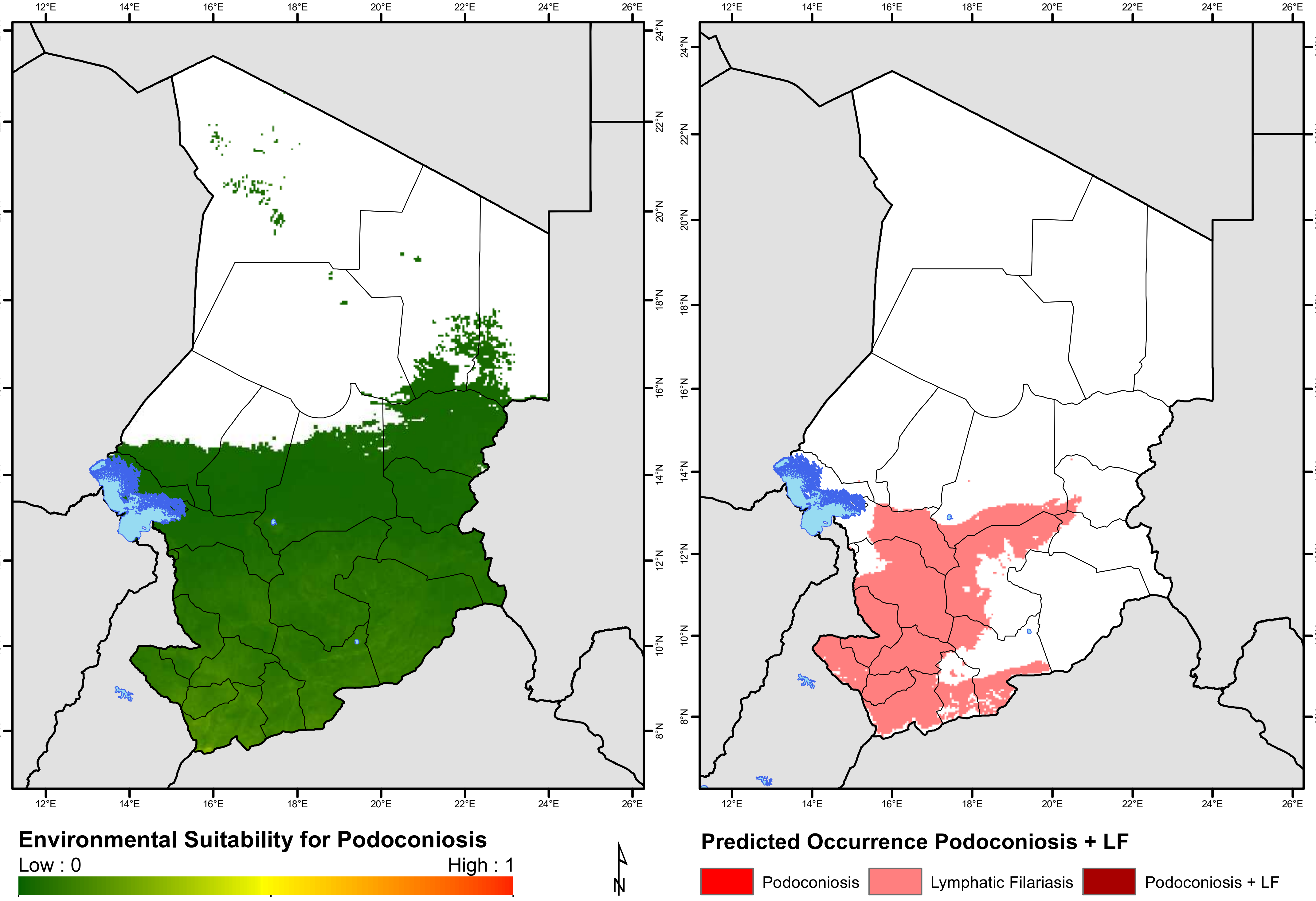
Environmental Suitability for Podoconiosis
Low : 0
High : 1
Predicted Occurrence Podoconiosis + LF
Podoconiosis
Lymphatic Filariasis
Podoconiosis + LF
N
12°E
14°E
16°E
18°E
20°E
22°E
24°E
26°E
8°N
10°N
12°N
14°N
16°N
18°N
20°N
22°N
24°N

### Congo

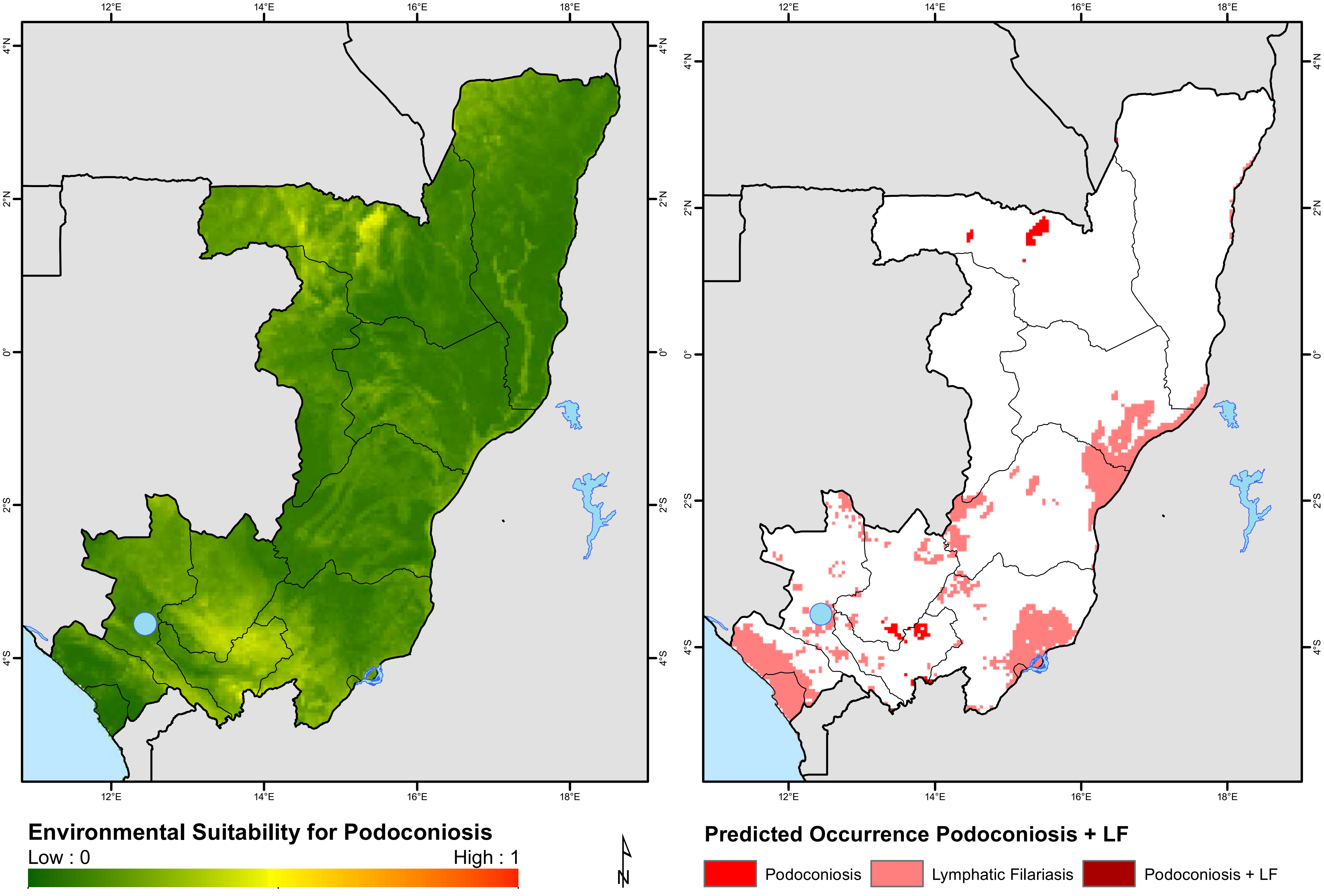

12°E
14°E
16°E
18°E
4°N
2°N
0°
2°S
4°S
Environmental Suitability for Podoconiosis
Low : 0
High : 1
N
Predicted Occurrence Podoconiosis + LF
Podoconiosis
Lymphatic Filariasis
Podoconiosis + LF

### Côte d'Ivoire

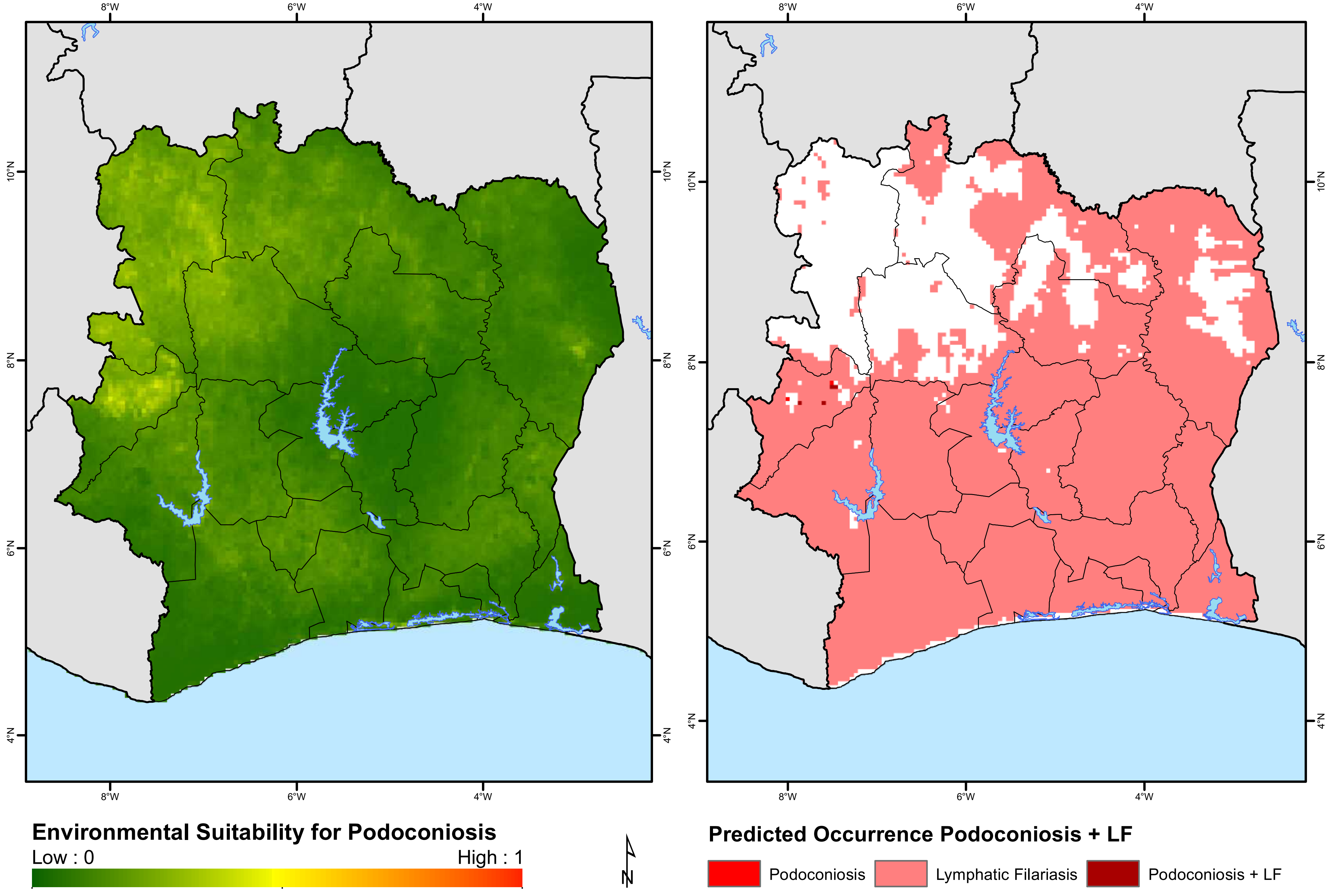

### Democratic Republic of Congo

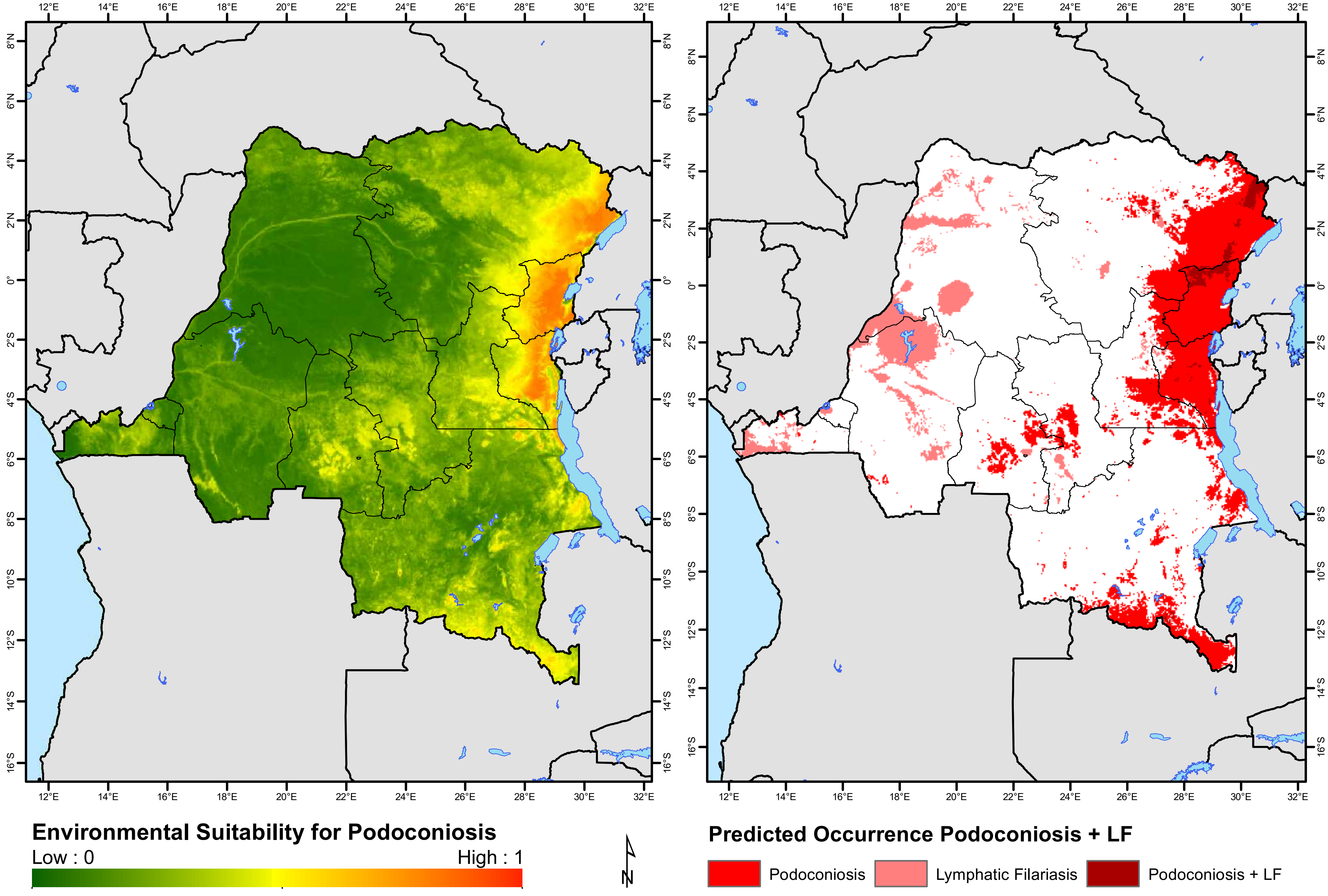

### Djibouti

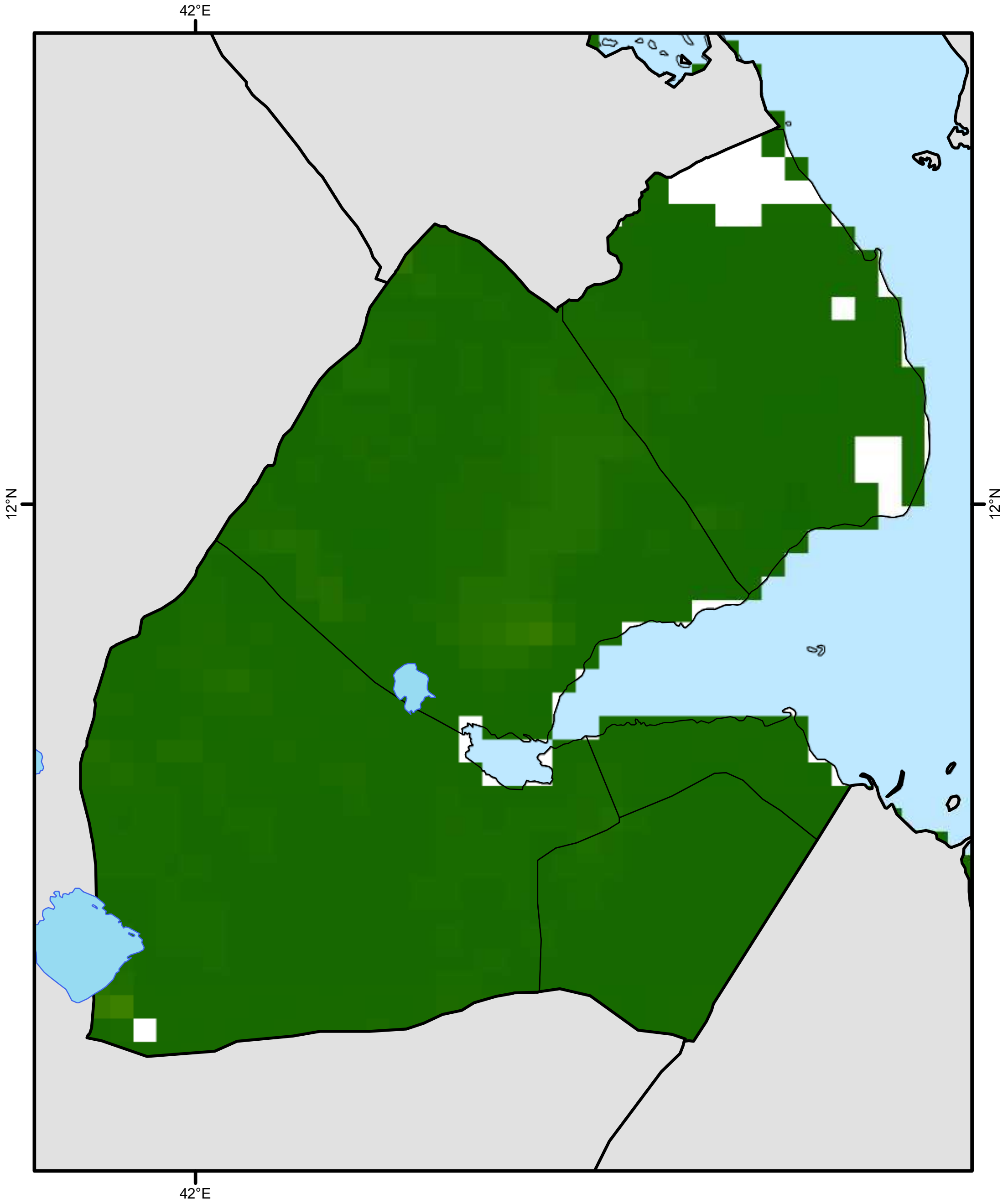


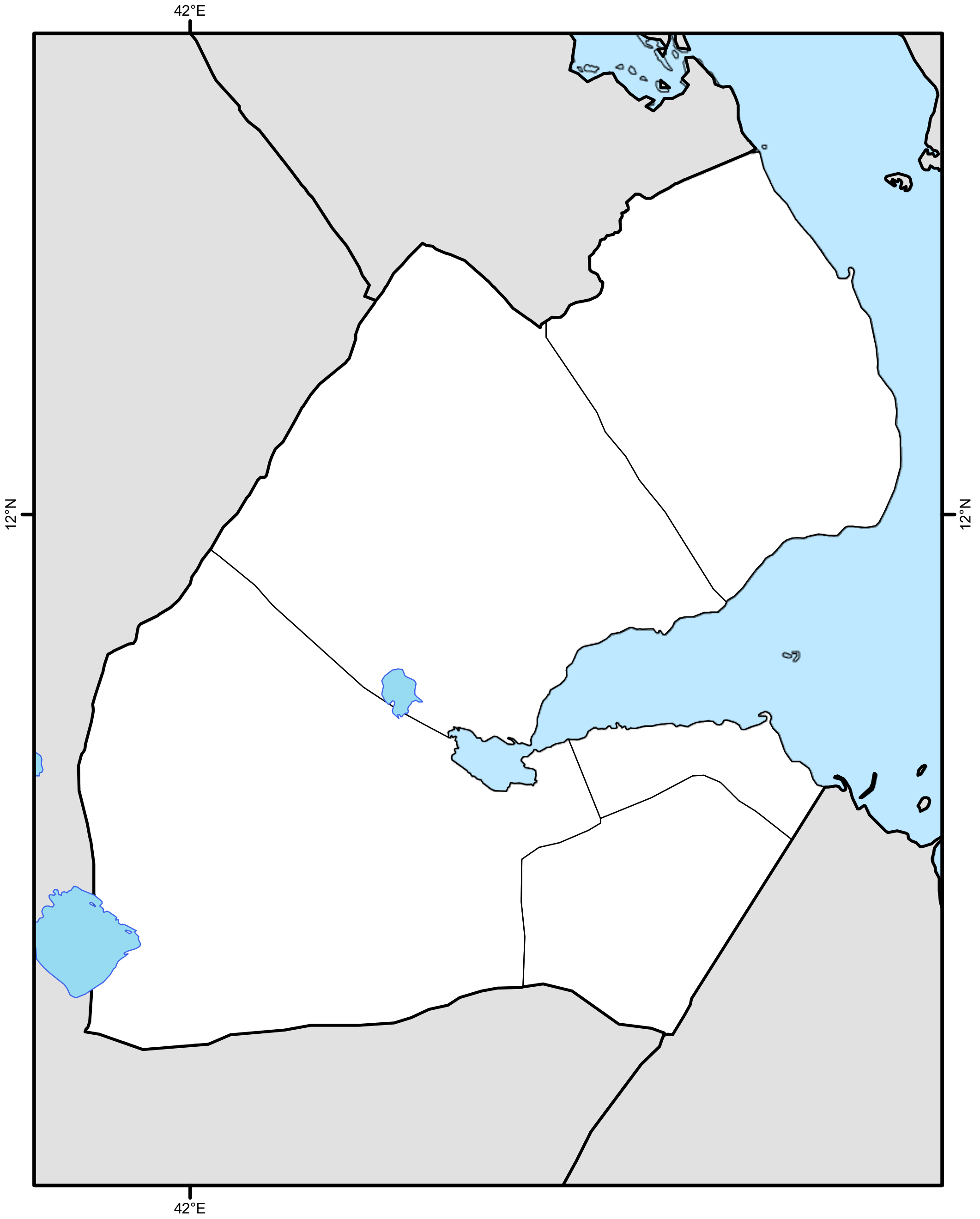


**Environmental Suitability for Podoconiosis**

Low : 0 High : 1

**Predicted Occurrence Podoconiosis + LF**

Podoconiosis | Lymphatic Filariasis | Podoconiosis + LF

### Equatorial Guinea

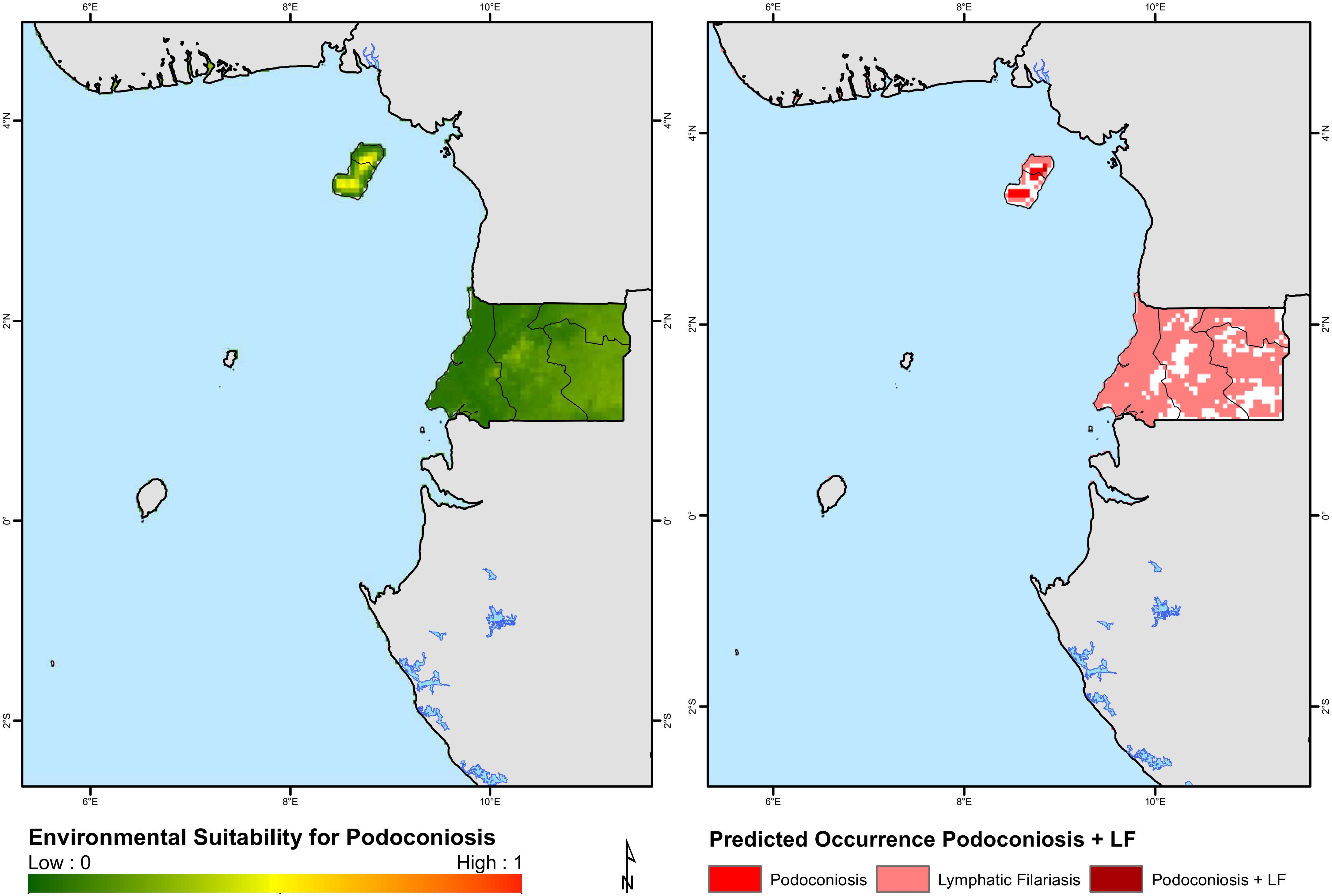

### Eritrea

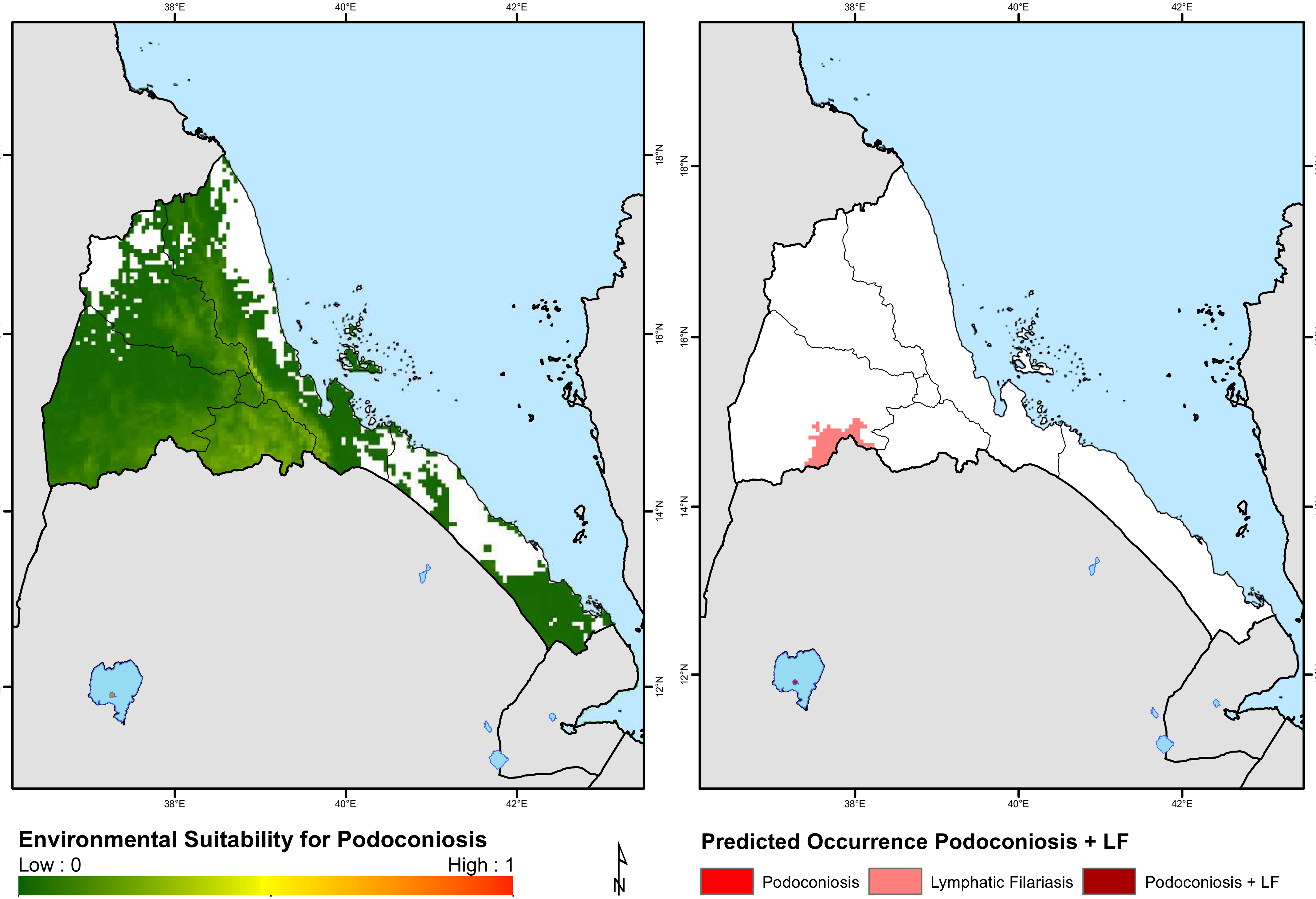

38°E
40°E
42°E
18°N
16°N
14°N
12°N
Environmental Suitability for Podoconiosis
Low : 0
High : 1
N
Predicted Occurrence Podoconiosis + LF
Podoconiosis
Lymphatic Filariasis
Podoconiosis + LF

### Ethiopia

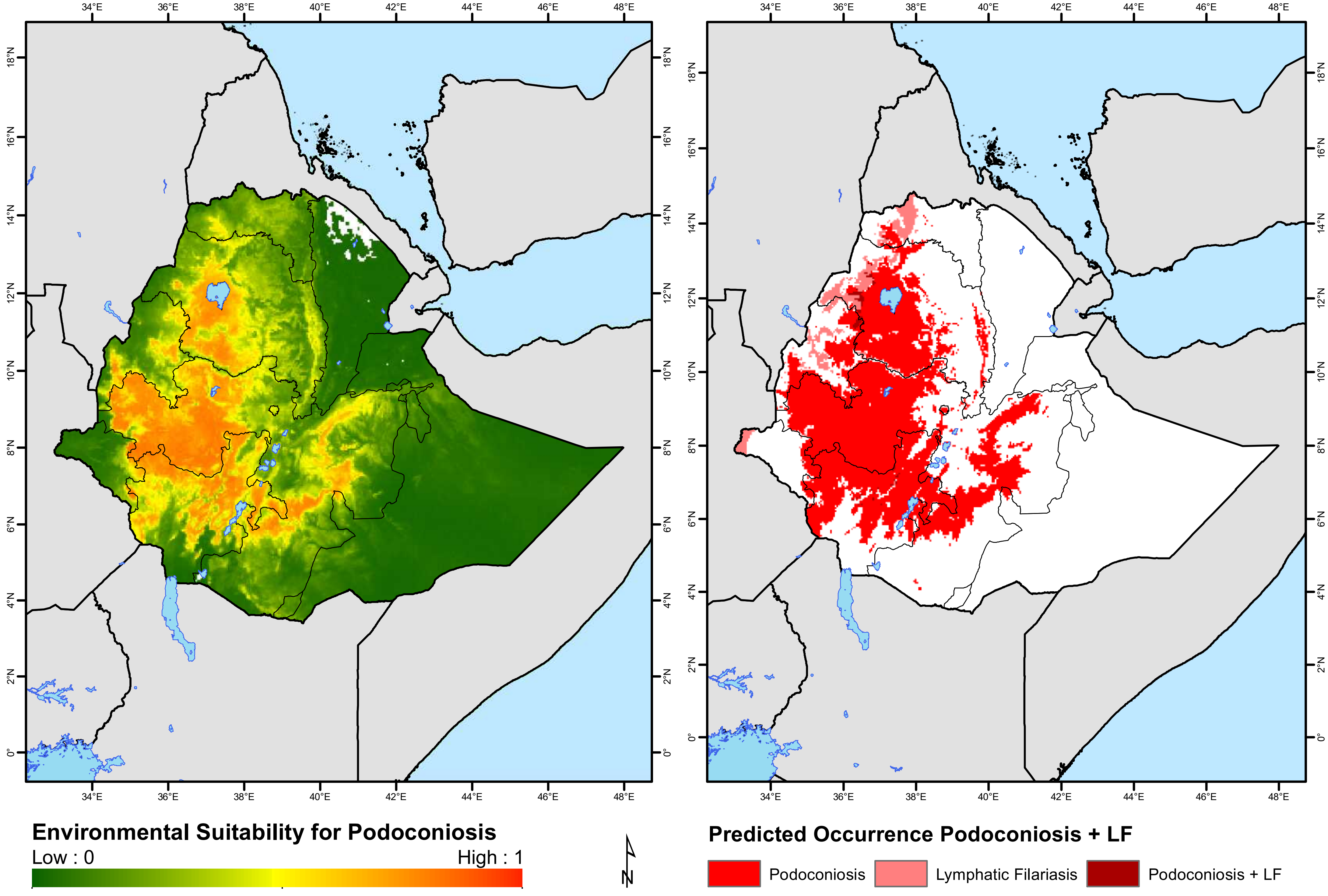

Environmental Suitability for Podoconiosis
Low : 0
High : 1
Predicted Occurrence Podoconiosis + LF
Podoconiosis
Lymphatic Filariasis
Podoconiosis + LF
N

### Gabon

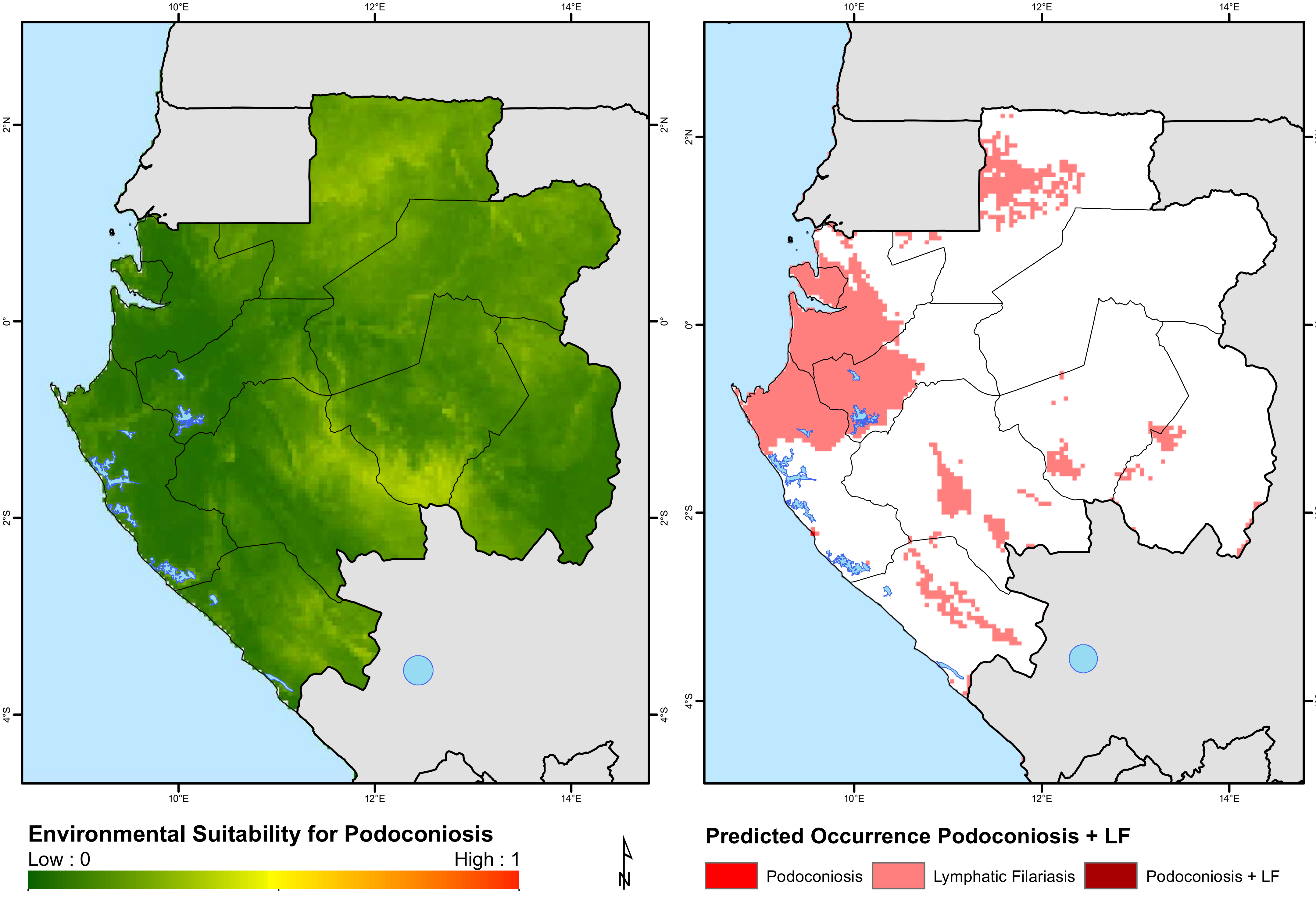

10°E
12°E
14°E
2°N
0°
2°S
4°S
Environmental Suitability for Podoconiosis
Low : 0
High : 1
N
Predicted Occurrence Podoconiosis + LF
Podoconiosis
Lymphatic Filariasis
Podoconiosis + LF

### Gambia

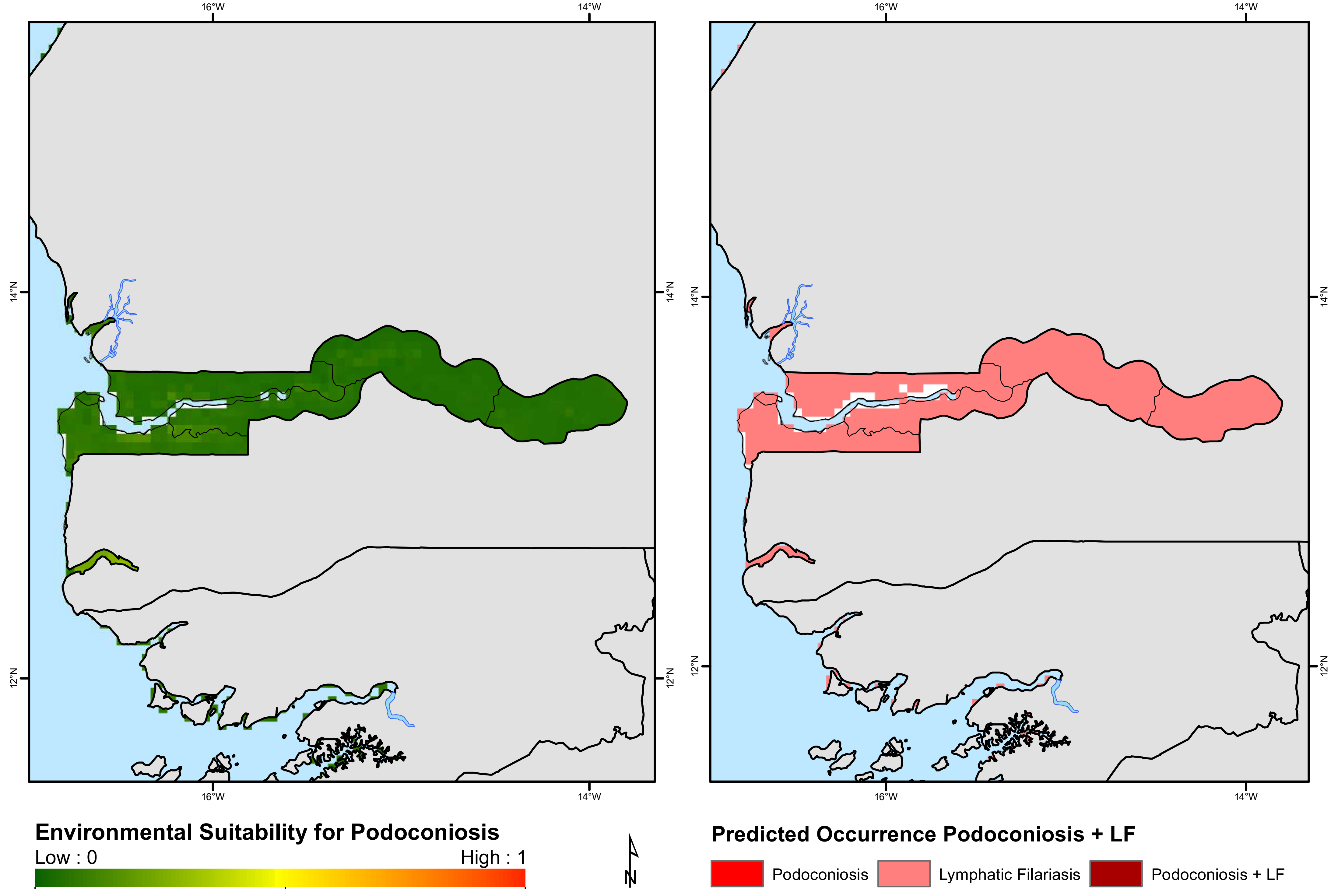
16°W
14°W
14°N
12°N
Environmental Suitability for Podoconiosis
Low : 0
High : 1
N
Predicted Occurrence Podoconiosis + LF
Podoconiosis
Lymphatic Filariasis
Podoconiosis + LF

### Ghana

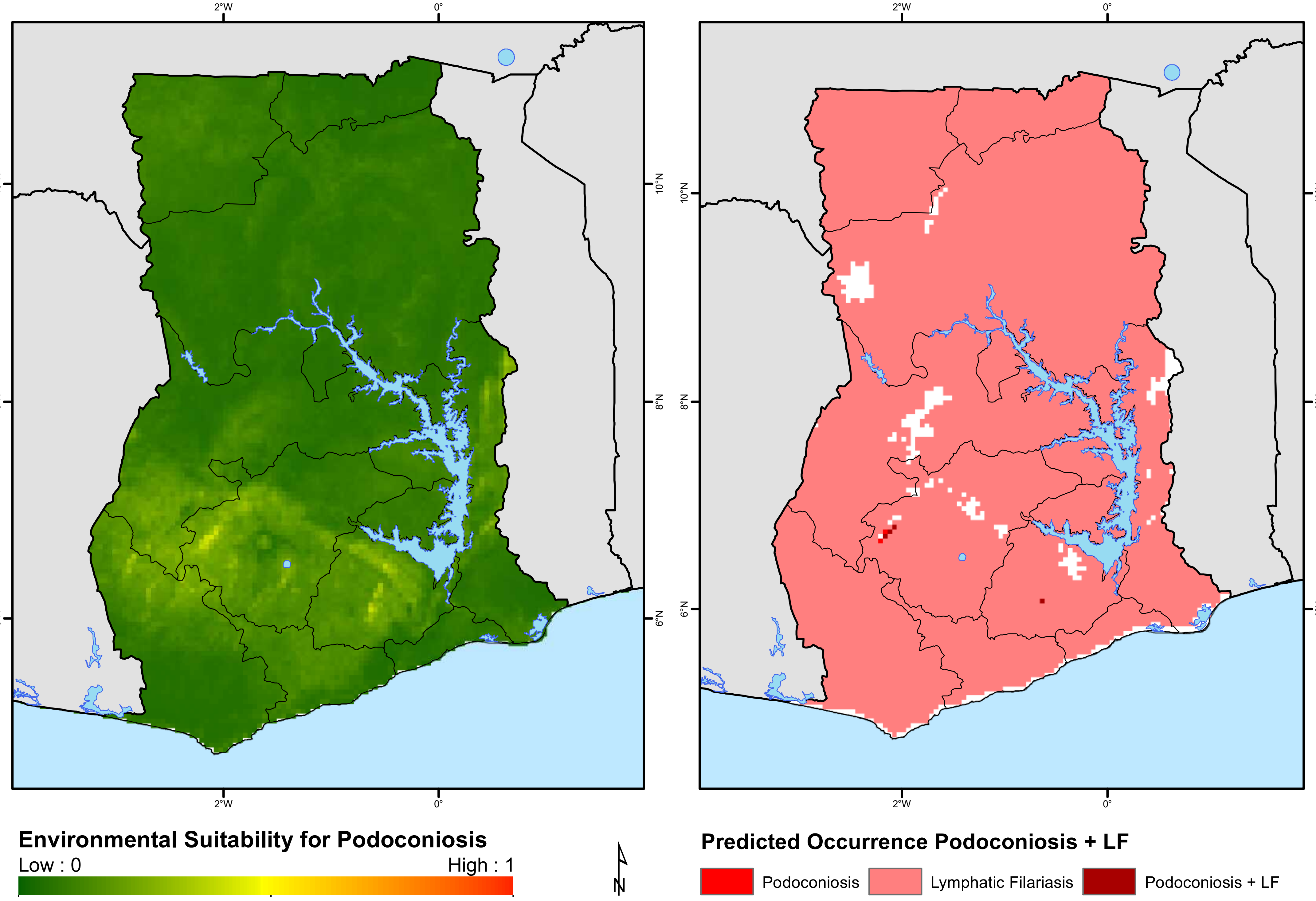
2°W
0°
10°N
8°N
6°N
Environmental Suitability for Podoconiosis
Low : 0
High : 1
N
Predicted Occurrence Podoconiosis + LF
Podoconiosis
Lymphatic Filariasis
Podoconiosis + LF

### Guinea

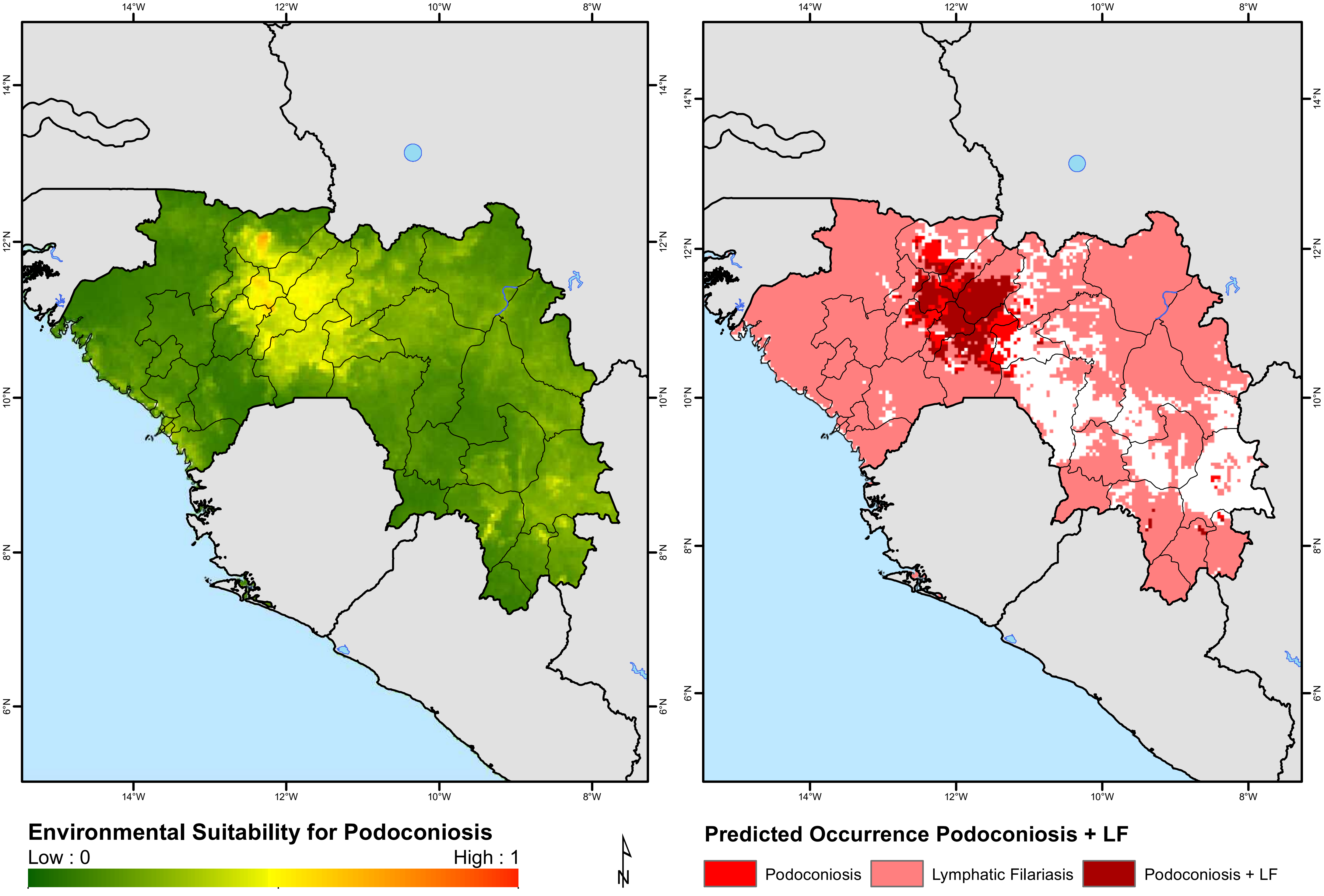
14°W
12°W
10°W
8°W
14°N
12°N
10°N
8°N
6°N
Environmental Suitability for Podoconiosis
Low : 0
High : 1
N
Predicted Occurrence Podoconiosis + LF
Podoconiosis
Lymphatic Filariasis
Podoconiosis + LF

### Guinea-Bissau

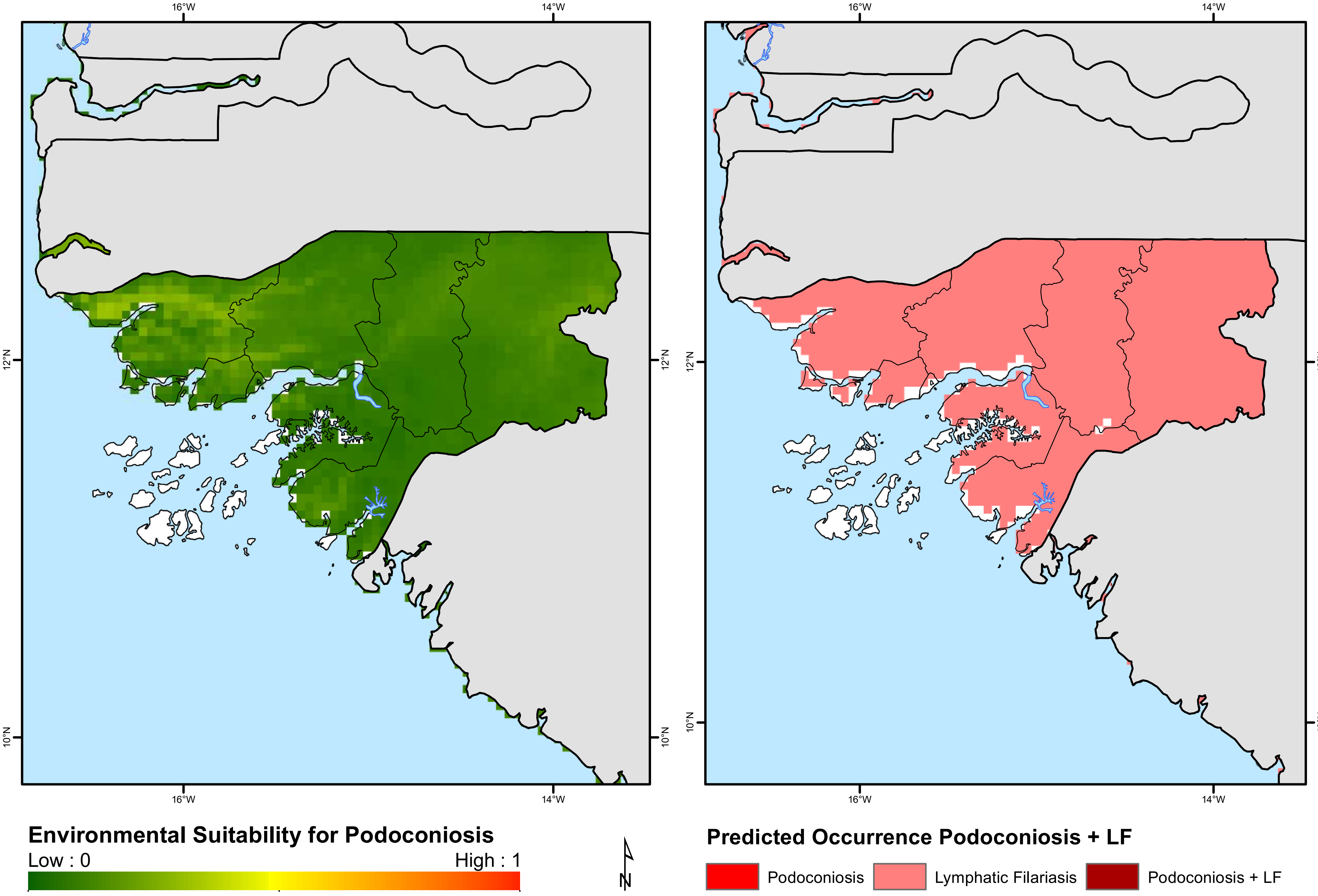

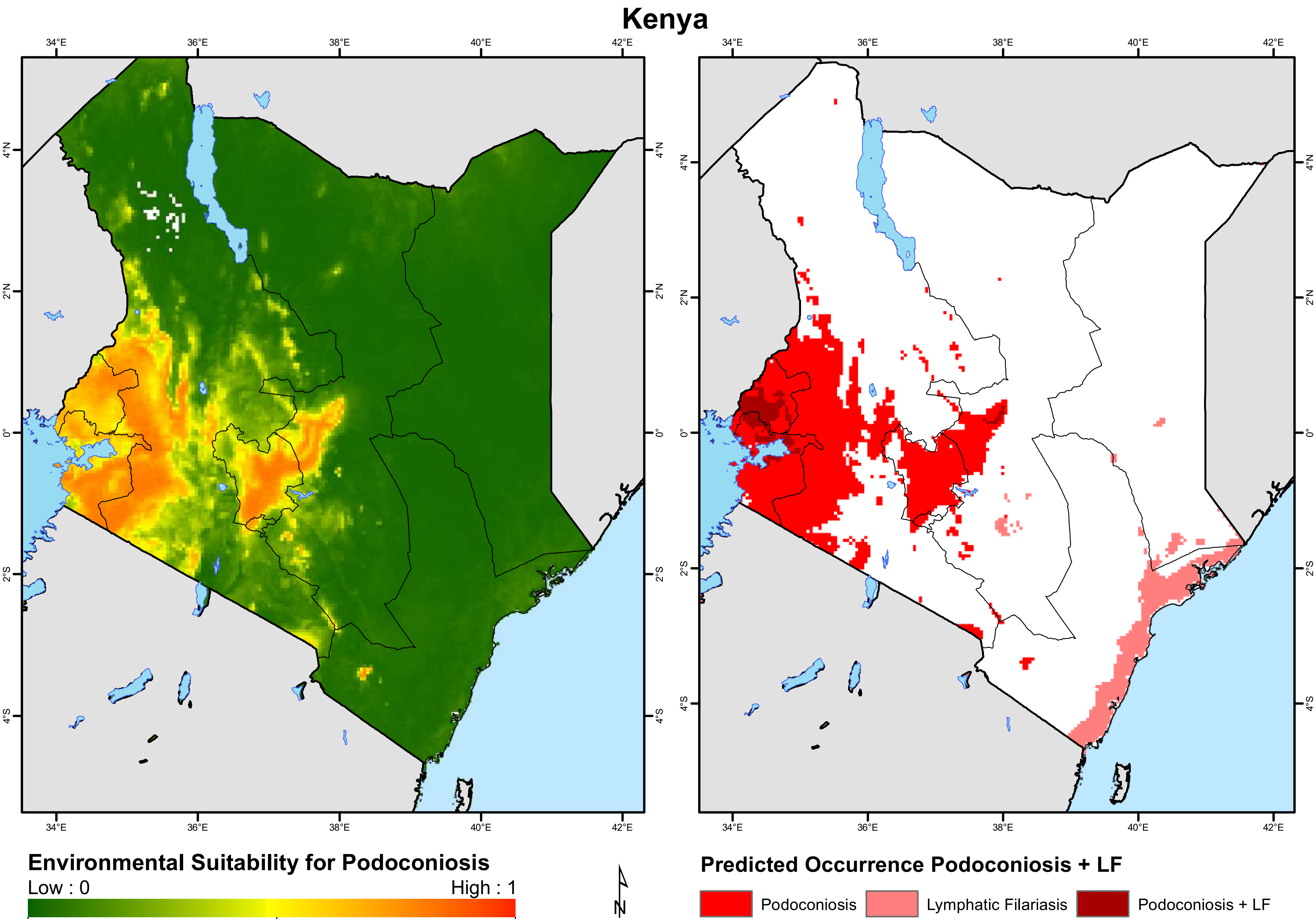
Kenya
34°E
36°E
38°E
40°E
42°E
4°N
2°N
0°
2°S
4°S
Environmental Suitability for Podoconiosis
Low : 0
High : 1
N
Predicted Occurrence Podoconiosis + LF
Podoconiosis
Lymphatic Filariasis
Podoconiosis + LF

### Lesotho

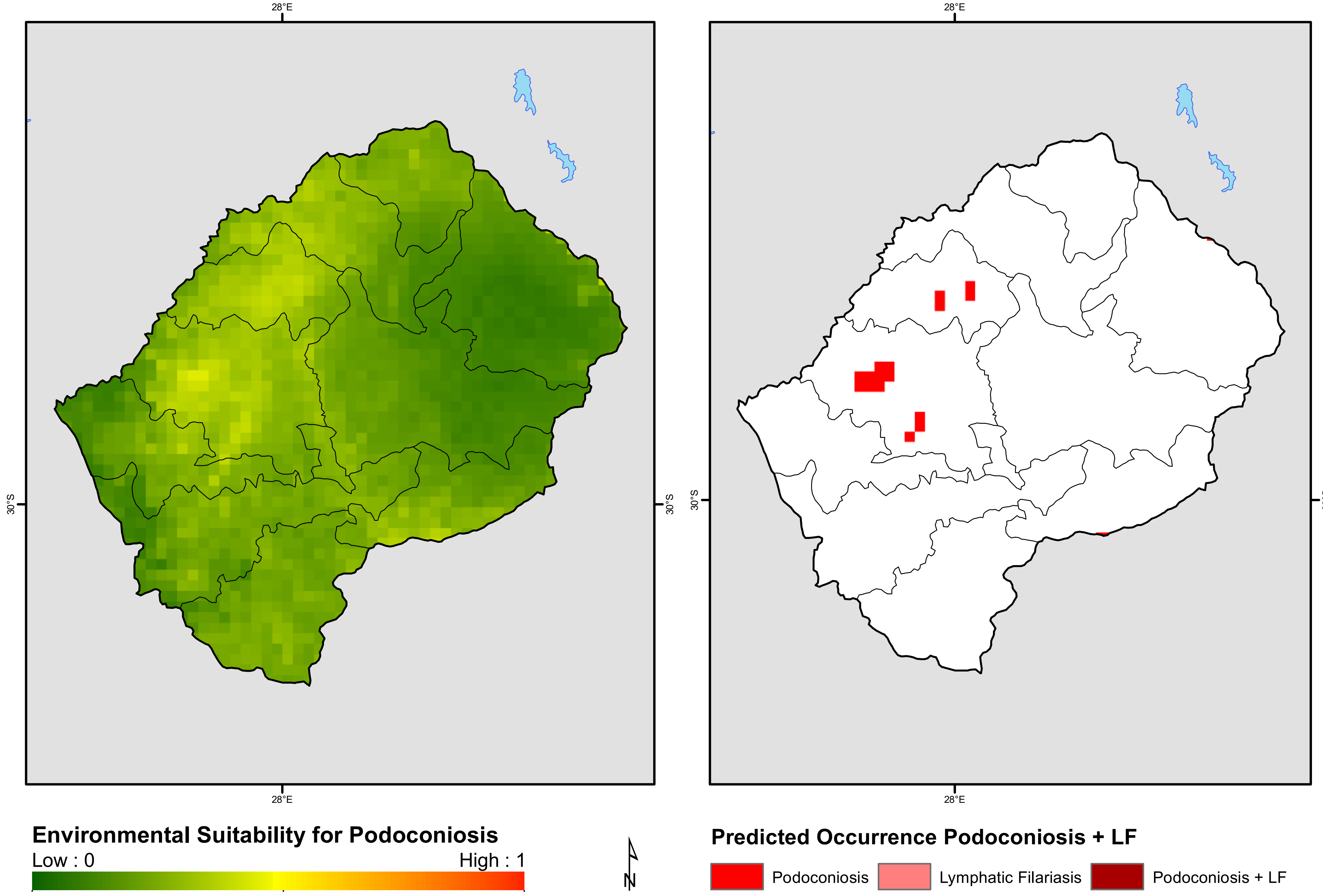

28°E
30°S
Environmental Suitability for Podoconiosis
Low : 0
High : 1
N
Predicted Occurrence Podoconiosis + LF
Podoconiosis
Lymphatic Filariasis
Podoconiosis + LF

### Liberia

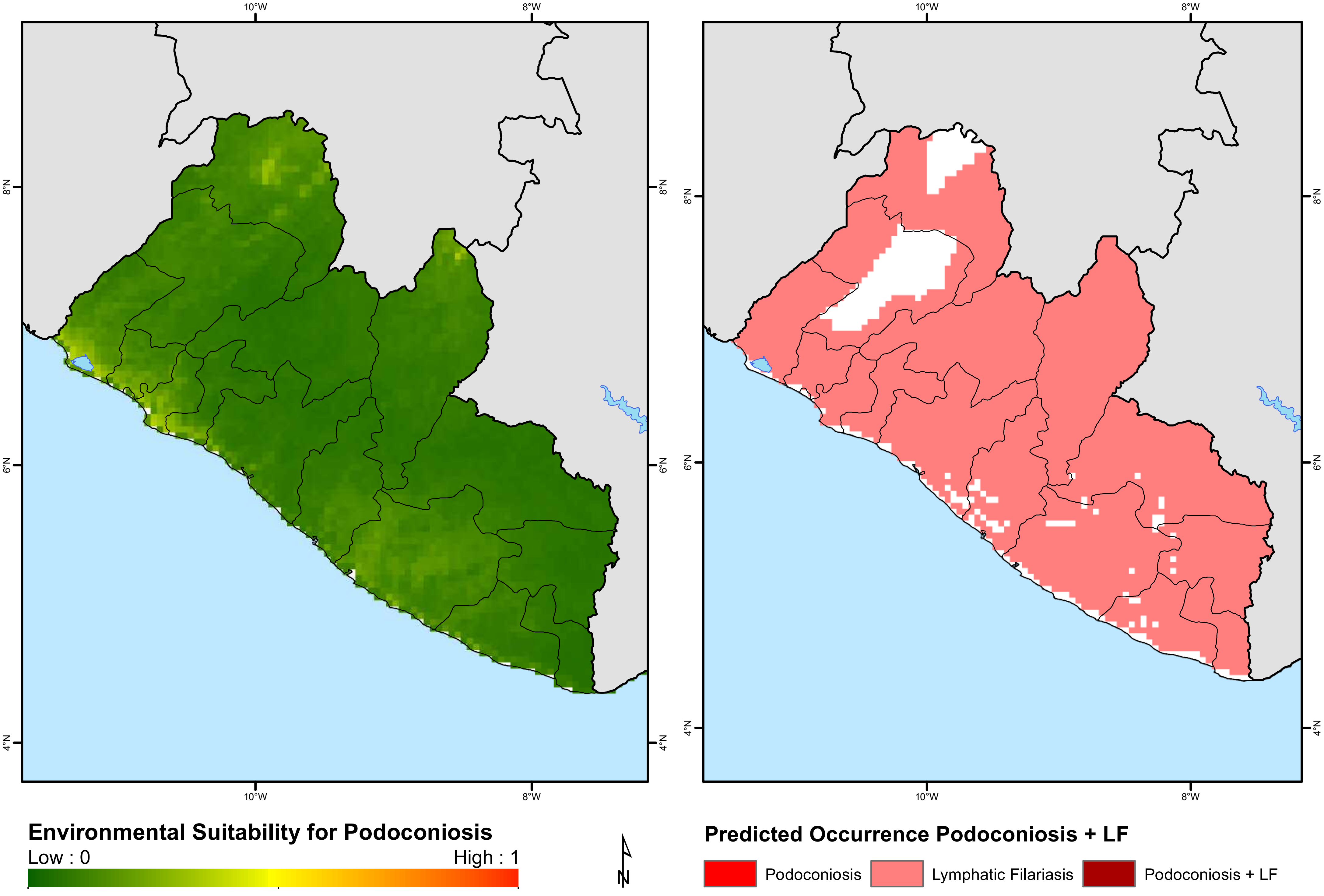

10°W
8°W
8°N
6°N
4°N
Environmental Suitability for Podoconiosis
Low : 0
High : 1
N
Predicted Occurrence Podoconiosis + LF
Podoconiosis
Lymphatic Filariasis
Podoconiosis + LF

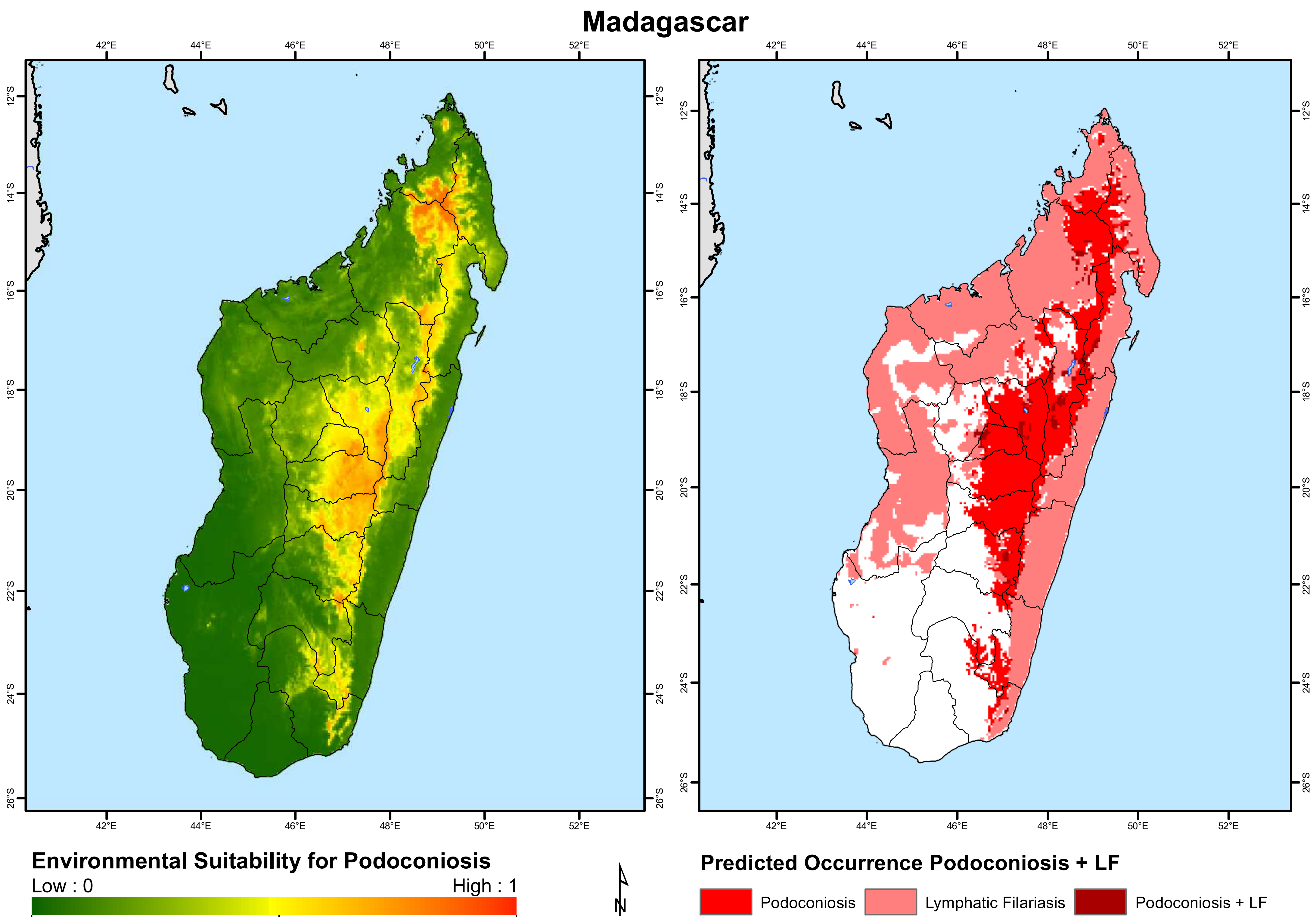

Madagascar
42°E
44°E
46°E
48°E
50°E
52°E
12°S
14°S
16°S
18°S
20°S
22°S
24°S
26°S
Environmental Suitability for Podoconiosis
Low : 0
High : 1
N
Predicted Occurrence Podoconiosis + LF
Podoconiosis
Lymphatic Filariasis
Podoconiosis + LF

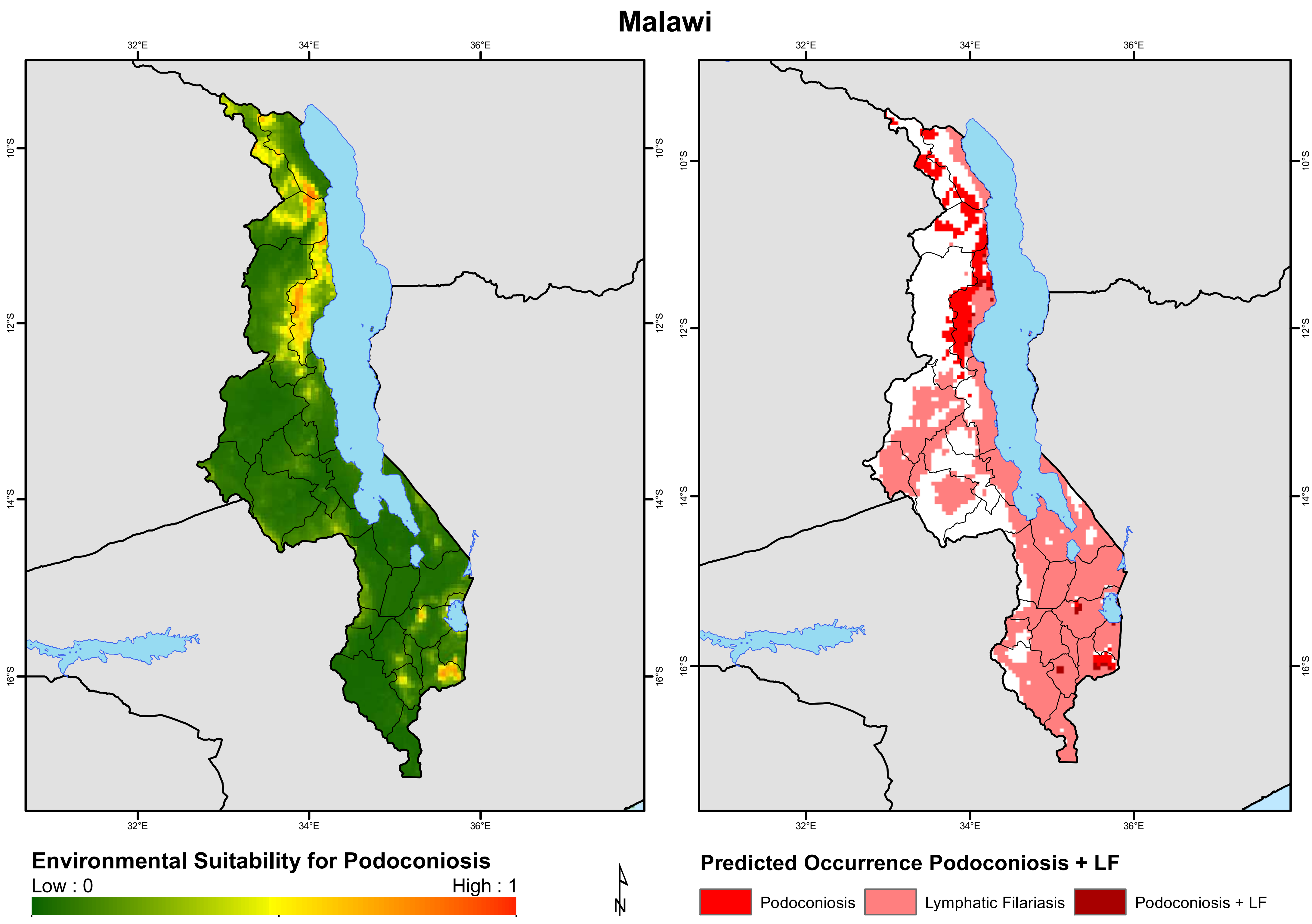
Malawi
32°E
34°E
36°E
10°S
12°S
14°S
16°S
Environmental Suitability for Podoconiosis
Low : 0
High : 1
N
Predicted Occurrence Podoconiosis + LF
Podoconiosis
Lymphatic Filariasis
Podoconiosis + LF

### Mali

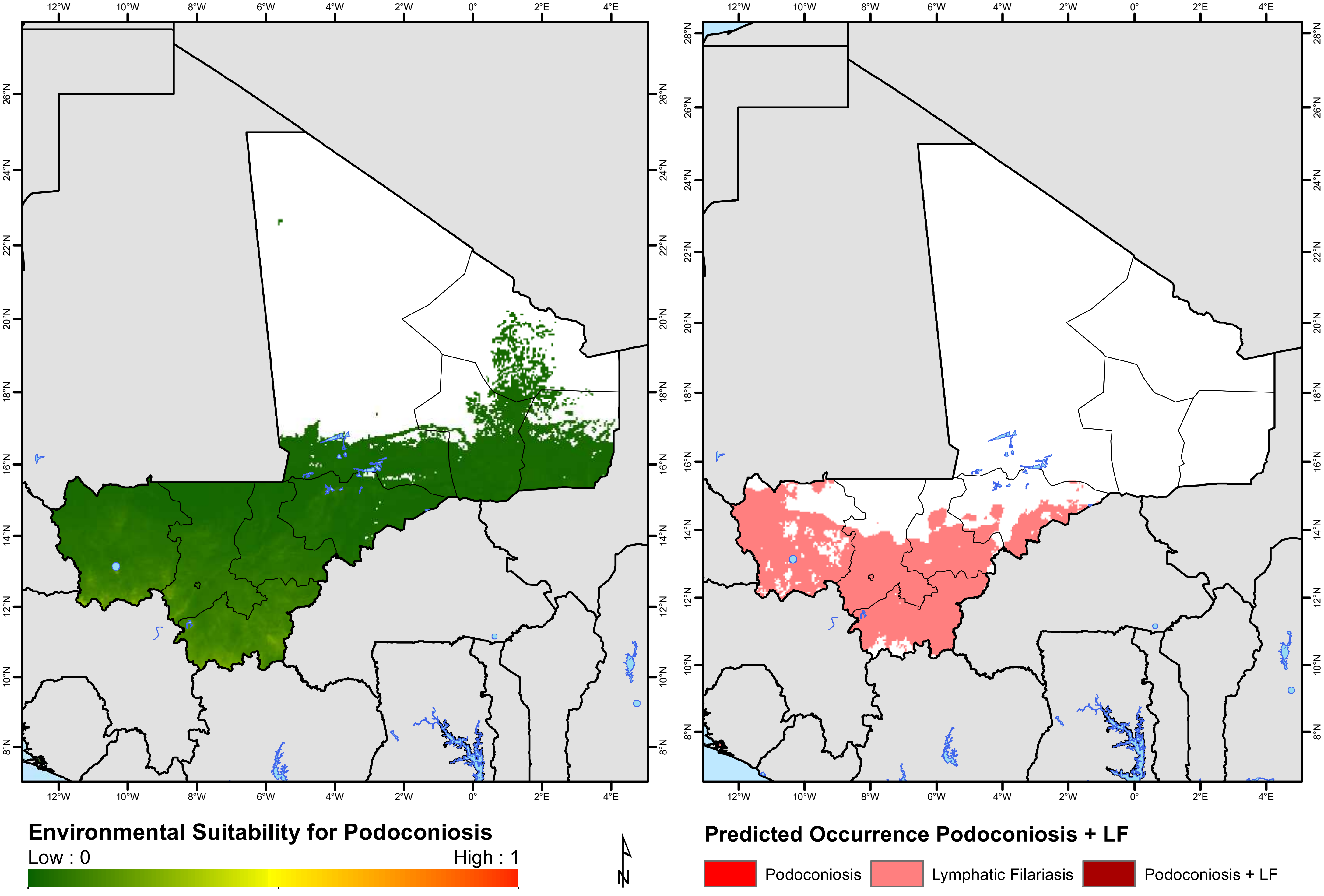
Environmental Suitability for Podoconiosis
Low : 0
High : 1
Predicted Occurrence Podoconiosis + LF
Podoconiosis
Lymphatic Filariasis
Podoconiosis + LF
N

### Mauritania

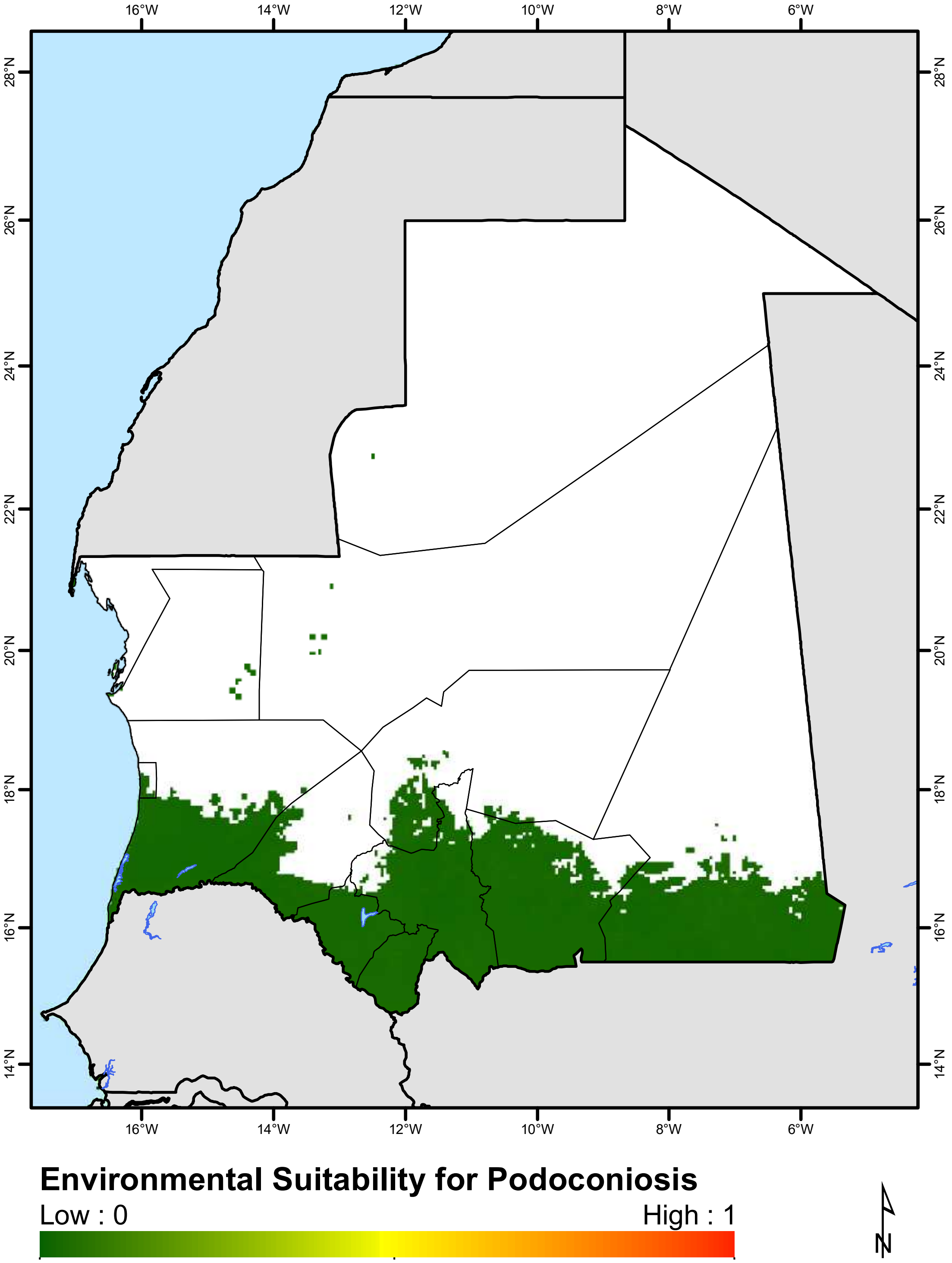
16°W
14°W
12°W
10°W
8°W
6°W
28°N
26°N
24°N
22°N
20°N
18°N
16°N
14°N
Environmental Suitability for Podoconiosis
Low : 0
High : 1
N

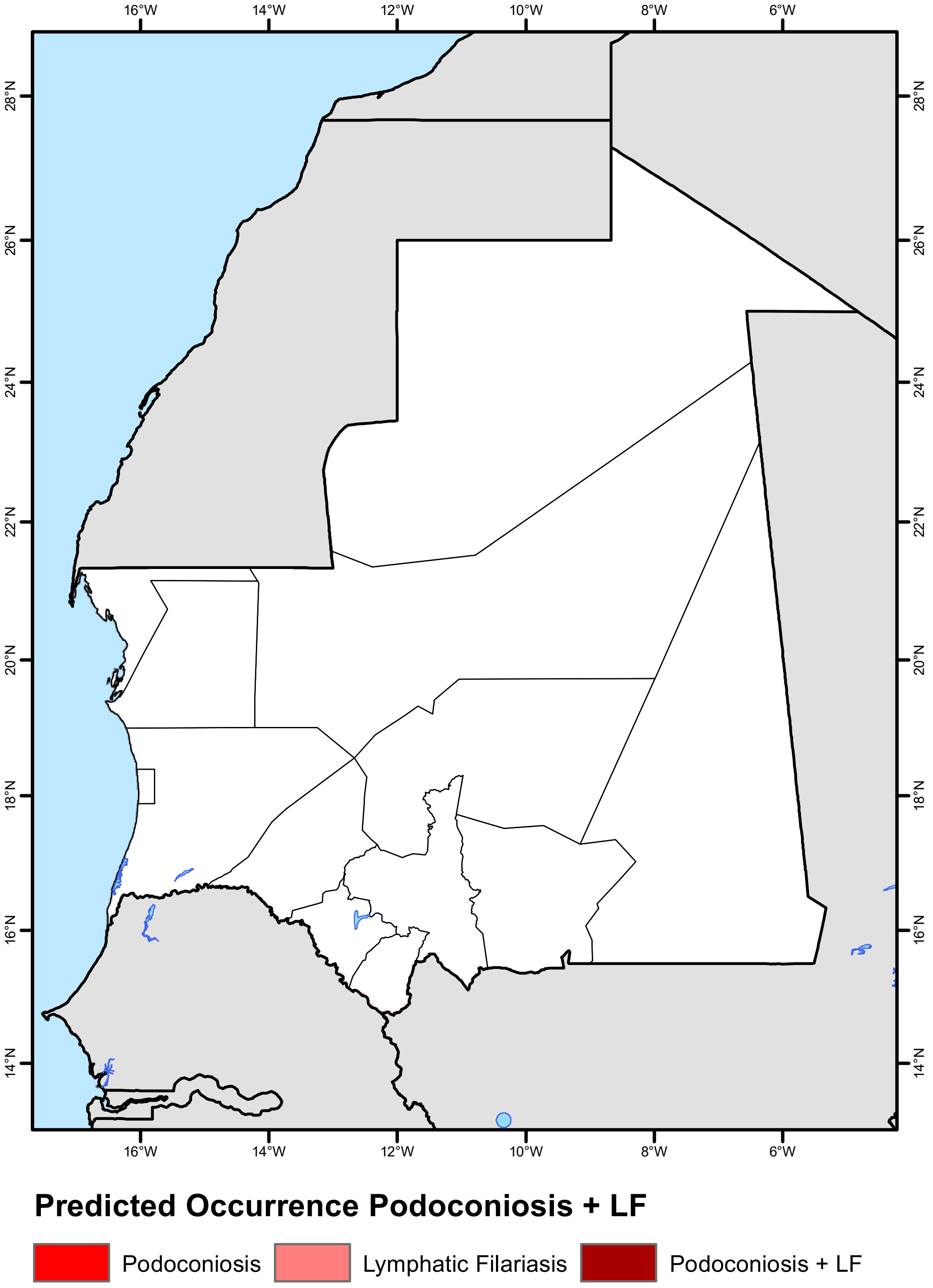
16°W
14°W
12°W
10°W
8°W
6°W
28°N
26°N
24°N
22°N
20°N
18°N
16°N
14°N
Predicted Occurrence Podoconiosis + LF
Podoconiosis
Lymphatic Filariasis
Podoconiosis + LF

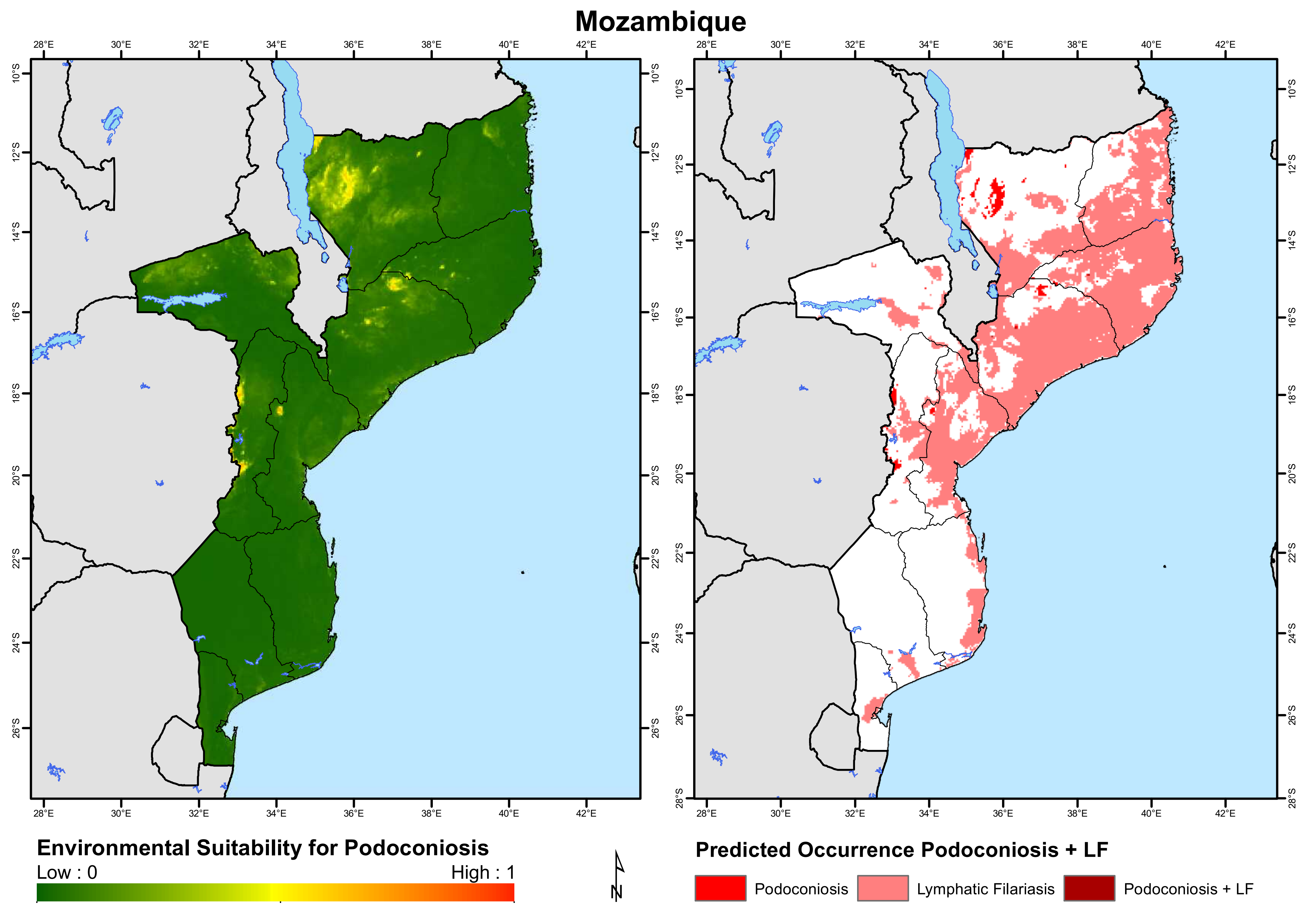

Mozambique
Environmental Suitability for Podoconiosis
Low : 0
High : 1
N
Predicted Occurrence Podoconiosis + LF
Podoconiosis
Lymphatic Filariasis
Podoconiosis + LF

### Namibia

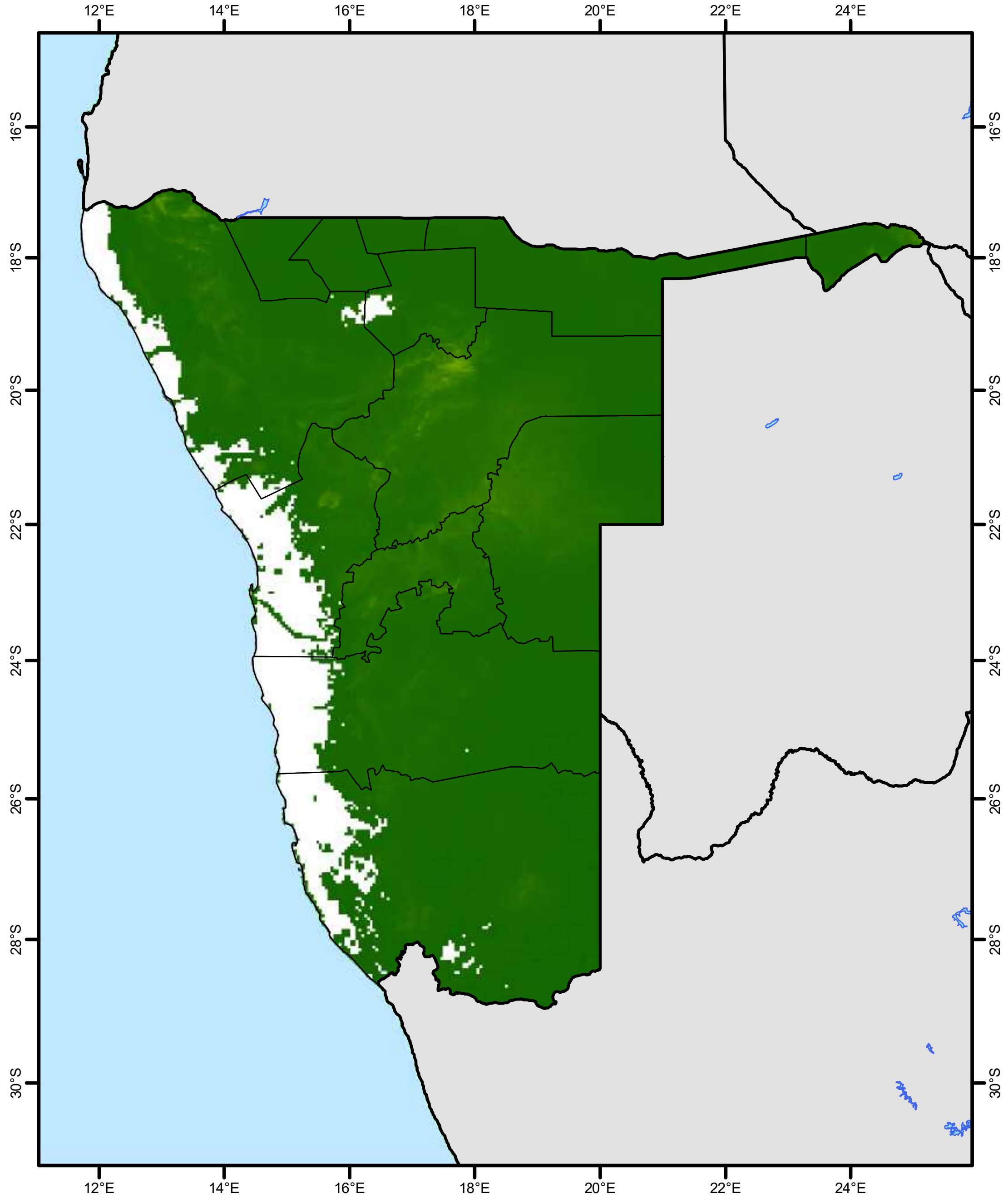


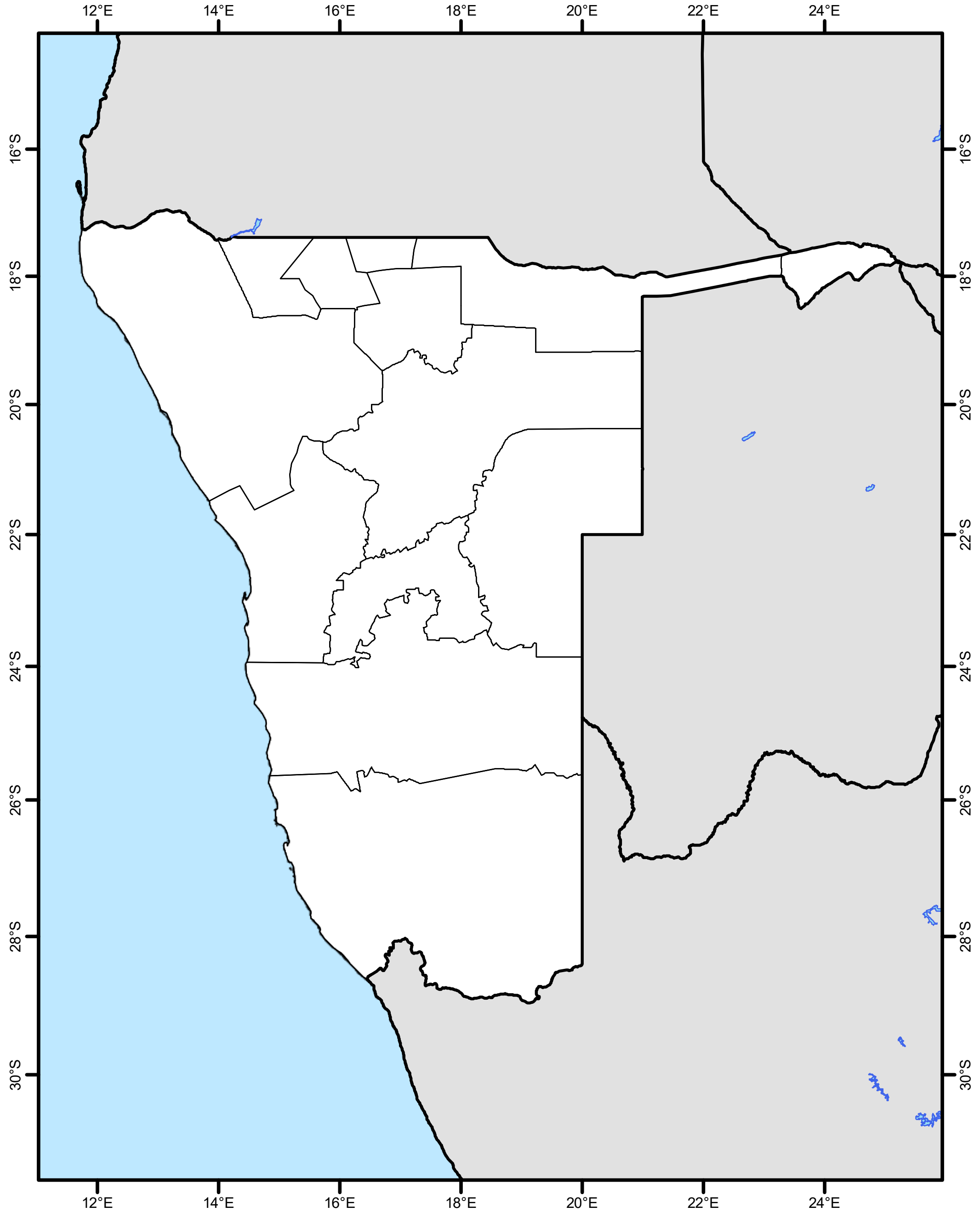


**Environmental Suitability for Podoconiosis**

Low : 0 High : 1

**Predicted Occurrence Podoconiosis + LF**

Podoconiosis
Lymphatic Filariasis
Podoconiosis + LF

### Niger

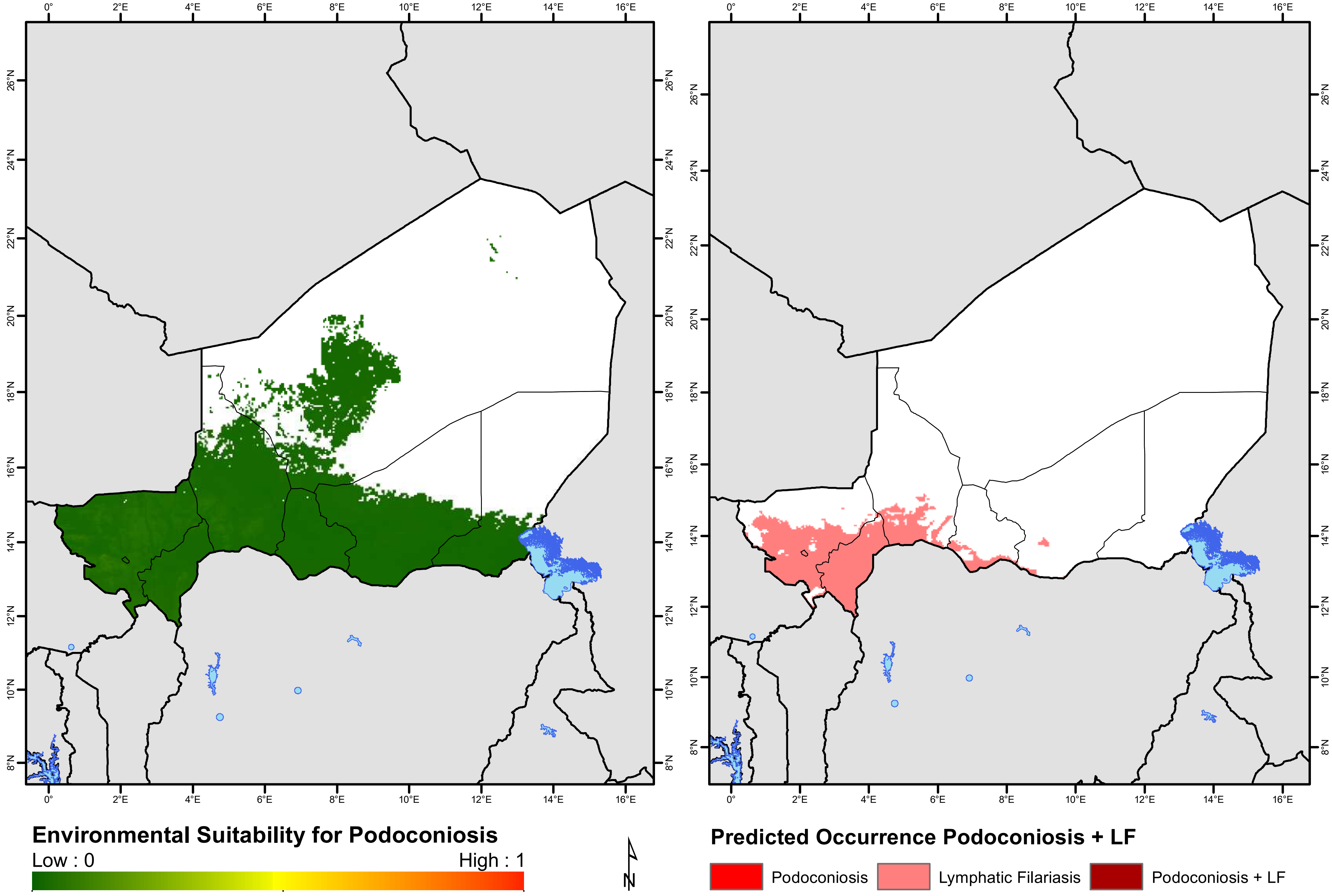
Environmental Suitability for Podoconiosis
Low : 0
High : 1
Predicted Occurrence Podoconiosis + LF
Podoconiosis
Lymphatic Filariasis
Podoconiosis + LF
N

### Nigeria

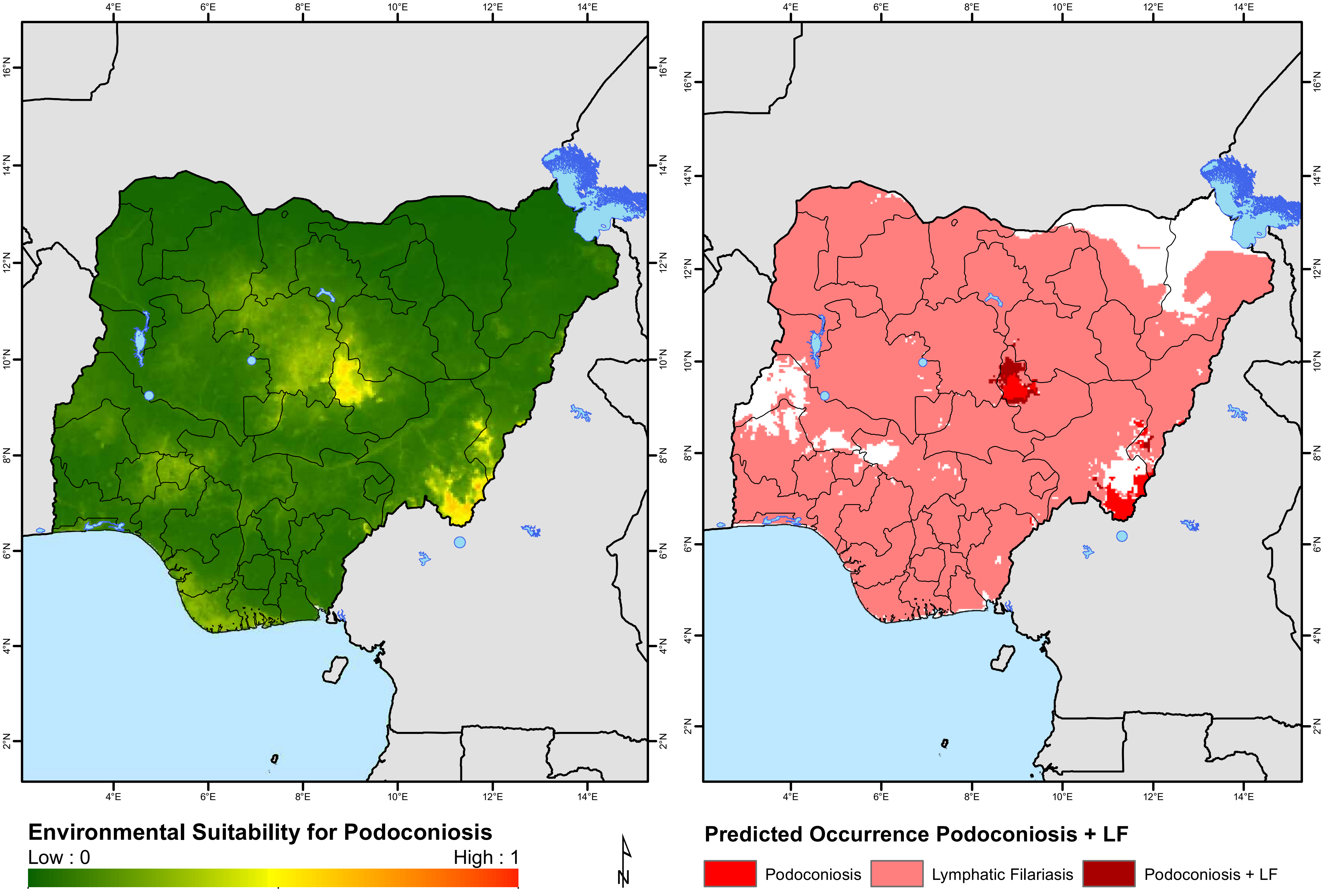
4°E
6°E
8°E
10°E
12°E
14°E
2°N
4°N
6°N
8°N
10°N
12°N
14°N
16°N
Environmental Suitability for Podoconiosis
Low : 0
High : 1
N
Predicted Occurrence Podoconiosis + LF
Podoconiosis
Lymphatic Filariasis
Podoconiosis + LF

### Rwanda

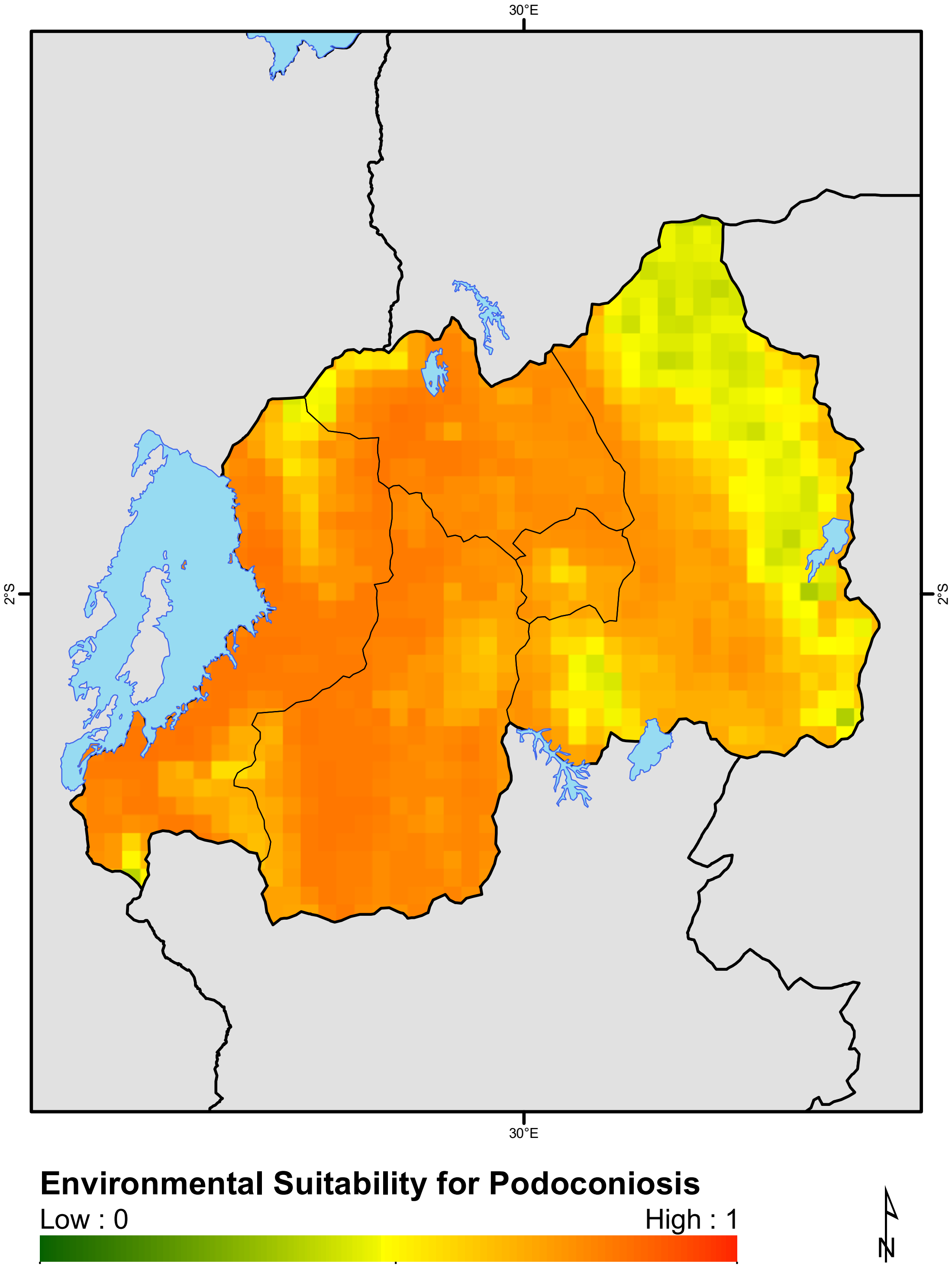
30°E
2°S
2°S
30°E
Environmental Suitability for Podoconiosis
Low : 0
High : 1

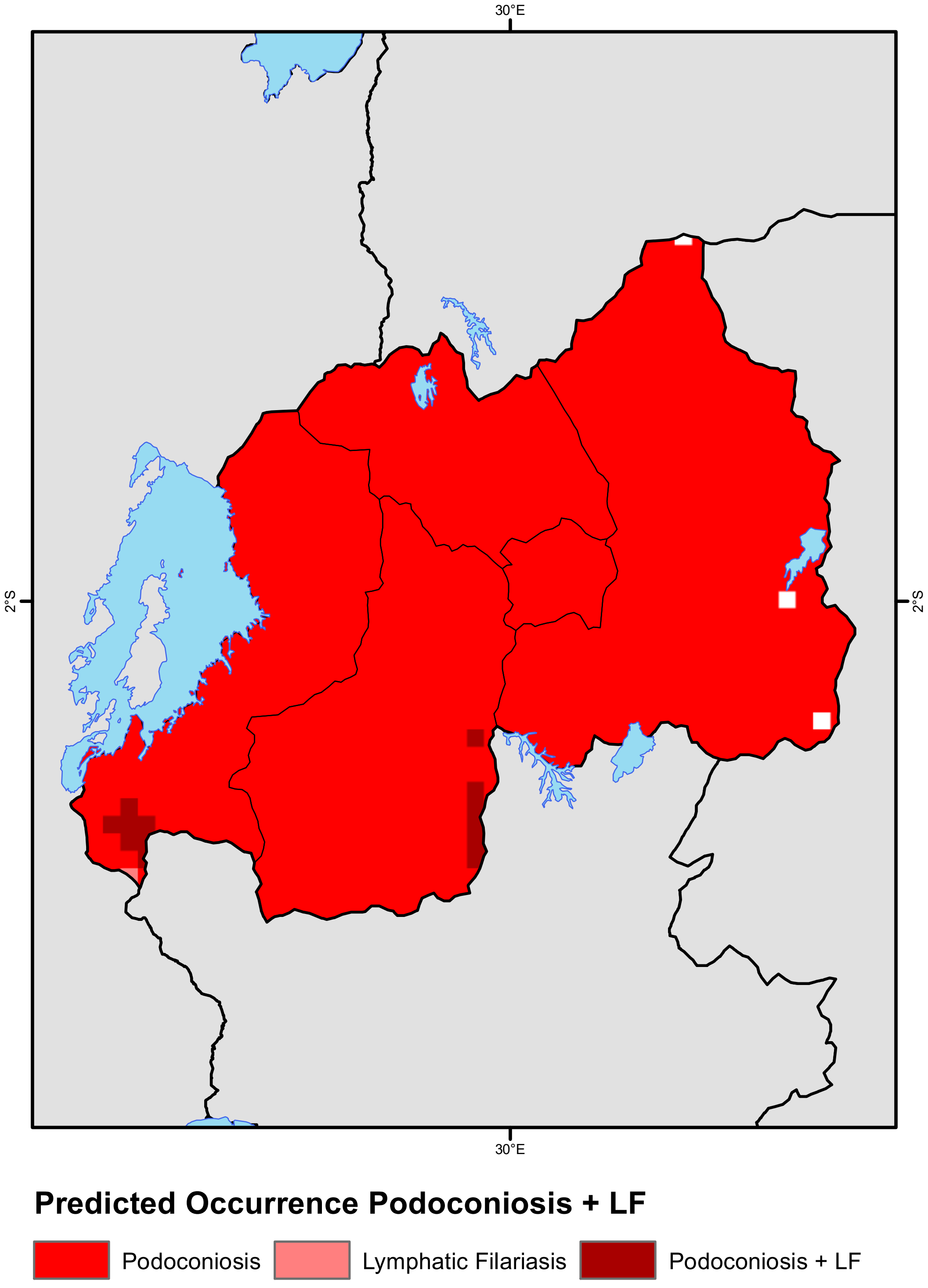
30°E
2°S
2°S
30°E
Predicted Occurrence Podoconiosis + LF
Podoconiosis
Lymphatic Filariasis
Podoconiosis + LF

#### Sao Tome and Principe

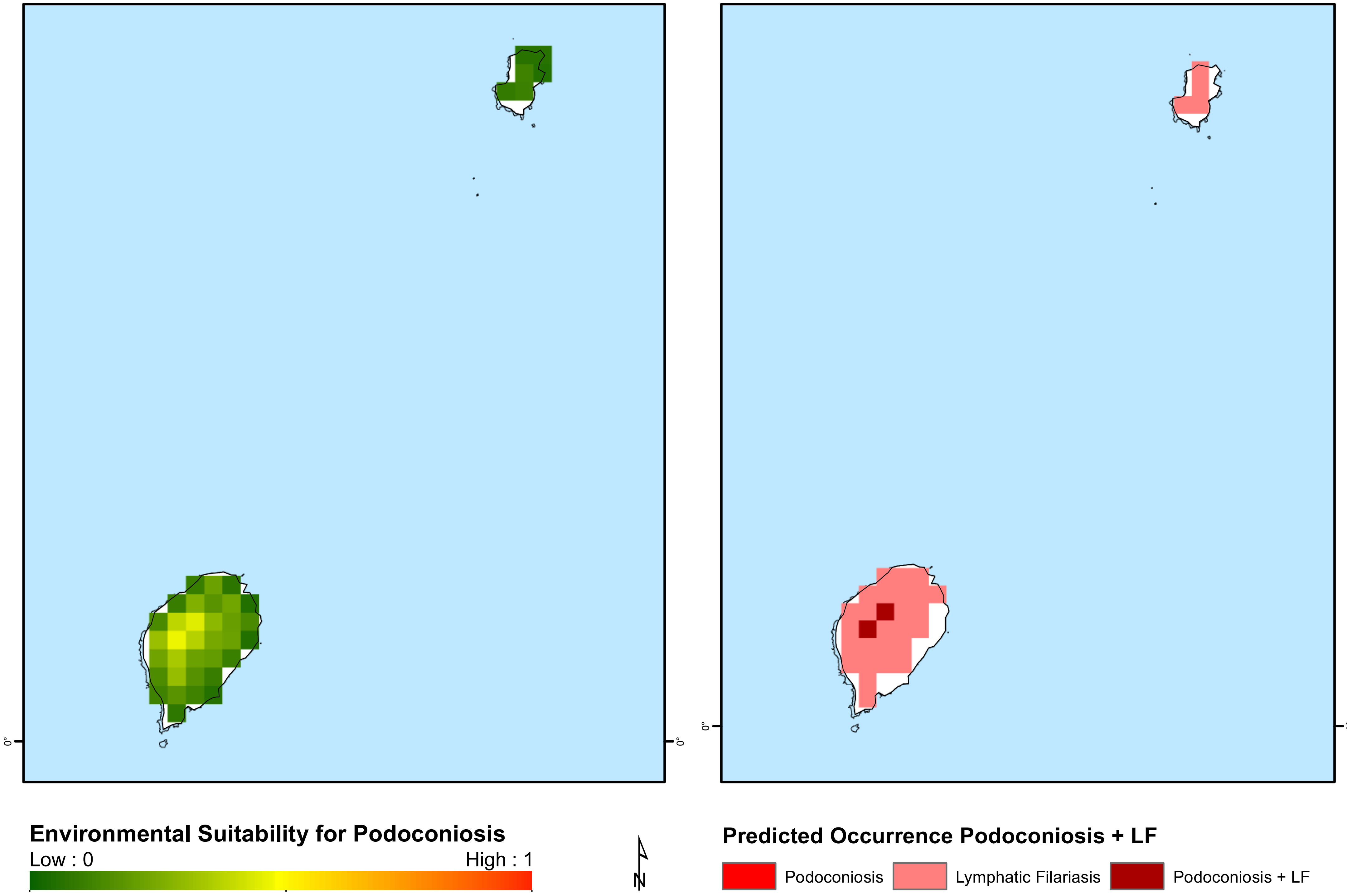

### Senegal

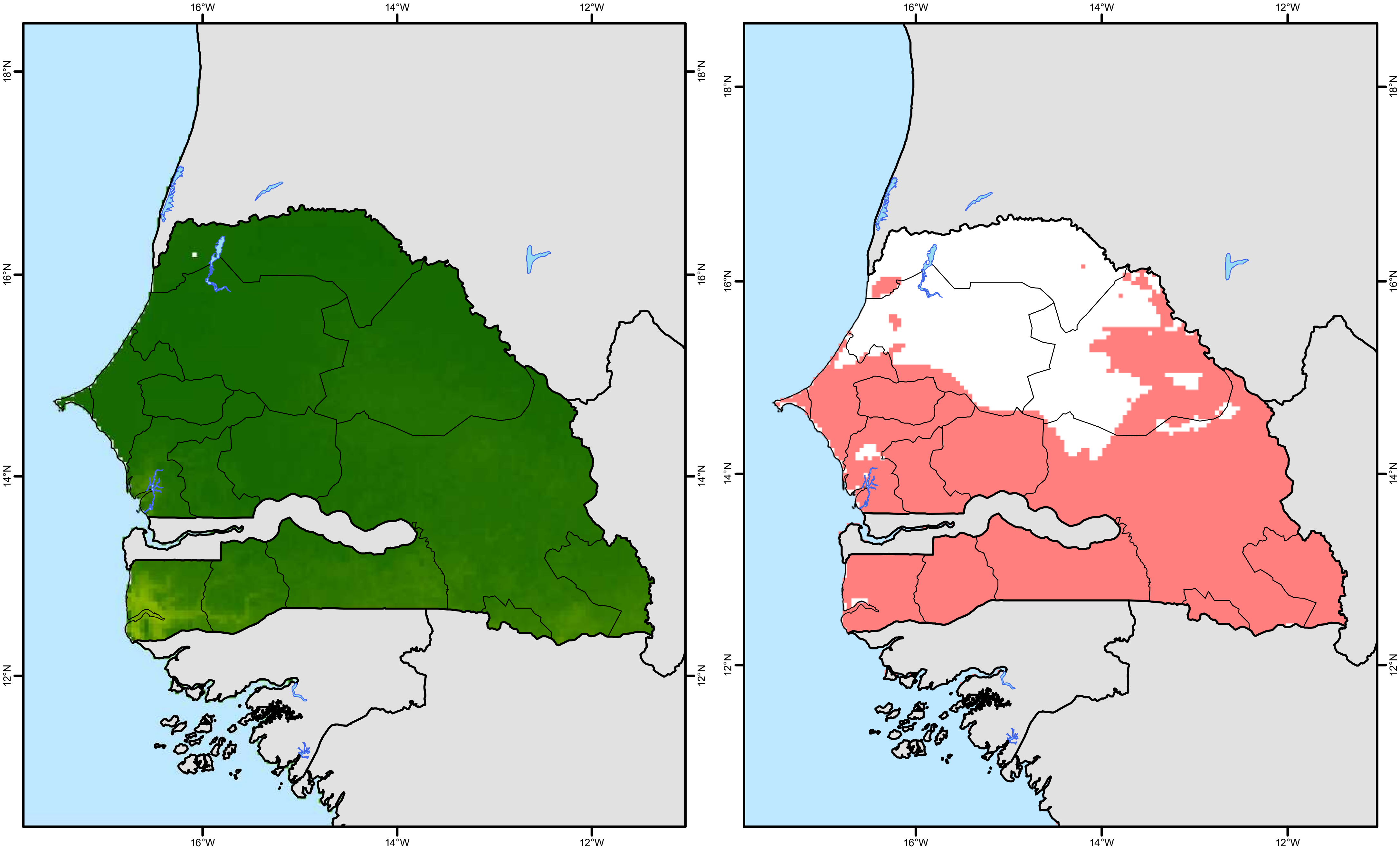


**Environmental Suitability for Podoconiosis**

Low : 0 High : 1

**Predicted Occurrence Podoconiosis + LF**

Podoconiosis
Lymphatic Filariasis
Podoconiosis + LF

### Sierra Leone

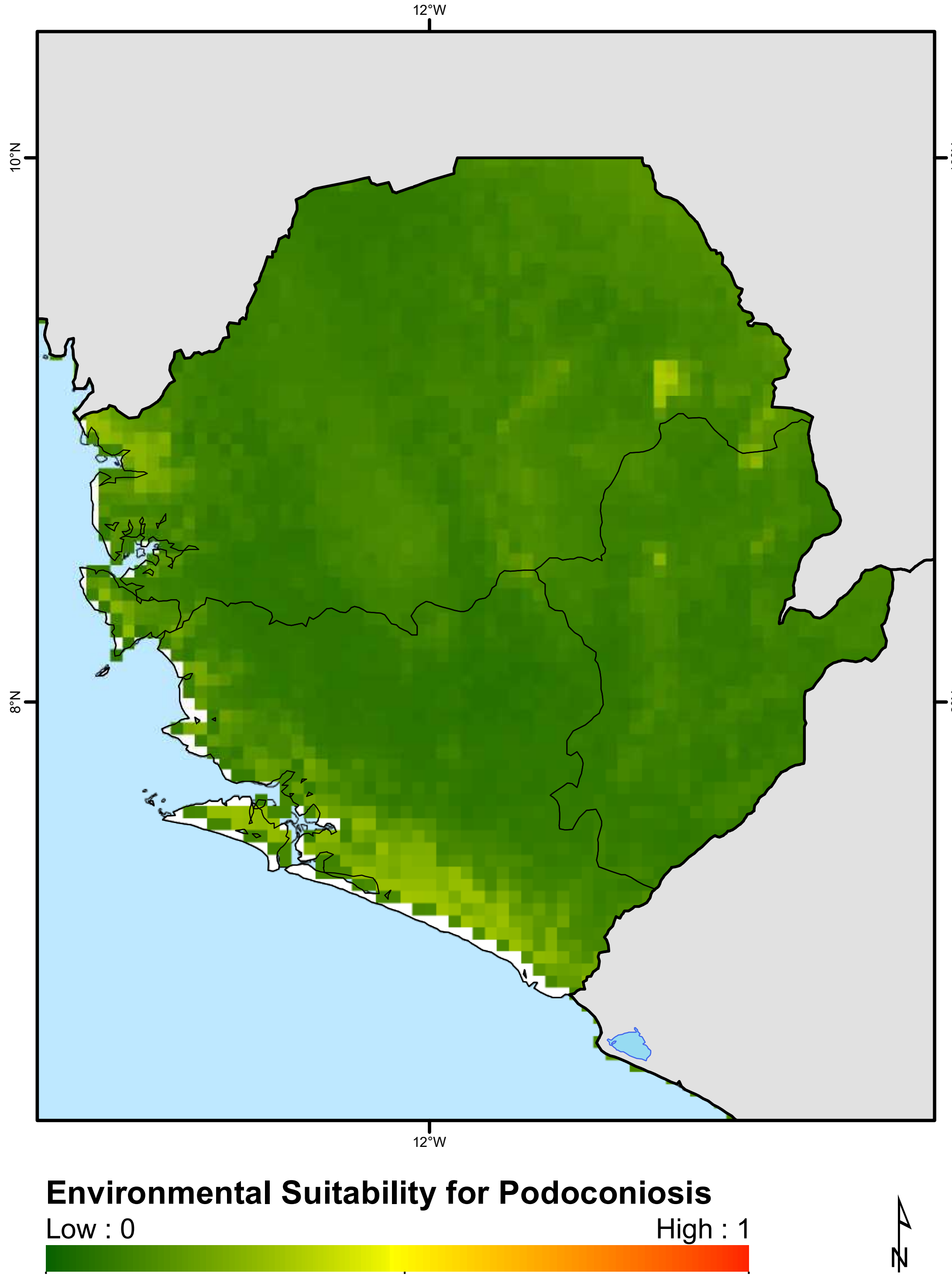


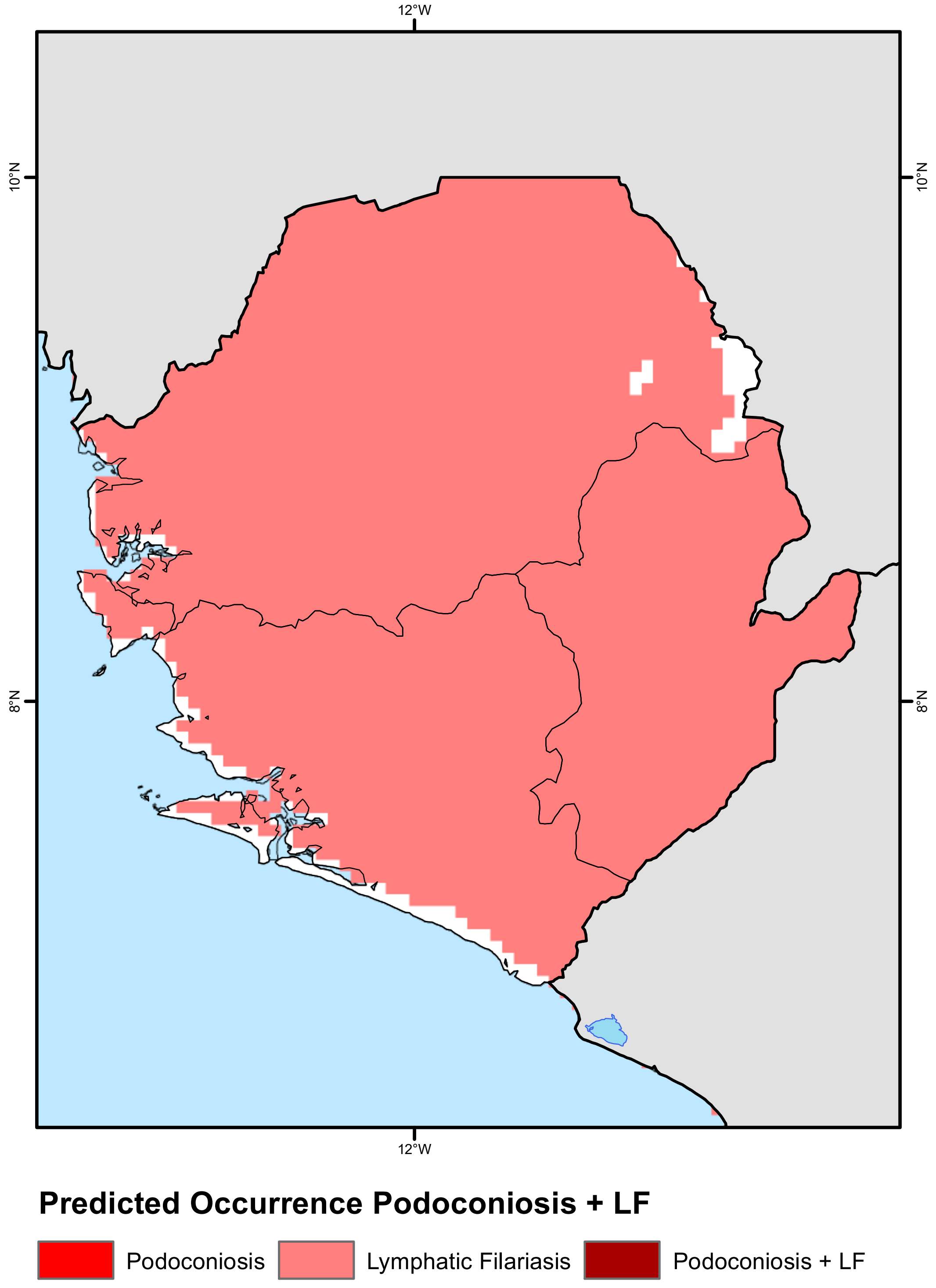

### Somalia

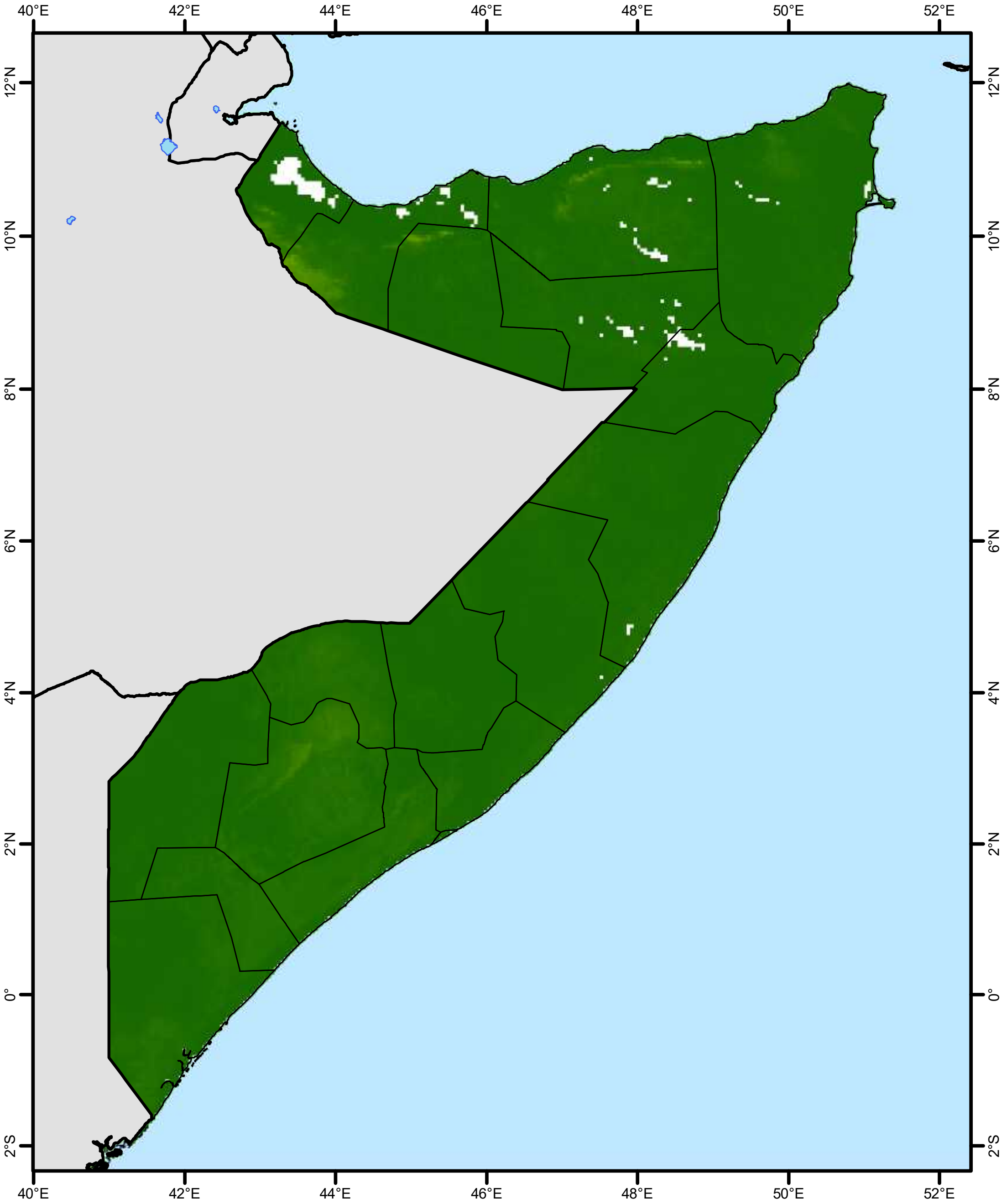


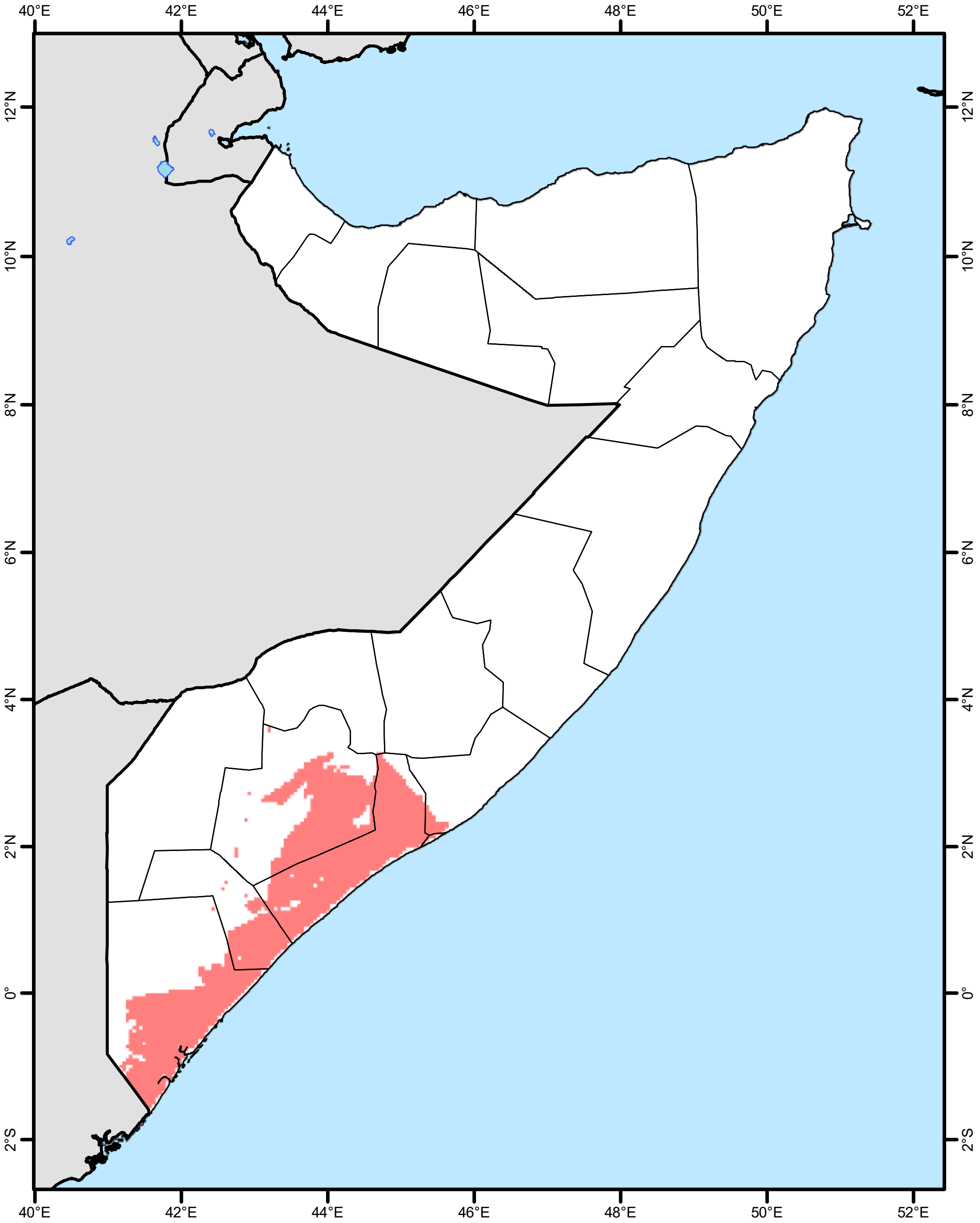


**Environmental Suitability for Podoconiosis**

Low : 0 High : 1

N

**Predicted Occurrence Podoconiosis + LF**

Podoconiosis

Lymphatic Filariasis

Podoconiosis + LF

### South Africa

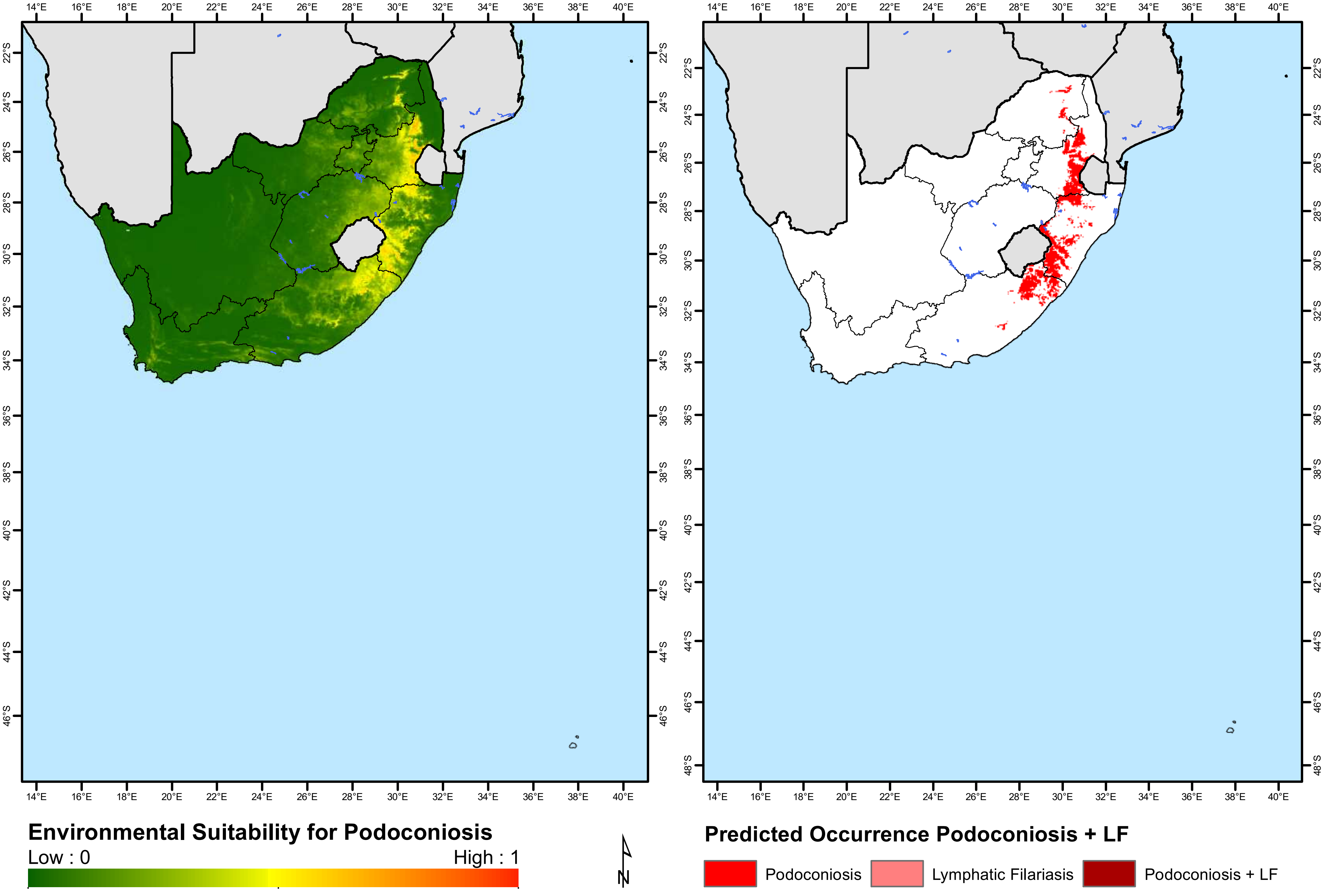

### South Sudan

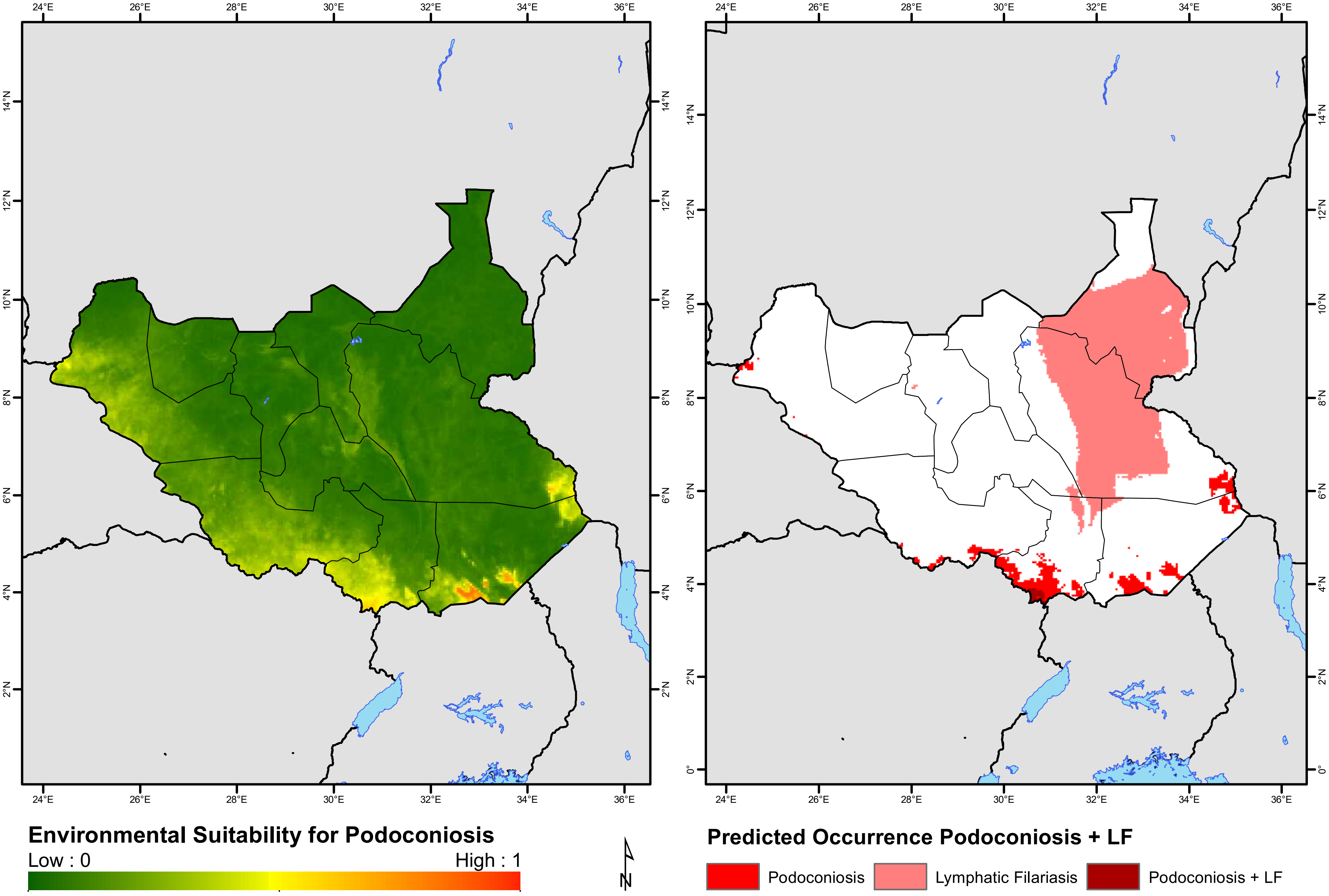

24°E
26°E
28°E
30°E
32°E
34°E
36°E
0°
2°N
4°N
6°N
8°N
10°N
12°N
14°N
Environmental Suitability for Podoconiosis
Low : 0
High : 1
N
Predicted Occurrence Podoconiosis + LF
Podoconiosis
Lymphatic Filariasis
Podoconiosis + LF

### Sudan

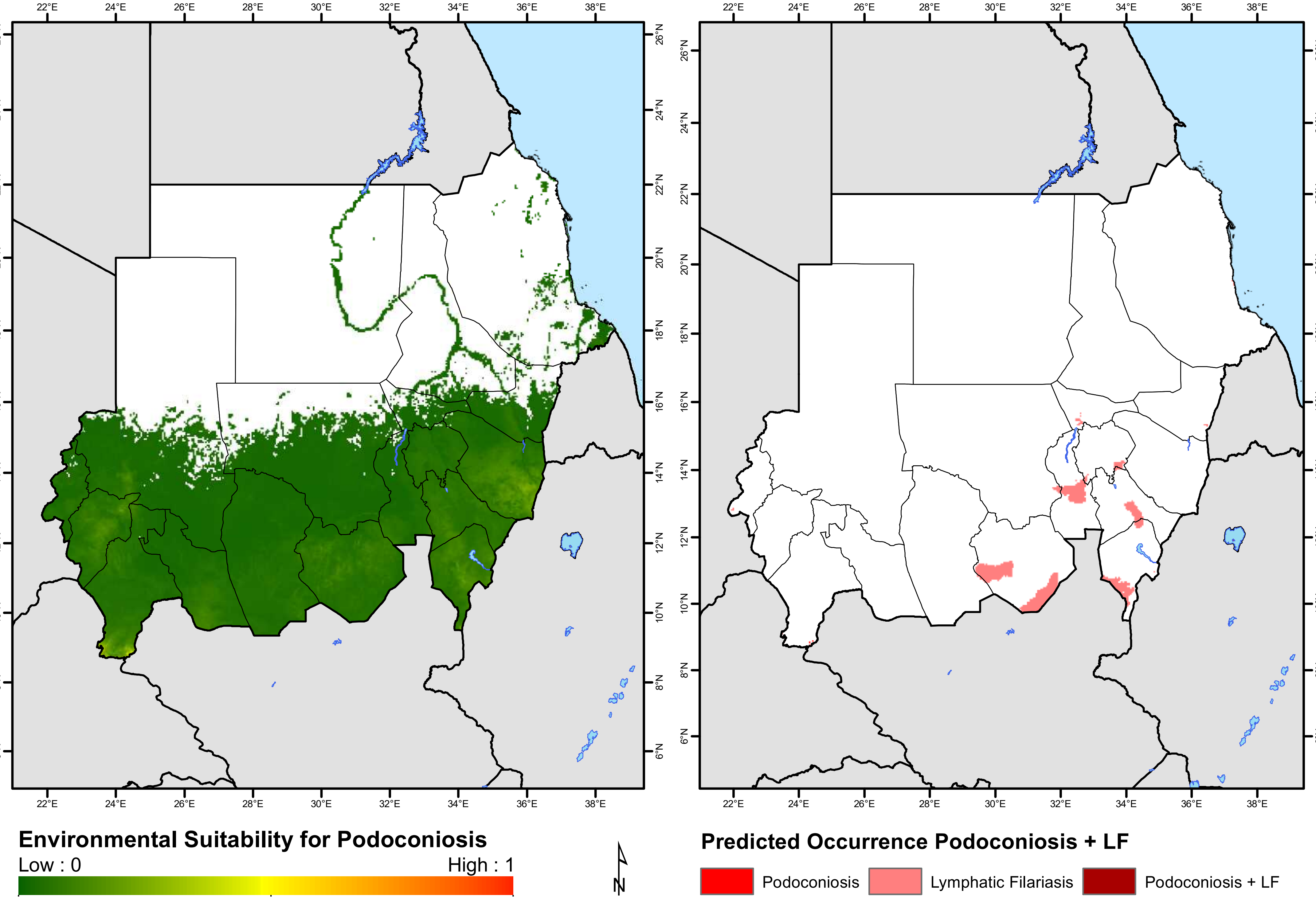

Environmental Suitability for Podoconiosis
Low : 0
High : 1
N
Predicted Occurrence Podoconiosis + LF
Podoconiosis
Lymphatic Filariasis
Podoconiosis + LF

### Swaziland

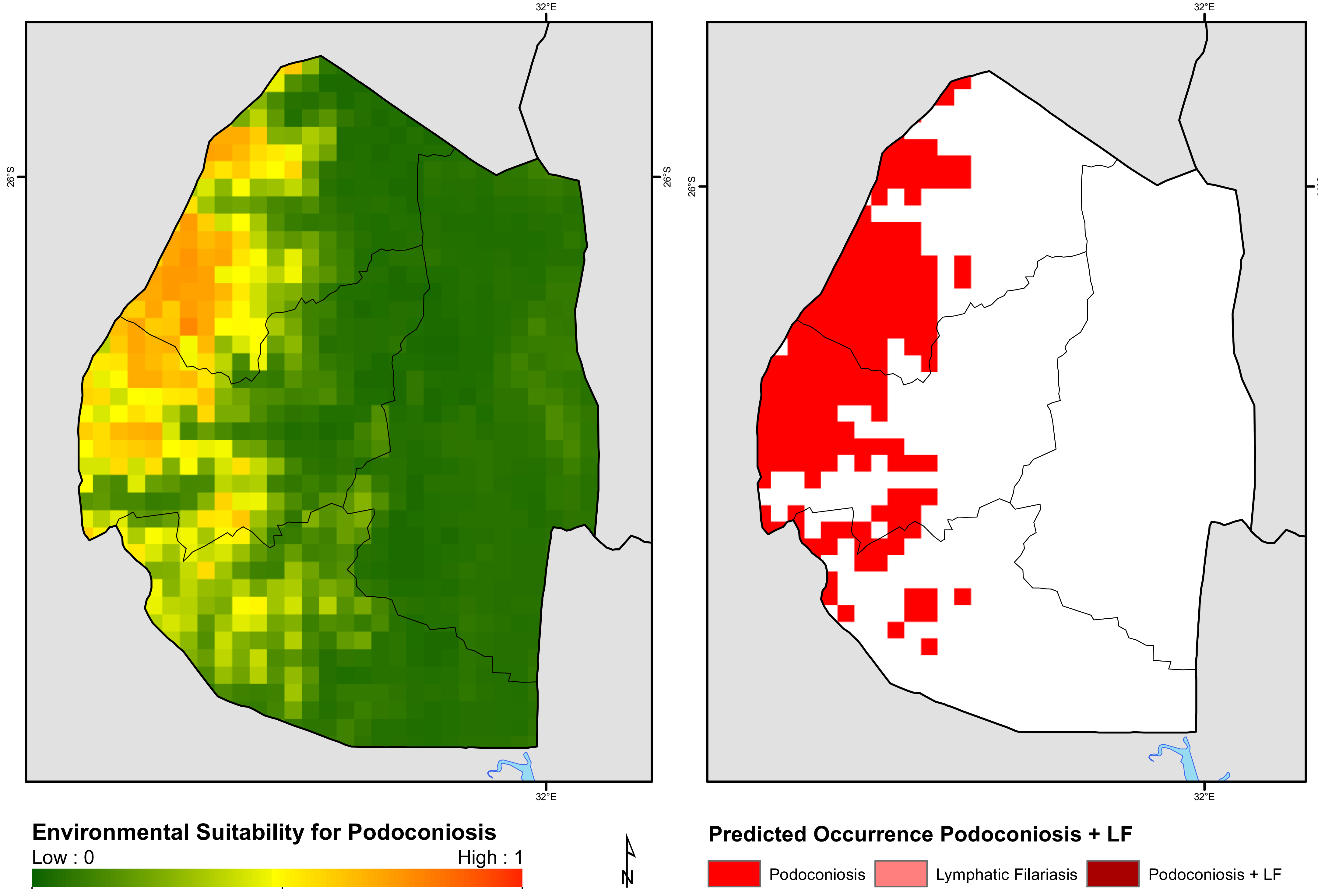
32°E
26°S
Environmental Suitability for Podoconiosis
Low : 0
High : 1
N
Predicted Occurrence Podoconiosis + LF
Podoconiosis
Lymphatic Filariasis
Podoconiosis + LF

### Togo

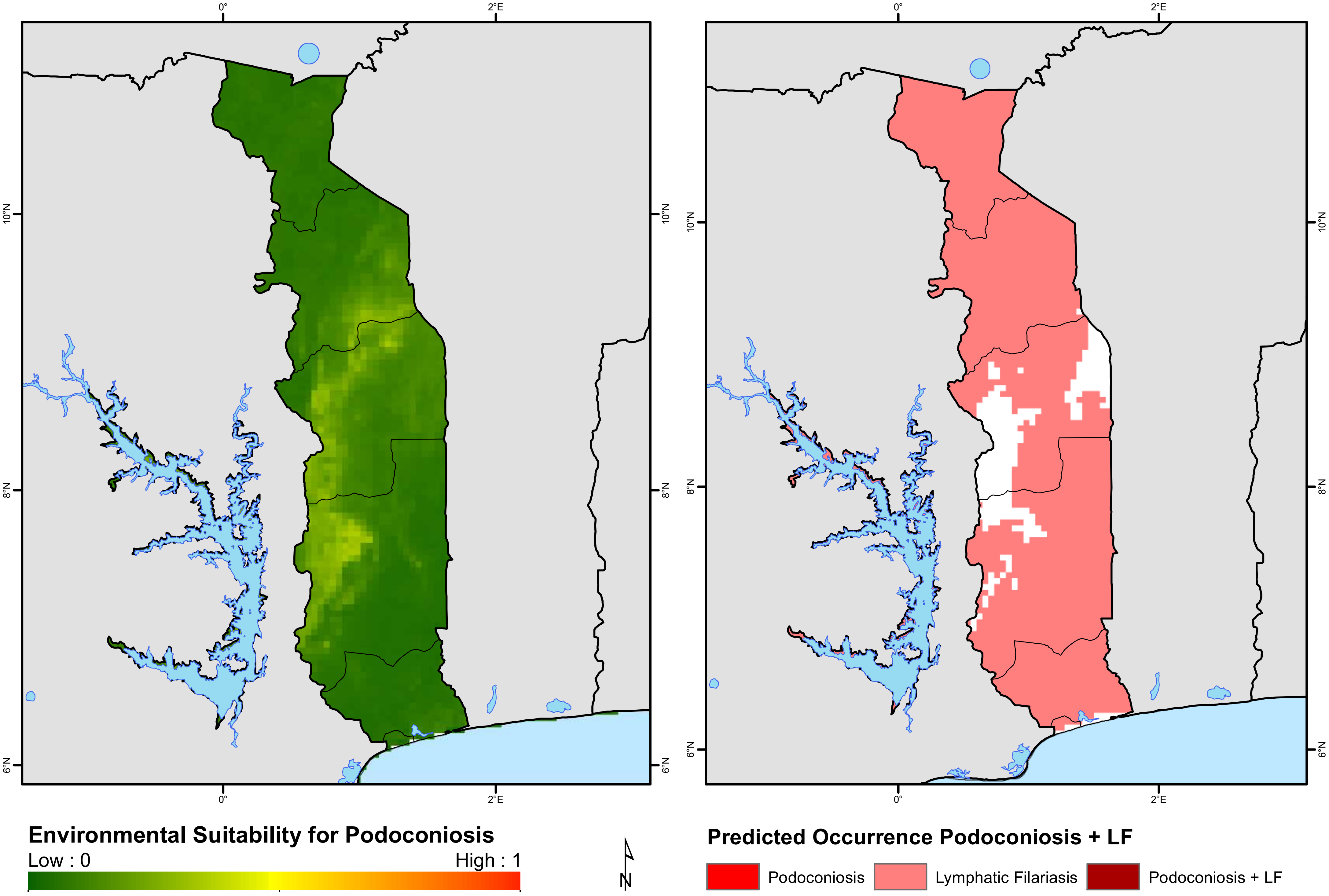
0°
2°E
10°N
8°N
6°N
Environmental Suitability for Podoconiosis
Low : 0
High : 1
N
Predicted Occurrence Podoconiosis + LF
Podoconiosis
Lymphatic Filariasis
Podoconiosis + LF

### Uganda

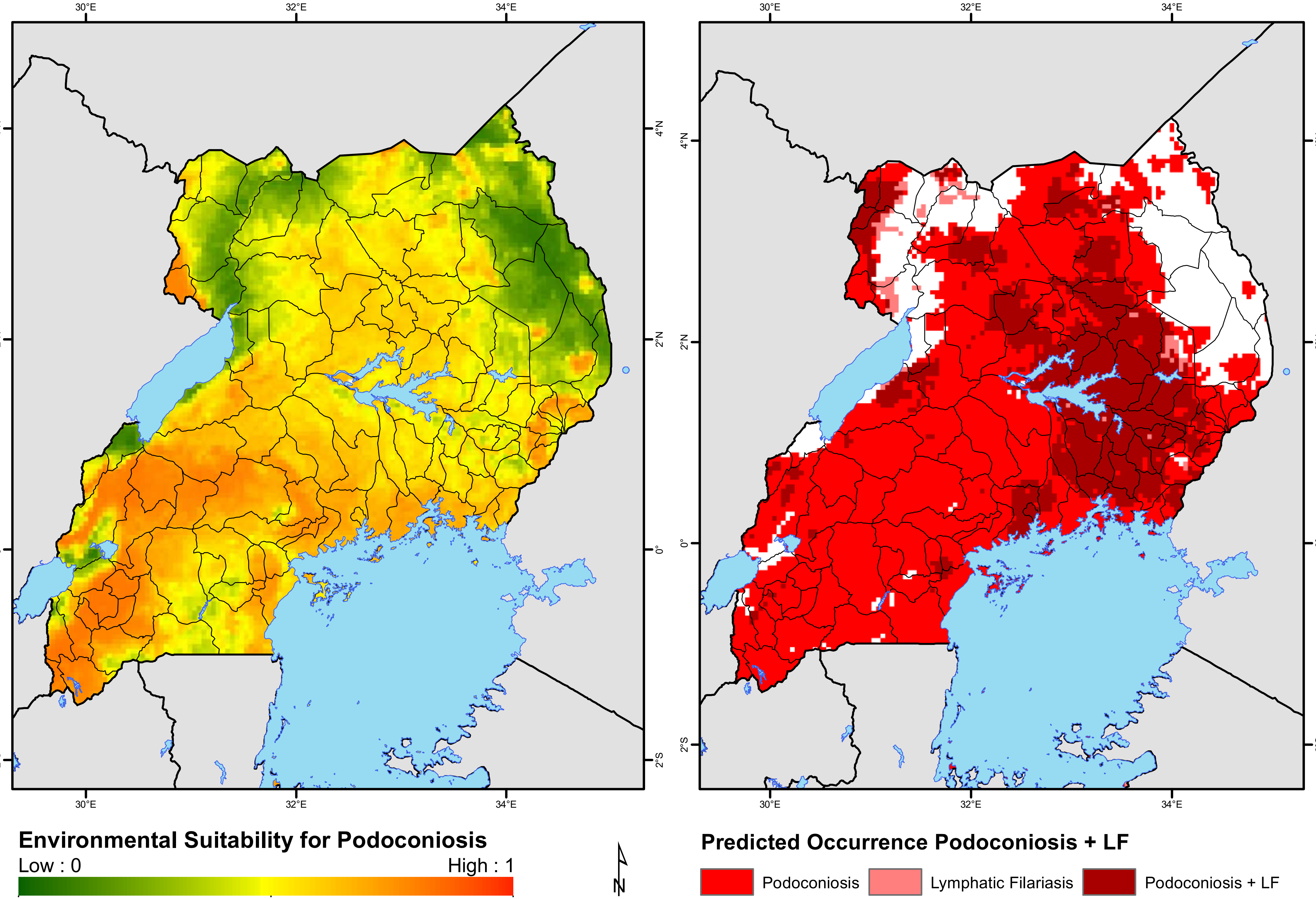
30°E
32°E
34°E
4°N
2°N
0°
2°S
Environmental Suitability for Podoconiosis
Low : 0
High : 1
N
Predicted Occurrence Podoconiosis + LF
Podoconiosis
Lymphatic Filariasis
Podoconiosis + LF

### United Republic of Tanzania

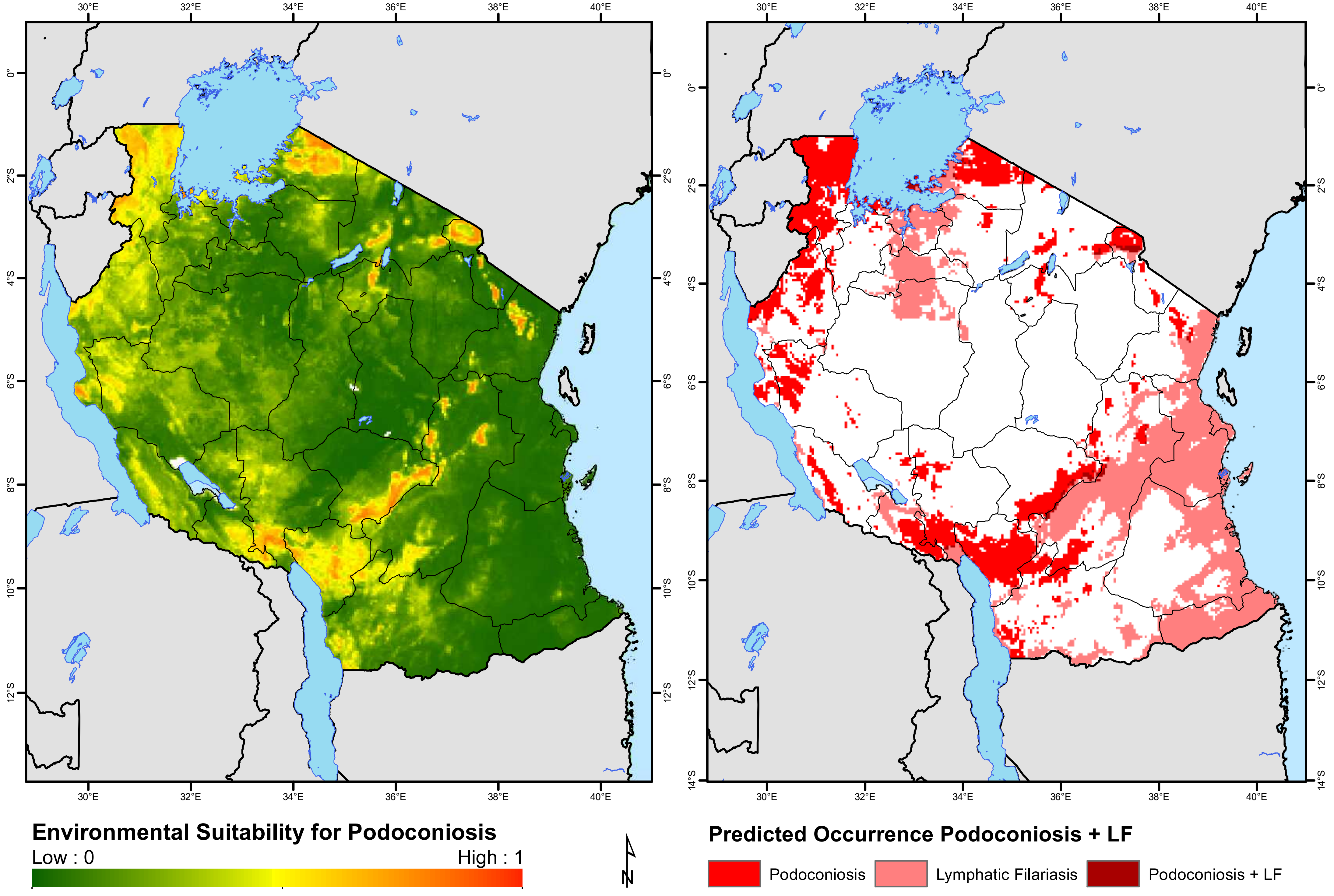

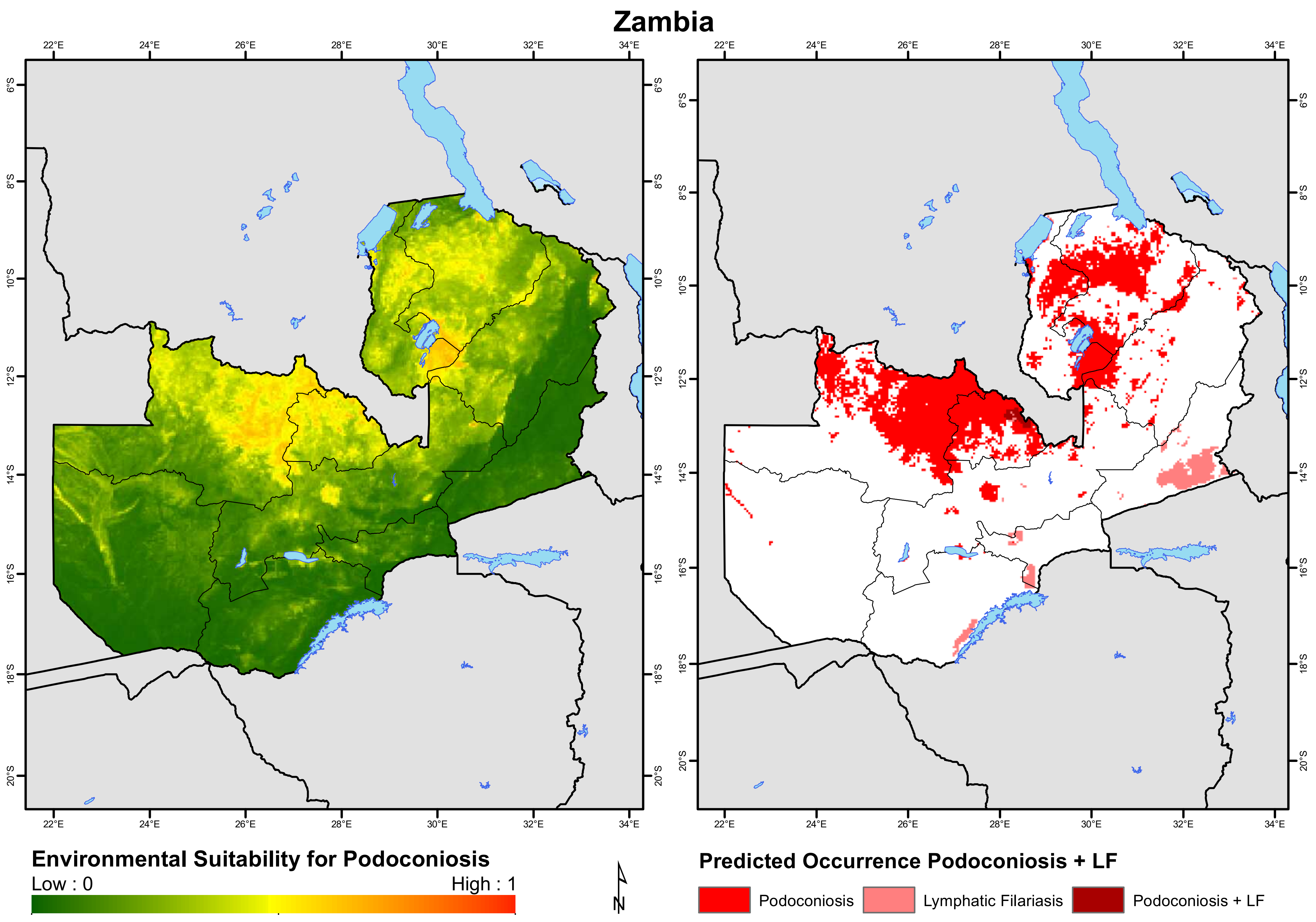
Zambia
22°E
24°E
26°E
28°E
30°E
32°E
34°E
6°S
8°S
10°S
12°S
14°S
16°S
18°S
20°S
Environmental Suitability for Podoconiosis
Low : 0
High : 1
N
Predicted Occurrence Podoconiosis + LF
Podoconiosis
Lymphatic Filariasis
Podoconiosis + LF

### Zimbabwe

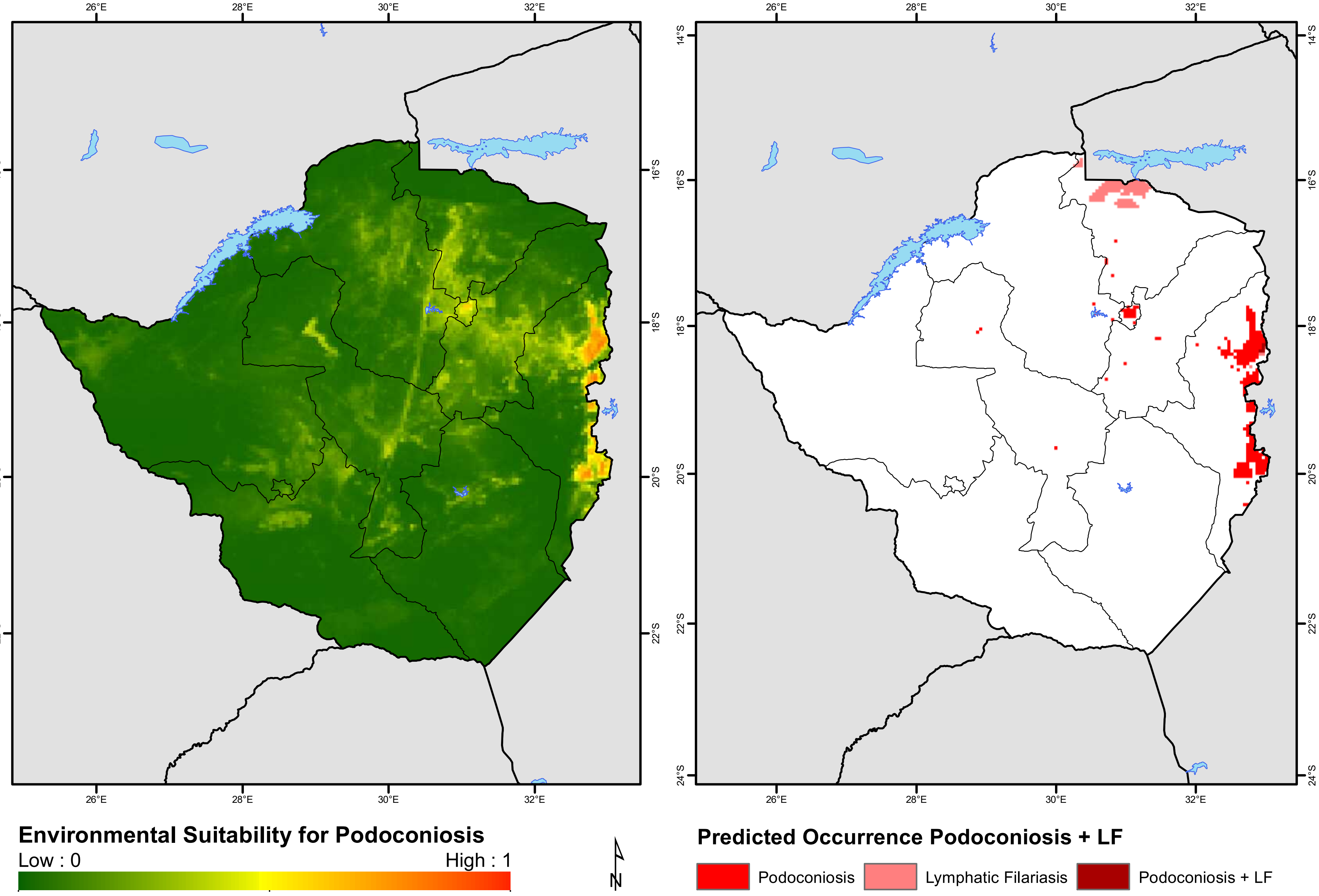

26°E
28°E
30°E
32°E
14°S
16°S
18°S
20°S
22°S
24°S
Environmental Suitability for Podoconiosis
Low : 0
High : 1
N
Predicted Occurrence Podoconiosis + LF
Podoconiosis
Lymphatic Filariasis
Podoconiosis + LF
